# A global synthesis on land-cover changes in watersheds shaping freshwater detrital food webs

**DOI:** 10.1101/2024.12.10.627188

**Authors:** Rebecca Oester, François Keck, Marcelo S. Moretti, Florian Altermatt, Andreas Bruder, Verónica Ferreira

## Abstract

Anthropogenic land-cover changes are among the most pressing global threats to both aquatic and terrestrial ecosystems, jeopardising biodiversity and the critical connections between these systems. Resource flows and trophic interactions intricately link aquatic and terrestrial ecosystems, with terrestrial-derived detritus playing a foundational role in supporting aquatic food webs. These detrital inputs form essential cross-ecosystem linkages, underpinning key ecological processes and providing vital resources for aquatic communities. Yet, little research has focused on how land-cover changes cascade across this linkage. To better understand how land-cover changes in the watershed influence freshwater detrital food webs, we conducted a meta-analysis of field studies reporting the effects of vegetation changes on freshwater detrital consumers and organic matter decomposition. The results from 144 studies, reporting 1235 comparisons, showed that, overall, land-cover changes in the watershed vegetation, especially through harvest and land-use conversion, have negative effects on aquatic biodiversity and ecosystem processes. These vegetation changes reduced diversity, abundance, and biomass across multiple trophic levels in freshwater detrital food webs. Studies examining multiple organism groups most often observed negative responses across multiple trophic levels, suggesting that the land-cover changes negatively affected multiple detrital food web components simultaneously. Our results also show that outcomes of restoration of watershed vegetation were context-dependent, and no clear trend of improvement was visible. Therefore, conservation of natural riparian and catchment vegetation are key to maintain freshwater ecosystem processes and aquatic biodiversity worldwide, and more efficient and evidence-based restoration measures are urgently needed. As our global synthesis shows that direct human-induced alterations of vegetation type in watersheds have significant negative effects on freshwater detrital food webs, there is a pressing need to consider cross-ecosystem consequences of land-cover changes in conservation and ecosystem management.

## Introduction

Human-driven land-cover changes represent a significant global threat to aquatic and terrestrial ecosystems, jeopardising biodiversity and the essential connections between these ecosystems (IPBES 2019). Land-cover changes that affect the vegetation composition, density or area in the watershed can have cascading effects on adjacent and downstream water bodies (England & Rosemond 2004; Hanna *et al*. 2020). These terrestrial impacts can significantly reduce freshwater biodiversity, negatively affect ecosystem processes, and disrupt cross- ecosystem linkages (Cereghetti & Altermatt 2023; Little & Altermatt 2018; Oester *et al*. 2023). In fact, one of the main direct drivers of aquatic biodiversity loss is land-use/land- cover change (Allan 2004; Harrison *et al*. 2018; IPBES 2019; Tickner *et al*. 2020; Zhang *et al*. 2023). In Europe, it has been estimated that 80 % of natural riparian vegetation has been lost during the last 200 years (Naiman *et al*. 1993). In most biomes, vast areas of riparian and catchment forests have already disappeared (Dudgeon 2000; Nunes *et al*. 2015; Warren *et al*. 2016) and the remaining fragments continue to degrade (Reid *et al*. 2019; Tolkkinen *et al*. 2020), especially in the tropics (Barnes *et al*. 2017; Dudgeon 2000). In different climatic regions, not only the trends and reasons for deforestations vary (Rudel *et al*. 2009), but also the composition of watershed vegetation (Kominoski *et al*. 2013), and aquatic biodiversity (Boyero *et al*. 2012). Vegetation changes such as harvesting trees for wood production, plantations of exotic timber species as well as logging or clear-cutting to converse watershed areas to agricultural land, can impact adjacent and downstream aquatic food webs in terms of abundance, but especially also of biodiversity and biomass of consumers and organic matter decomposition (Frainer & McKie 2021; Harding *et al*. 1998; Molinero & Pozo 2004; Silva- Araújo *et al*. 2020). For example, while forest harvesting can negatively affect macroinvertebrate biodiversity (Hernandez *et al*. 2005; Reid *et al*. 2010), plantations (e.g., with *Eucalyptus* spp.; Ferreira *et al*. 2019) have been shown to reduce leaf litter decomposition rates, and land-use conversion can change the metabolism of the entire ecosystem through shifts from heterotrophy to autotrophy (Hagen *et al*. 2010; Hebert *et al*. 2023). Also the scale of land-cover change (e.g., locally in the riparian zone or in the catchment) may differently affect streams as suggested for exotic eucalyptus plantations that have more severe negative impacts on leaf litter decomposition when present in both the catchment and the riparian zone than when only in the catchment (Ferreira *et al*. 2016). Due to the enormous scale and speed of ongoing land-cover changes, it is necessary to assess their consequences across ecosystem boundaries to “bend the curve of global freshwater biodiversity loss” (Tickner *et al*. 2020).

Aquatic and terrestrial ecosystems are closely linked through flows of resources and trophic interactions that occur at their interfaces (Baxter *et al*. 2005; Gounand *et al*. 2018). An important aquatic-terrestrial linkage is based on the influx, transportation, deposition and subsequent decomposition of organic matter from the riparian vegetation in adjacent and downstream water bodies (England & Rosemond 2004; Scherer-Lorenzen *et al*. 2022; Stoler & Relyea 2020). These cross-ecosystem resource flows not only influence the local communities (England & Rosemond 2004; Oester *et al*. 2023; Silva-Araújo *et al*. 2020) but also shape the energy pathways of the aquatic food web (Ho *et al*. 2023; Jacquet *et al*. 2022; Oester *et al*. 2024a). In many landscapes, the natural riparian and catchment vegetation around streams (especially around headwater streams) consists of native trees with species adapted to the local light and hydrological conditions (Tonello *et al*. 2021; Västilä & Järvelä 2018). The detritus of these trees, like leaves, branches or seed enters the water body either directly or after surface transport from the riparian zone (Riis *et al*. 2020; Scherer-Lorenzen *et al*. 2022).

When terrestrial detritus enters a water body, a suite of microbial decomposers, detritivorous macroinvertebrates, and other consumers feed on and break down this resource (Gessner *et al*. 2010). Microbes, such as aquatic fungi and heterotrophic bacteria, play crucial roles in decomposing leaf litter (Baldy *et al*. 1995; Gessner *et al*. 1999) through their own use and also by making it more palatable for higher-level consumers through nutrient immobilisation and maceration (García-Palacios *et al*. 2017). Detritivorous macroinvertebrates, such as the functional feeding group of shredders (*sensu* Moog 1995), preferentially feed on microbially conditioned detritus (Arsuffi & Suberkropp 1985; Graça & Cressa 2010) and further fragment this material and make it available to a variety of other aquatic organisms (Graça 2001). However, not only specialised shredders but also omnivores, like other macroinvertebrates, fish, and crayfish, opportunistically use terrestrial detritus as resource (Allen *et al*. 2024; Dudley *et al*. 2021; Evangelista *et al*. 2014). Multitrophic biodiversity in the freshwater detrital food web can accelerate the decomposition of detritus and other ecosystem processes (Gessner *et al*. 2010; Handa *et al*. 2014; Oester *et al*. 2024b). Decomposition is an essential ecosystem function that integrates the feeding activity and trophic relationships in the detrital food web and responds sensitively to anthropogenic change (Gessner *et al*. 2010; Hebert *et al*. 2023). Hence, decomposition rates could provide a complementary index for ecosystem assessments (Frainer *et al*. 2021; Lopes *et al*. 2015). To fully understand the freshwater food-web dynamics and their responses to environmental change, it is essential to assess the components of the detrital food web—both terrestrial resources and aquatic consumers—and the decomposition processes that link them.

As the watershed vegetation not only mediates abiotic conditions (e.g., light, nutrients, erosion, temperature), but also terrestrial matter input and habitat provision (Ferreira *et al*. 2023; Richardson & Béraud 2014; Riis *et al*. 2020; Tolkkinen *et al*. 2020), land-cover changes can affect the adjacent stream biodiversity and ecosystem processes through multiple interactive pathways (Maloney & Weller 2011). The direction and magnitude of land-cover effects on aquatic food web dynamics can be context-dependent (Ferreira *et al*. 2019; Richardson & Béraud 2014) and can vary, for example, across climate regions (GarcíaLPalacios *et al*. 2016; Handa *et al*. 2014) or taxa (Martínez *et al*. 2013; Truchy *et al*. 2022). Likely because of the complicated interplay of direct and indirect effects, there is so far no clear overview or consensus on the relationship between land-cover changes in the watershed vegetation and freshwater detrital food web composition and processes.

Here we present a systematic meta-analysis of 144 field studies addressing the effects of direct human-induced watershed vegetation change (i.e., restoration, plantation, harvest, and land-use conversion) on freshwater detrital food webs published between 1993 and 2023. Using a systematic appraoch, we (i) determined the significance, magnitude and direction of the mean effect of vegetation change on freshwater detrital communities and organic matter decomposition, (ii) assessed which moderators (i.e., explanatory variables) influence the magnitude and direction of the overall effects and the effects for each trophic level separately, and (iii) assessed if studies observe similar directions of effect sizes across trophic levels. We expected to find (i) an overall negative effect of vegetation change on freshwater detrital food webs, (ii) especially in the tropics and subtropics (as the climatic regions undergoing currently the most drastic land-cover changes), where the watershed vegetation was completely converted to other land-use types, where watershed vegetation is entirely (i.e., locally and in the catchment) altered, for diversity and biomass metrics and, for microbe and shredders (as the most detritus-dependent groups), and (iii) that these effects are simultaneously detectable at multiple trophic levels of freshwater detrital food webs (Barnes *et al*. 2017; Burwood *et al*. 2021; Casotti *et al*. 2015; Cornejo *et al*. 2020).

## Material and Methods

### Literature search and study selection

We conducted a systematic literature search using the ISI Web of Science (WoS) Core Collection (Edition: Science Citation Index Expanded) to identify records examining the consequences of direct human alterations in land cover in the riparian zone and catchment on elements of freshwater detrital food webs (i.e., detritus, microbial decomposers, shredders, and omnivores; Figure 1A). Specifically, we searched for empirical studies comparing community metrics (i.e., α-diversity, abundance, biomass) or organic matter decomposition in freshwater detrital food webs with a before-after or a control-impact design. Reference conditions are defined as near-pristine or natural watershed vegetation such as native riparian forests, whereas direct human-induced alterations in the vegetation conditions could, for example, consist of plantations, vegetation restoration activities, land conversion (e.g., from forest to pasture) or forest harvest, but not damages caused by storms, wildfires, or exotic species. We also *a priori* excluded comparisons between reference and urban sites, as the vegetation in urban sites is often replaced by impervious surfaces (Maloney & Weller 2011).

**Figure 1:**
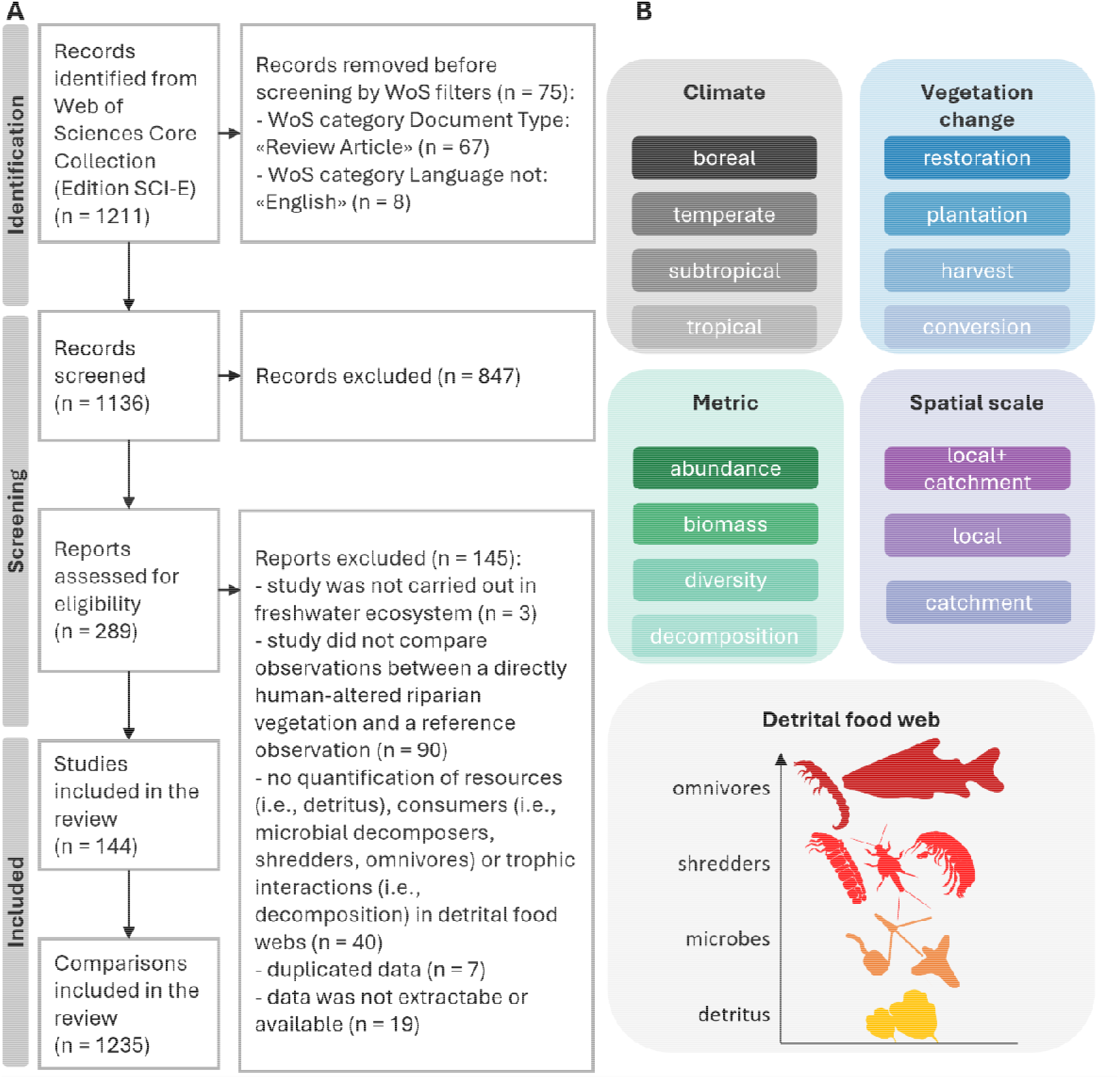
Overview of methods used in this review. A) PRISMA flow diagram with the number of studies located in the literature search and study selection and B) Main moderators extracted for the assessment of the effects of vegetation change on freshwater detrital food webs.

We used the following search string in the search field “topic”, which searches title, abstract and keywords: (stream OR river OR lake OR pond OR freshwater) AND ("land use" OR "land cover" OR deforestation OR logging OR fragmentation OR "forest composition" OR plantation OR forestry OR restoration OR conservation OR revitalization OR revitalisation OR clear OR harvest OR removal) AND (riparian OR forest OR buffer) AND (decomposition OR processing OR breakdown OR decay OR brown) AND ("food web” OR trophic OR detritivor OR shredder OR invertebrate OR decomposer OR hyphomycete OR fung OR bacteri OR microb). We searched for records published before 16 April 2024, which yielded 1211 records. We then applied the WoS filters of Document Type to exclude reviews (n = 67) and Non-English Records (n = 8; Figure 1A).

Next, we screened titles and abstracts of 1136 records to determine whether the record had potential to contain suitable data, using the following criteria: 1) the study was carried out in a freshwater ecosystem; 2) the study compared field observations and experimental outcomes between a directly human-altered watershed vegetation and a reference condition; 3) the study quantified resources (i.e., detritus), detritus-consuming communities (i.e., microbial decomposers, shredders, omnivores at least partially consuming detritus) or trophic interactions (i.e., decomposition). We excluded records that did not meet these three criteria (n = 847). In a next step, we read the full text of the 289 remaining records and excluded 145 additional records that did not fulfil the three above-mentioned criteria. We also excluded records that reported duplicate data (n = 7) or where data was not extractable or available (n = 19). The final set of records included 144 studies. For further details on the identification and screening process according to the Preferred Reporting Items for Systematic reviews and Meta-Analyses (PRISMA; O’Dea *et al*. 2021; Page *et al*. 2021) see Figure 1A.

### Extraction of primary data

A single study could report several comparisons between reference and human-altered conditions if using, for example, different litter species (e.g., Cornejo *et al*. 2020; Masese *et al*. 2014), consumer groups (e.g., Encalada *et al*. 2010), community metrics (e.g., Oester *et al*. 2023), or regions (e.g., Ferreira *et al*. 2019; Riipinen *et al*. 2009), resulting in a total of 1235 comparisons (Figure 1A). We did not extract data which were not unambiguously comparable. These cases include: 1) different levels of the same type of vegetation change across a gradient (e.g., “low”, “medium” and “high” logging intensity), in which case we compared the two extreme ends of the gradient; 2) different taxonomic levels (e.g., family, genus, species) of the same community, in which case we took the finest available level (i.e., species if possible); 3) different calculations for similar diversity metrics (e.g., Shannon index, richness, evenness), in which cases we extracted richness if provided, or different decomposition quantifications (e.g., % mass loss, decomposition rates), in which cases we extracted decomposition rates in degree days if provided. We recorded the mean, variation (i.e., standard deviation (SD), standard error (SE) or confidence interval (CI)), and number of replications for each metric in the reference and altered condition being compared.

We extracted data from text, tables, figures, or requested them from authors. We used WebPlotDigitizer v.5.0 (https://automeris.io/WebPlotDigitizer/), in manual mode, to extract values from figures, which corresponded to 37.8 % of all comparisons. If variation in the primary studies was reported or provided by the authors as SE or CI, we converted it into SD. However, no measure of variation was available in 7.7 % of cases, and we estimated missing SD values by imputation based on the cases in the dataset that reported SD values associated with the same metric and trophic level (Lajeunesse 2013). Extracting data from figures and imputing missing variation values can potentially introduce a bias in the dataset. Consequently, we assessed the possible bias introduced by these potentially low-quality data through sensitivity analyses (see below).

### Extraction of moderators

Several biotic and abiotic explanatory variables (i.e., moderators), may affect the magnitude and direction of the response of elements of freshwater detrital food webs to human-induced changes in the watershed vegetation. Based on our hypotheses, we included five general moderators (Figure 1B, Table S1): climate (Ferreira *et al*. 2019), type of vegetation change (Goodman *et al*. 2006; Inoue *et al*. 2012; Jinggut *et al*. 2012), spatial scale of vegetation change (Ferreira *et al*. 2016), community or decomposition metric (Iñiguez- Armijos *et al*. 2018; Oester *et al*. 2023), and trophic level (Hebert *et al*. 2023). We also recorded type of water body (i.e., lake, pond, river, streams, bromeliads), but as more than 99 % of all comparisons were based on streams (Table S1), we did not consider this moderator for further analyses.

We extracted watershed vegetation type of reference and altered sites as well as the main human-induced change in the altered sites. We later coded the different types of changes into the following four categories: restoration, plantation, harvest, conversion (Figure 1B, Tables S1). For restoration activities, we used the restored site as the reference condition, which allowed us to align it with other types of changes in watershed vegetation. By reversing this comparison, we interpreted the direction of change as indicative of what would have happened without restoration measures. Land-use conversions mostly included changes from a native forest to pasture, grassland or other agricultural land uses.

For each study, we also collected the location (latitude and longitude) and climate (i.e., boreal, temperate, subtropical, tropical; Figure 1B, Table S1). If the exact latitude and longitude were not reported, we extracted locality information (e.g., country, state, catchment) from manuscript figures, tables, text or requested it from authors. We also extracted the spatial scale of vegetation change either including both the local riparian vegetation and the catchment (local+catchment), only the local riparian vegetation (local) or only the catchment (catchment) according to the information in the publication or provided by authors (Figure 1B, Table S1).

To quantify the effects of vegetation change on the freshwater detrital food web, we extracted the metric of the food web element assessed (i.e. α-diversity, abundance, biomass, decomposition) and the trophic group (i.e., detritus, microbes, shredders, omnivores; Figure 1B, Table S1). In case where changes in detritus were assessed, we extracted the organic matter type (i.e., leaf, wood, cotton strip, reproductive material, or mixed matter) and collected the taxon name if available and applicable. If both single and mixed species treatments were assessed, we included only single species treatments to avoid mixing effects. We extracted the broad taxonomic group for microbial communities (i.e., microbe, bacteria, fungi), and for secondary consumer communities (i.e., shredders, macroinvertebrates, fish+crayfish). We also extracted the microhabitat where the communities were collected (i.e., detritus, benthic, water column). The three community metrics could be measured at all trophic levels, while the decomposition metrics (i.e., microbial decomposition, shredder- mediated decomposition, total decomposition) were assigned to detritus only.

### Effect size

We calculated the effect size of human-induced vegetation change on freshwater detrital food webs as standardised mean difference “Hedges’ g”: (m1_i_−m2_i_)/pSD_i_, where the mean difference between altered (m1) and reference (m2) conditions is divided by the pooled standard deviation (pSD; Hedges 1981). Hedges’ g provides a standardised effect size that accounts for variability within the groups and corrects for bias in small samples, making it useful for comparing studies with different scales or measures (Hedges 1981). Effect sizes of 0 indicate no difference between altered and reference conditions, while negative or positive effect sizes indicate that the altered condition shows lower or higher values than the reference condition, respectively. Absolute effect sizes of around 0.2 indicate small effects, 0.5 indicate moderate effects and 0.8 or higher indicate large effects (Cohen 1988).

### Data analyses

We used mixed-effect models in R v. 4.4.1 (R Core Team 2024) to test the effect of different climate, type and scale of vegetation change, trophic level and metrics on freshwater detrital food webs. To account for heterogeneity within and between studies, we constructed linear random/mixed-effects models, using the variances of each mean as weighing factors and study identity as random effect (intercept). These models were built using the default settings of the restricted maximum likelihood method in the metafor package (Viechtbauer 2024). Accordingly, we used this type of models to assess the overall mean using the entire dataset and including moderators as fixed effects. We included all the moderators separately with only one moderator at once to provide an overview of the individual effects of each moderator.

Based on the high heterogeneity level of the dataset, we then split the dataset according to trophic level (i.e., detritus, microbes, shredders, omnivores) to analyse the interactive effects of moderators and component of the detrital food web separately. For each trophic level, we tested the effects of climate, type of vegetation change, spatial scale of vegetation change, and community metric.

We present results as effect size plots using ggplot2 (Wickham et al. 2024). Negative and positive effect sizes indicate lower and higher values in the altered compared to the reference conditions, respectively, with significant effects occurring when 95 % CIs do not overlap 0. We calculated the percentage of total variation explained by between-study variation (I^2^) for the grand mean and separate outcomes for each moderator level for our mixed-effect models according to Jackson *et al*. (2012).

Lastly, to assess whether changes in the watershed vegetation influenced multiple trophic levels simultaneously, we analysed effect sizes of studies with comparisons of at least two trophic levels. We calculated linear regressions and “Spearman” rank correlations using a simple linear model with the median effect size of one trophic level against another trophic level for each study. We present these results as quadrant graphs showing studies that provided data that fall within either quadrant of: -/-, +/-, -/+ or +/+ effect size space, with negative and positive values indicating negative and positive effects of change in the vegetation on the median effect size value of a trophic level, respectively.

### Publication bias and sensitivity analyses

We assessed publication bias by visually inspecting funnel plots for potential asymmetry driven by small-studies (Møller & Jennions 2001). Additionally, we performed a File Drawer Analysis (Rosenberg 2005) to further confirm the robustness of our findings against selective publication. Rosenberg’s fail-safe number (N_fs_) provides the number of missing effect sizes showing a non-significant effect that would be needed to nullify the mean effect size, with N_fs_ > 5 × n + 10 (n = number of effect sizes) indicating that the dataset can be considered robust to publication bias. Additionally, we applied a newly developed two-step approach by Yang *et al*. (2024) for mixed-effect models to assess publication bias and non- independence using the orchard (Nakagawa *et al*. 2023) and clubSandwich packages (Pustejovsky 2024). The results of these analyses (Table S2, Figures S1–S5) suggest that our findings are robust and not unduly influenced by publication bias. Only the subset of data regarding effects on omnivores was not robust to publication bias (N_fs_ = 1936, n = 389). In fact, the bias robust estimate was in the opposite direction to that found based on the dataset (Figure S5), which indicates missing effect sizes to the right of the mean effect size possibly due to the lack of unpublished studies with significantly positive results.

To assess the sensitivity of the results to the inclusion of estimated and imputed data, we repeated the analyses including only reported comparisons. Reasons for concern about the inclusion of data of potentially less quality (i.e., estimated and imputed data) in the analyses would occur if the interpretations of results would differ when considering all data or only reported comparisons. We report the sensitivity analysis in the Supplement (Figures S6, S7, Tables S3, S4).

## Results

### Overview of studies

The 144 studies comprising 1235 comparisons covered a broad range of latitudes (- 45.29 to 65.57; Figure 2A), types of changes in the watershed vegetation (Figure 2B) and spatial scale of these changes (Figure 2C), community metrics and decomposition (Figure 2D), and trophic levels (Figure 2E). The studies spanned a 30-year period (1993–2023; Figure S8), with on average 8.6 comparisons per study (Figure S9).

**Figure 2:**
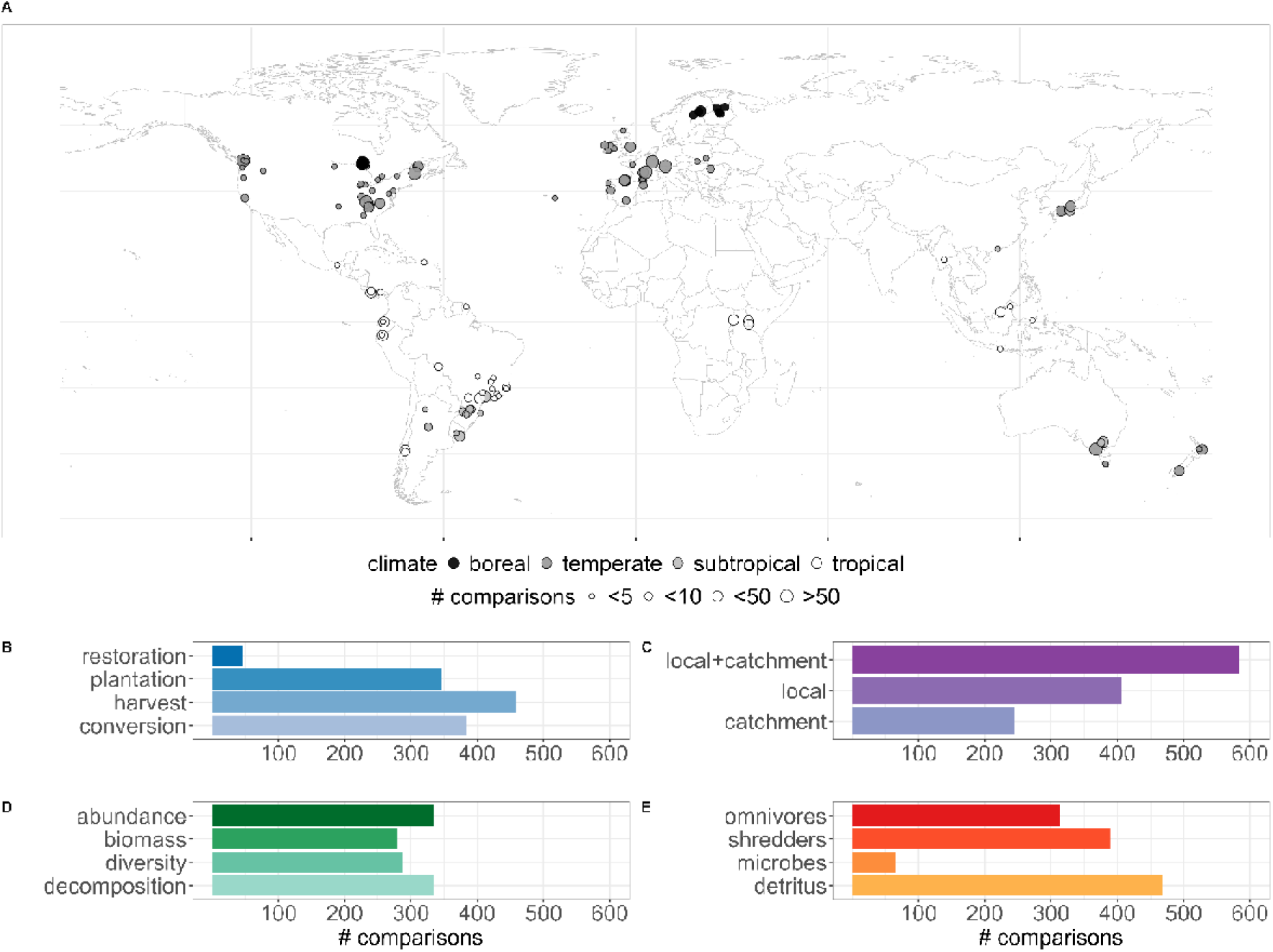
Geographic distribution of studies included in this review and histograms of data distribution of different moderators. A) World map showing the study locations with each circle representing a study. Panels B – E represent histograms for each level in the moderators: B) type of vegetation change, C) scale of vegetation change, D) metric and E) trophic level.

Most studies (n = 80) and comparisons (n = 753) were conducted in temperate zones, followed by studies from the tropical (38 studies), subtropical (16) and boreal zones (10), and only one study reported comparisons of multiple climate zones (Ferreira *et al*. 2019). We found large spatial gaps in data coverage in Africa and Asia (Figure 2A).

Most commonly assessed vegetation changes in the temperate zone were plantations (n = 313 comparisons), harvest (n = 294) and conversion (n = 117), while in the tropics and subtropics it was conversion (n = 197 and n = 70, respectively), and in the boreal zone it was harvest (n=74; Figure S10). While most comparisons reflected changes at local and catchment scales simultaneously (n = 583), we also found 407 comparisons at local and 245 comparisons at catchment scales (Figure 2C).

Most studies assessed one trophic level and two metrics. Most common trophic level and metric combinations were detritus decomposition (n = 334 comparisons), omnivore abundance (n = 175) and diversity (n = 159), and shredder abundance (n = 151).

### Overall effects of changes in the watershed vegetation on freshwater detrital food webs

The majority (58 %) of individual effect sizes were negative with a large portion (37 %) being moderately to strongly negative (effect size < -0.5), which resulted in a grand mean effects size of -0.29 (95 % CI = [-0.41, -0.17]; Figure 3A), indicating a significant reduction in the components and processes in the freshwater detrital food web as a consequence of direct human-induced changes in the vegetation. Even after correction for publication bias, the bias-robust estimate remained significantly negative (-0.06 [-0.11, -0.01]; Figure S1). There was, nevertheless, high between-study heterogeneity contributing to total heterogeneity (I^2^ = 93.83).

**Figure 3:**
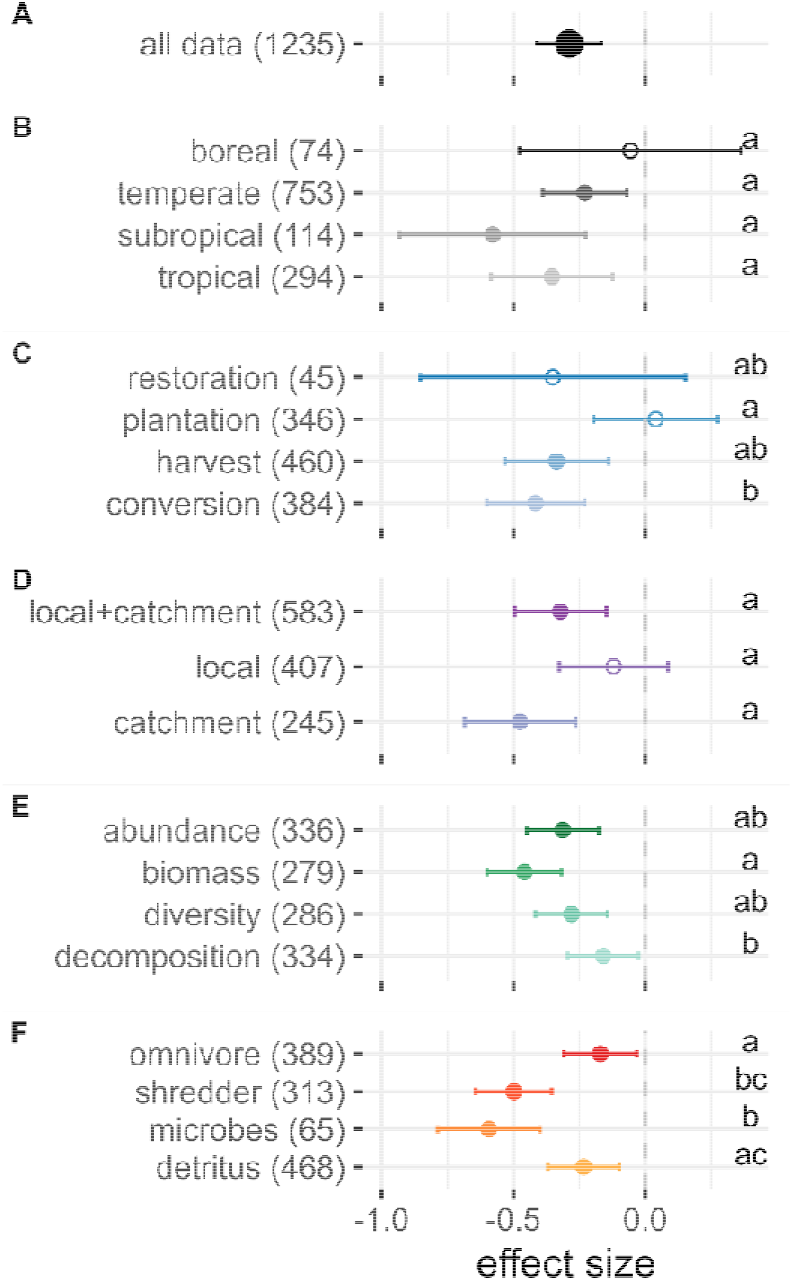
Impacts of human-induced alterations in the watershed vegetation on freshwater detrital food webs. The global response (all data) is shown on the first row A) and is separated by moderators in the following rows for B) climate, C) type of vegetation change, D) spatial scale, E) metric and F) trophic level. The numbers in brackets after each moderator level represent the number of comparisons. For each moderator level, the circle represents the mean effect size with 95 % confidence intervals (CIs) computed from the random effects model. Filled circles indicate statistically significant effects, whereas empty circles have CIs that cross the 0-line and are thus statistically non-significant. Negative effects sizes (standardised mean differences Hedges’ g between altered and reference vegetation conditions < 0) indicate that the values in the altered conditions were lower compared to the reference conditions. Groups sharing the same letter (e.g., ’a’, ’ab’, ’b’) are not significantly different from each other as their CIs overlap.

### Effects of main moderators

The magnitude of the effect of human-induced vegetation change on freshwater detrital food webs depended not on climate but on type of vegetation change, spatial scale, trophic level and community metric (Figure 3, Table 1). Although the effects of vegetation change on freshwater detrital food webs did not significantly differ among climate zones (Table 1), they were significantly negative for temperate, subtropical and tropical climates and non-significant for boreal climate (Figure 3B). Whitin types of vegetation change, effects were significantly stronger for land-use conversions than for plantations (Figure.3C). While there were overlaps in the CIs of responses to different spatial scales (Figure 3D), spatial scale of vegetation change was still a significant moderator for mean land-cover effects on the freshwater detrital food web (Table 1). Within community metrics, effects were significantly stronger for biomass than for decomposition (Figure 3E). Within trophic levels, effects were significantly stronger for shredders than for omnivores and for microbes than for omnivores and detritus (Figure 3F).

**Table 1.** Effects of moderators tested in the analyses based on the entire dataset (n = 1235 comparisons from 144 studies). Test for heterogeneity between levels within moderators (Q_M_), degrees of freedom (df) and p-values (significant differences among moderator levels exist if p-values < 0.05; see Figure 3) are shown. Rosenberg’s fail-safe number (N_fs_) was 6892, indicating the robustness (N_fs_ > 5 × n +10, n = 1235) of our dataset to publication bias.

| Moderator | Levels | $Q_M$ | df | p-value |
| --- | --- | --- | --- | --- |
| All data | | 4225.18 | 1234 | $< 0.01$ |
| Climate | 4 levels: boreal, temperate, subtropical, tropical | 4.62 | 3 | 0.20 |
| Type of vegetation change | 4 levels: restoration, plantation, harvest, conversion | 10.31 | 3 | 0.02 |
| Spatial scale | 3 levels: local+catchment, local, catchment | 6.50 | 2 | 0.04 |
| Metric | 4 levels: abundance, biomass, diversity, decomposition | 32.97 | 3 | $< 0.01$ |
| Trophic level | 4 levels: omnivores, shredders, microbes, detritus | 64.63 | 3 | $< 0.01$ |

### Effects of moderators within trophic levels

To test for interactions between trophic levels and other moderators, we compared the effects of moderators on different trophic levels and found that trophic levels differently responded to climate, type of vegetation change, scale of vegetation change, and metric observed (Figure 4, Tables 2, S2).

**Figure 4:**
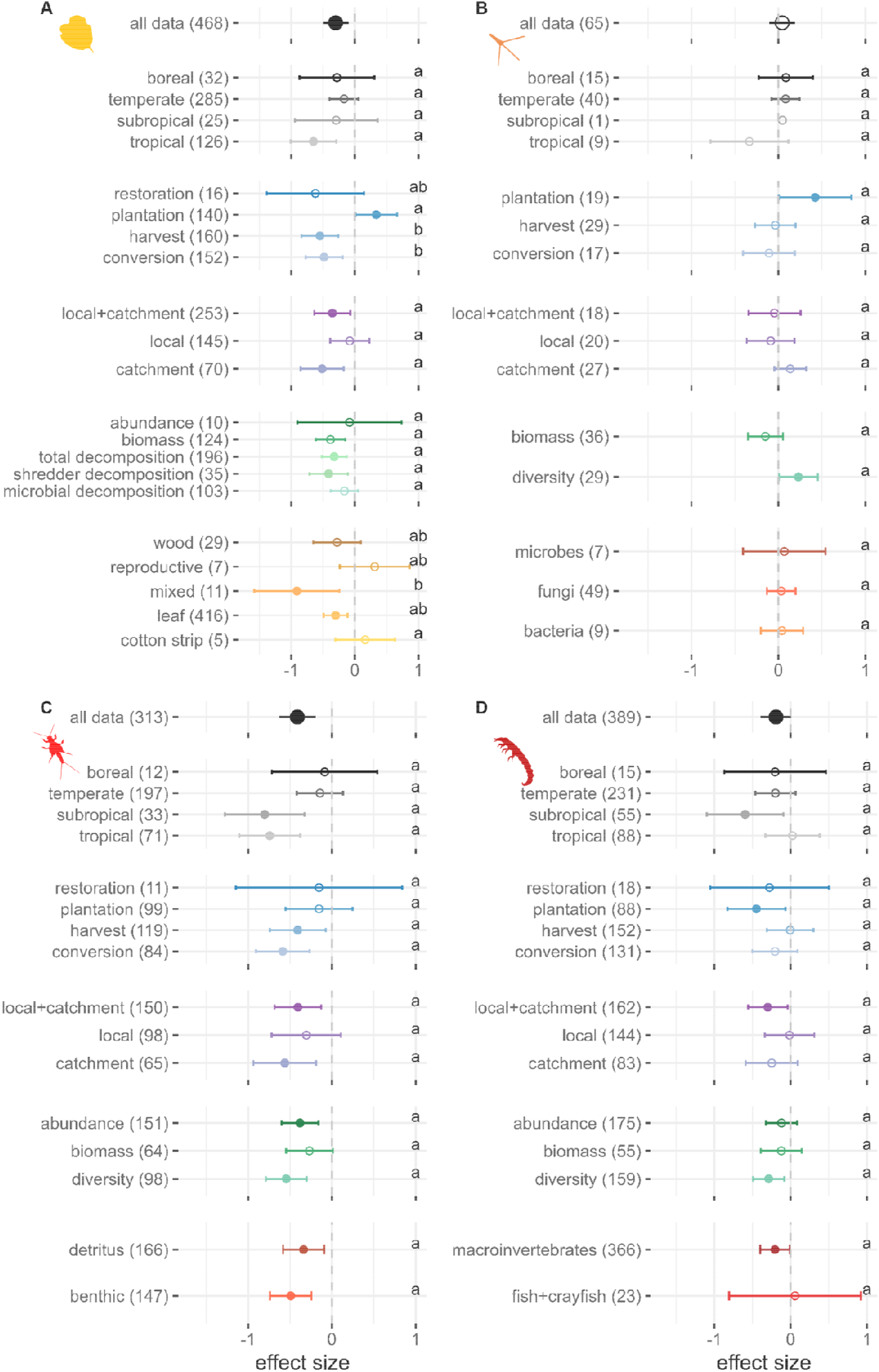
Moderator analysis separated for the subsets of each trophic level with A) detritus, B) microbes, C) shredders and D) omnivores. For each moderator level, the circle represents the mean effect size with 95 % confidence intervals (CIs) computed from the random effects model. Filled circles indicate statistically significant effects, whereas empty circles have CIs that cross the 0-line and are thus statistically non-significant. Negative effects sizes (standardised mean differences Hedges’ g between altered and reference vegetation conditions < 0) indicate that the values in the altered conditions were lower compared to the reference conditions. Groups sharing the same letter (e.g., ’a’, ’ab’, ’b’) are not significantly different from each other as their CIs overlap.

For detritus, type of vegetation change was the most influential moderator with most negative effects of harvest and land-use conversion, non-significant effects of restoration and positive effects of plantation (Figure 4A, Tables 2, S2). While type of detritus was an important moderator, climate zones, spatial scale and metric did not influence the effect of vegetation change on detritus. However, we also found statistically significant negative effects in tropical sites, where vegetation changes at catchment and local+catchment scales, and in terms of biomass. For decomposition of detritus, we found significant negative responses for total and shredder-mediated decomposition but not or microbial decomposition.

For microbes, community metric was a statistically significant moderator with positive effects of vegetation change on diversity and non-significant negative effects on biomass (Figure 4B, Tables 2, S2). The other moderators for this trophic level were not significant, with the only other statistically significant effect in plantations.

For shredders, climate mostly moderated the effect of vegetation change with significantly negative effects in the tropics and subtropics and non-significant negative effects in boreal and temperate zones (Figure 4C, Tables 2, S2). While type of vegetation change was not a statistically significant moderator, we found significant negative effect sizes for harvest and land-use conversion. Spatial scale of vegetation change was also not a statistically significant moderator, but effects were significantly negative at catchment and local+catchment scales but not at local scales. Effects of vegetation change also did not significantly differ among metrics, although effects were significantly negative for abundance and diversity but not biomass.

For omnivores, none of the moderators was statistically significant. However, we found negative effects of vegetation change on omnivores in the subtropics (Figure 4D, Tables 2, S2), when the vegetation changes included plantations, when the spatial scale encompassed local+catchment, in terms of diversity, and when macroinvertebrates were considered.

### Multitrophic effects

Most studies showed simultaneous negative effects for different combinations of trophic levels (Figure 5). For detritus together with microbes (Figure 5A), shredders (Figure 5B) and omnivores (Figure 5C), we found that 43 %, 55 % and 44 % of all studies showed negative effects sizes for both trophic levels, respectively. Also for microbes together with shredders (Figure 5D) and omnivores (Figure 5E), there were 40 % and 22 % of studies showing simultaneous negative effect sizes, respectively. Lastly, for shredders together with omnivores, we found 44 % of all studies showing negative effect sizes (Figure 5F). For studies, simultaneously assessing detritus together with microbes, and shredders together with omnivores, we found statistically significant regressions and correlations (detritus-microbes: intercept = -0.18, slope = 0.45, SE = 0.16, rho = 0.66, p-value < 0.01; shredders-omnivores: intercept = 0.23, slope = 0.90, SE = 0.15, rho = 0.48, p-value < 0.001).

**Figure 5:**
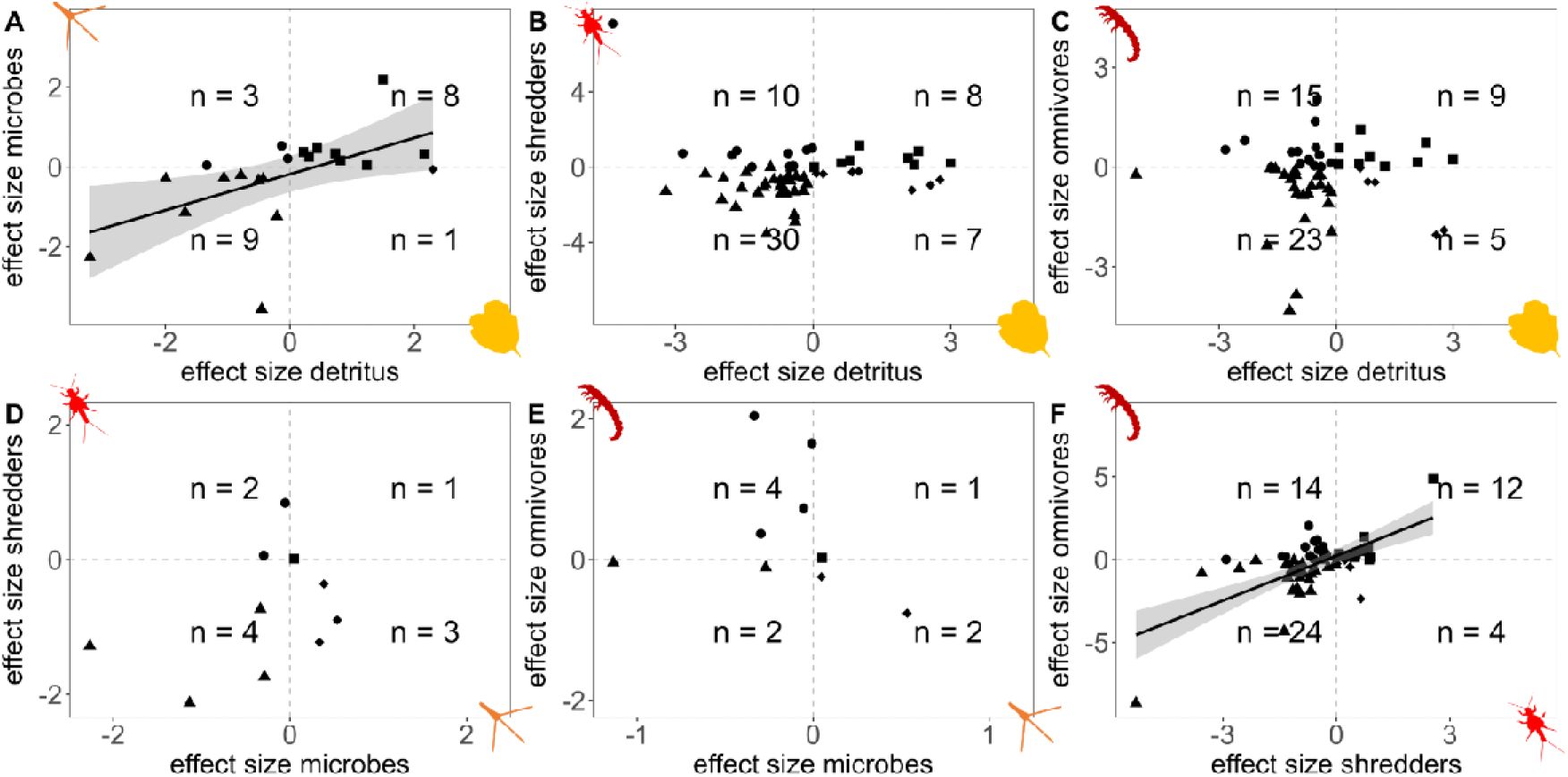
Quadrant graphs of median effect sizes of one trophic level against another in all possible combinations. Each point represents a study, and the symbols indicate the quadrant in which the effect sizes of the study lie in the x/y-axis (triangles (-/-), circles (-/+), diamonds (+/-) and squares (+/+)). The quadrants are separated by a dashed grey line going through 0 on both axes. Negative effects sizes (standardised mean differences Hedges’ g between altered and reference vegetation conditions < 0) indicate that the values in the altered conditions were lower compared to the reference conditions. For panels A) and F) we found statistically significant positive relationships indicated by the regression with the grey shaded area representing the confidence area around the linear regression.

### Sensitivity analyses

Repeating the analyses without estimated values (i.e., without values extracted from figures or imputed values) did not change our results and interpretations much (Figures S6, S7, Tables S3, S4). The overall response to land-cover changes in the watershed on freshwater detrital food webs remained significantly negative (Figures S1, S6). Only considering reported values did also not affect the effect significance or direction for either climate, type of vegetation change or metric. The only changes occurred in local effects that became significantly negative and effects on microbes that became non-significant (Figure S6, Table S3). For the individual trophic levels, the significance level changed only in 9 out of 72 cases, most often with significant effects becoming non-significant except for effects of changes at local scales on shredders that became significantly negative, and whenever effect sizes changed in directionality, these changes never resulted in statistically significant effects (Figure S7).

## Discussion

Our systematic review of 144 studies showed that human-induced land-cover changes in the watershed negatively affected freshwater detrital food webs. This negative effect of land-cover change on freshwater ecosystems was moderated by type of vegetation change, spatial scale of vegetation change, metric and trophic level. Moreover, our results show that different trophic levels within the freshwater detrital food web were influenced by distinct moderators. Studies observing negative effects of one trophic level most often reported also negative effects of another trophic level. This simultaneous decrease highlights that land-cover changes often negatively affect multiple detrital food web components simultaneously. Together, these comprehensive results emphasise the pressing need to conserve naturally vegetated watersheds for freshwater food webs and to consider cross-ecosystem consequences in land-use/land-cover management.

### Overall effect of vegetation change on freshwater detrital food webs

Even after accounting for potentially missing studies, the responses of freshwater detrital food webs remained mostly significantly negative and, hence, corroborate strongly that land-cover changes cascade across ecosystem boundaries and reduce biodiversity and ecosystem processes in adjacent ecosystems (Figure 3A; Allan 2004; Hanna *et al*. 2020; IPBES 2019; Tiegs *et al*. 2024). Our robust, new results not only confirm but also expand and refine our current understanding of the global consequences of harvesting (Richardson & Béraud 2014), exotic tree plantations (Ferreira *et al*. 2016, 2019), and other land-cover changes on freshwater biodiversity (Camana *et al*. 2024; Petsch *et al*. 2021) and ecosystem processes (Brauns *et al*. 2022; Tiegs *et al*. 2024) through the joint analysis of outcomes from the largest dataset of studies in this field. Still, although our dataset covered a broad latitudinal gradient, we observed a significant underrepresentation of studies from Africa and Asia compared to other regions of the world, highlighting a geographical bias in the existing literature (Figure 2A). This disparity may be attributed to limited research funding, infrastructure, and capacity, as well as potential dissemination biases that favour studies from more developed regions and published in English. Grey literature and literature published in languages other than English could have potentially filled some of these gaps and created a more comprehensive picture (O’Dea *et al*. 2021) of our observed negative effects of land-cover changes on freshwater detrital food webs.

### Moderator effects on food webs

The negative impact of human-induced vegetation changes in watersheds on freshwater detrital food webs was not primarily moderated by the climate zone but rather by the type and spatial scale of vegetation change, metrics, and trophic levels involved (Figure 3, Table 1). These findings underscore the critical role of these moderating factors in shaping aquatic ecosystems and determining how food webs respond to land-cover changes across the world.

While we found negative effect sizes of land-cover changes on freshwater detrital food webs in the tropics and subtropics, boreal comparisons resulted in no statistically significant effect sizes due to large variance caused by strong differences in effect sizes between and within studies (Figure 3B). This large variance stems from moderators other than types of vegetation change, as our boreal dataset only included effects of harvesting (i.e., forestry management including logging and clear-cutting). In these boreal forest ecosystems, stand age (i.e., age since clear cut) seems to be a particularly important factor leading to non-linear and increasingly variable responses in adjacent detrital food webs (Frainer & McKie 2021). Apart from the specific forestry management, the effects of harvesting might also depend on whether the local riparian forest consisted of deciduous or coniferous species (Kominoski *et al*. 2011) or the width of riparian buffers (Jyväsjärvi *et al*. 2020).

While the overall effects of harvest and land-use conversions on freshwater detrital food webs were negative, we found no statistically significant effects of restoration and plantations (Figure 3C). For restoration, these outcomes likely stem from different goals and approaches (e.g., study design, definition of reference conditions, timing of measurements) with some studies measuring the effects of replanting native riparian vegetation (Giling *et al*. 2015) while others focused on removing invasive species from the riparian vegetation (McNeish *et al*. 2017). These results highlight the context-dependency of the observed restoration measures (Nilsson *et al*. 2015) and–together with the small number of available studies on restoration effects–the general need for more studies reporting a before-after or control-intervention of vegetation restoration on aquatic food webs (Bellmore *et al*. 2017; Naiman *et al*. 2012; Weber *et al*. 2018; see Loch *et al*. (2020) for a review on food web restoration studies). More quantitative assessments of restoration effects would also increase the efficacy of restoration projects to reach ecological goals (Palmer 2009). For plantations, overall effects were non-significant, due to the opposing responses of different trophic levels, with a decrease for omnivores and increase for microbes and detritus (Figure 4).

With respect to spatial scale of vegetation change, the freshwater detrital food web was most affected when land-cover change occurred at catchment and local+catchment scale (Figure 3D) in line with previous studies (Ferreira *et al*. 2016). While local changes in the riparian vegetation did not result in statistically significant effects in the freshwater detrital food web, it needs to be noted that the intactness of riparian vegetation is still crucial for many aquatic organisms and essential for their survival as demonstrated in numerous studies (e.g., Casotti *et al*. 2015; Encalada *et al*. 2010; Giling *et al*. 2015; Kominoski *et al*. 2011; Masese *et al*. 2014; Oester *et al*. 2023).

We found negative effects for all metrics considered in this meta-analysis to assess the community and ecosystem processes in the freshwater detrital food web to changes in the watershed vegetation (Figure 3E). The most negative effects on decomposition occurred when total decomposition and shredder-mediated decomposition were assessed. These metrics also reflect the activity of higher consumers and how well they can perform their ecological functions (Cummins *et al*. 1989; Gessner *et al*. 2010; Handa *et al*. 2014). Hence, watershed vegetation not only influences the communities but also the ecosystem processes in freshwaters.

For the moderator of trophic level, we found the most negative overall effect sizes for microbes and shredders (Figure 3F). These negative effects are not surprising as the shredders and microbes observed in these studies are known to be highly specialised and dependent on detritus (Cummins *et al*. 1989; Wallace & Webster 1996). Moreover, many shredders and other macroinvertebrate groups also depend on the terrestrial environment during their terrestrial adult stages, when they use local and upstream riparian vegetation for shelter, refuge, courtship, and mating (Reinhart & VandeVoort 2006; Vilela & Sanmartín-Villar 2019; Yoshimura 2012). Hence, to protect these sensitive taxa, a natural vegetation is crucial (Cummins *et al*. 1989; Tolkkinen *et al*. 2020).

### Moderator effects on individual trophic levels

Deconstructing the response to land-cover changes in the detrital food web into different trophic levels (Figure 4, Table 2), we found that detritus, microbes and shredders were influenced by distinct moderators. Together, these new results shed light on distinct drivers of different trophic levels of the freshwater detrital food web and help us better understand the intricate relationships within ecological communities in nature.

**Table 2.**
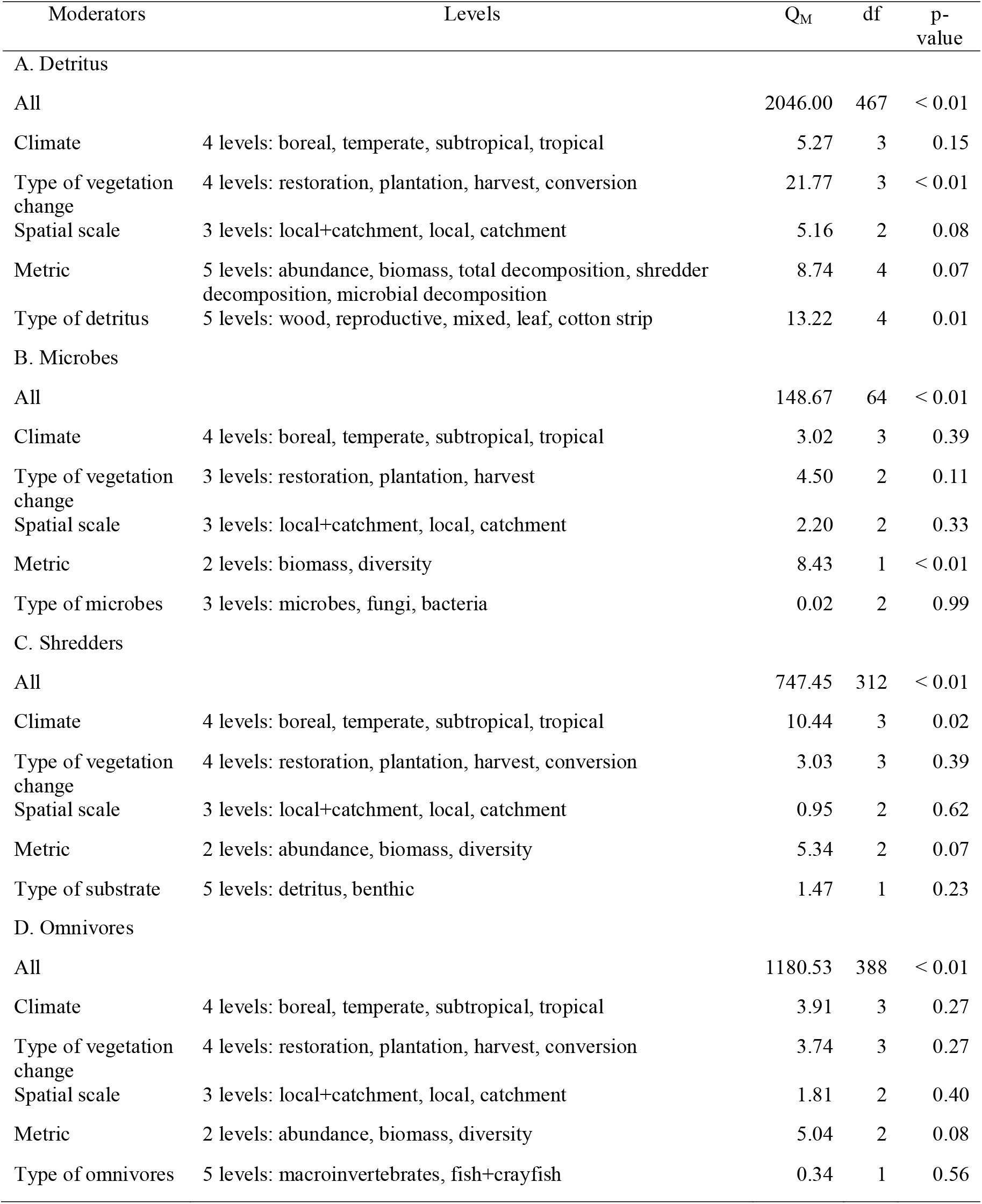
Effects of moderators tested for different trophic levels. Test for heterogeneity between levels within moderators (Q_M_), degrees of freedom (df) and p-values (significant differences among moderator levels exist if p-values < 0.05; see Figure 4) are shown. Rosenberg’s fail-safe number (N_fs_) for the datasets of detritus (N_fs_ = 4675, n = 468) and shredders (N_fs_ = 4661, n = 313) indicate the robustness (N_fs_ > 5 × n +10) of the datasets to publication bias; for omnivores (N_fs_ = 1936, n = 389), results need to be interpreted with caution as the N_fs_ is higher than the threshold for considering the dataset robust to publication bias (see also Figure S5); the effect of forest change on the microbial dataset was non- significant (Figure 4B) and hence there is no N_fs_.

While effect sizes for detritus were negative in harvested and converted vegetation, they were positive in plantations (Figure 4A). Depending on the condition and composition of watershed vegetation, the quantity, quality and timing of detritus entering the water body and its subsequent decomposition can vary substantially (Ferreira *et al*. 2019; Larrañaga *et al*. 2023; Tonin *et al*. 2017). It needs to be noted that faster decomposition not always linearly scales with and represents the ecological state of an ecosystem (Frainer *et al*. 2021; Masese *et al*. 2014; Truchy *et al*. 2022; Woodward *et al*. 2012). For instance, detritus from species used in plantations might be more nutritious, more palatable or more available for the detrital food web than detritus from the pristine vegetation (Castro *et al*. 2018; Collier & Halliday 2000; Tank *et al*. 2010). While non-native eucalyptus plantations often have negative impacts on detrital food webs (Ferreira *et al*. 2016, 2019; Graça *et al*. 2002), the direction of the effects of conifer (e.g., Riipinen *et al*. 2009; Sakai *et al*. 2013), palm oil (Chellaiah & Yule 2018), citrus (Dézerald *et al*. 2014), coffee and rubber (Walsh *et al*. 2002), or banana plantations (Casotti *et al*. 2015; Kiffer Jr *et al*. 2018) depended on the reference conditions, resource environment and metrics considered.

However, the responses of consumers like microbes and shredders varied with community metrics (Figure 4B) and climate zones (Figure 4C), respectively. As expected, shredders in tropics and subtropics, where deforestation and other land-cover changes are currently most drastic, showed the most negative responses to watershed vegetation changes. Tropic and subtropic freshwaters as biodiversity hotspots are still poorly understood in terms of biodiversity and ecosystem processes and, hence, more research and conservation in these areas is urgently needed to not only assess and describe local communities but also to examine cross-ecosystem consequences of ongoing land-cover changes (Boyero *et al*. 2012, 2021; Encalada *et al*. 2010).

### Multitrophic effects

We observed negative effects due to land-cover changes in watershed vegetation not only within a single trophic level of the detrital food web but also often simultaneously for multiple trophic levels (Figure 5). Although we may be missing studies reporting positive effects on omnivores (Figure S5; Table S2), the datasets for other trophic groups were robust (Figures S1–4; Table S2). These results suggest two possibilities: either the land-cover change was so severe that multiple trophic levels reacted negatively independently, or the decrease in one trophic level cascaded through the food web affecting other trophic levels (Barnes *et al*. 2017; Knight *et al*. 2005). Regardless, shifts in the freshwater detrital food web—whether through community changes or alterations in trophic interactions and ecosystem processes— can have severe consequences for trophic pathways to higher-order consumers in both aquatic and terrestrial environments (Albertson *et al*. 2018; Ballinger & Lake 2006; Nakano & Murakami 2001).

## Conclusion

In conclusion, our findings reveal the overall negative impact of land-cover changes in watershed vegetation on freshwater detrital food webs, with no indication that restoration measures were able to revert these, at least over the time-scales covered by the different studies. Moreover, our results show that different trophic levels were influenced by distinct moderators. These findings thus highlight the urgent need for the conservation and more effective restoration of watersheds and consider multiple metrics and trophic levels when assessing vegetation changes as they differ in their sensitivity to changes. Hence, land-cover management must account for cross-ecosystem consequences at both local and catchment scales. Adopting holistic conservation strategies that account for ecological complexity including multiple trophic levels, trophic dependencies and ecosystem processes is crucial for preserving and restoring freshwater ecosystems in landscapes with increasing human pressure (Bellmore *et al*. 2017; Harrison *et al*. 2018; Naiman *et al*. 2012).

## Acknowledgment

We thank the members of the Freshwater Biodiversity and Functional Ecology Lab at SUPSI, the Lab for Aquatic Insect Ecology at UVV, and the Altermatt Lab at EAWAG for their support, inputs, and feedback. Many thanks go to the numerous authors providing data and answering inquiries about their studies. Lastly, we thank the funding agencies SNSF, URPP, FAPES, CNPq and FCT.

## Author contribution

Conceptualisation: RO, FK, MM, FA, VF, AB; Methodology: RO, FK, VF; Writing: RO, FK, MM, FA, VF, AB; Funding acquisition: MM, FA, AB; Supervision: MM, FA, VF, AB

## Data availability

The data and code that support the findings of this study are openly available on Dryad and Zenodo: https://doi.org/10.5061/dryad.c866t1ghg

