## Supplementary Material for "A global synthesis on land-cover changes in watersheds shaping freshwater detrital food webs"

**Affiliations**
1 Institute of Microbiology, University of Applied Sciences and Arts of Southern Switzerland, via Flora Ruchat Roncati 15, CH-6850 Mendrisio, Switzerland

2 Department of Evolutionary Biology and Environmental Studies, University of Zurich, Winterthurerstr. 190, CH-8057 Zurich, Switzerland

3 Eawag: Swiss Federal Institute of Aquatic Science and Technology, Department of Aquatic Ecology, Überlandstrasse 133, CH-8600 Dübendorf, Switzerland

4 Laboratory of Aquatic Insect Ecology, Universidade Vila Velha, Av. Comissário José Dantas de Melo 21, 29102-920 Vila Velha, ES, Brazil

5 MARE – Marine and Environmental Sciences Centre, ARNET – Aquatic Research Network, Department of Life Sciences, University of Coimbra, Calçada Martim de Freitas, 3000–456 Coimbra, Portugal

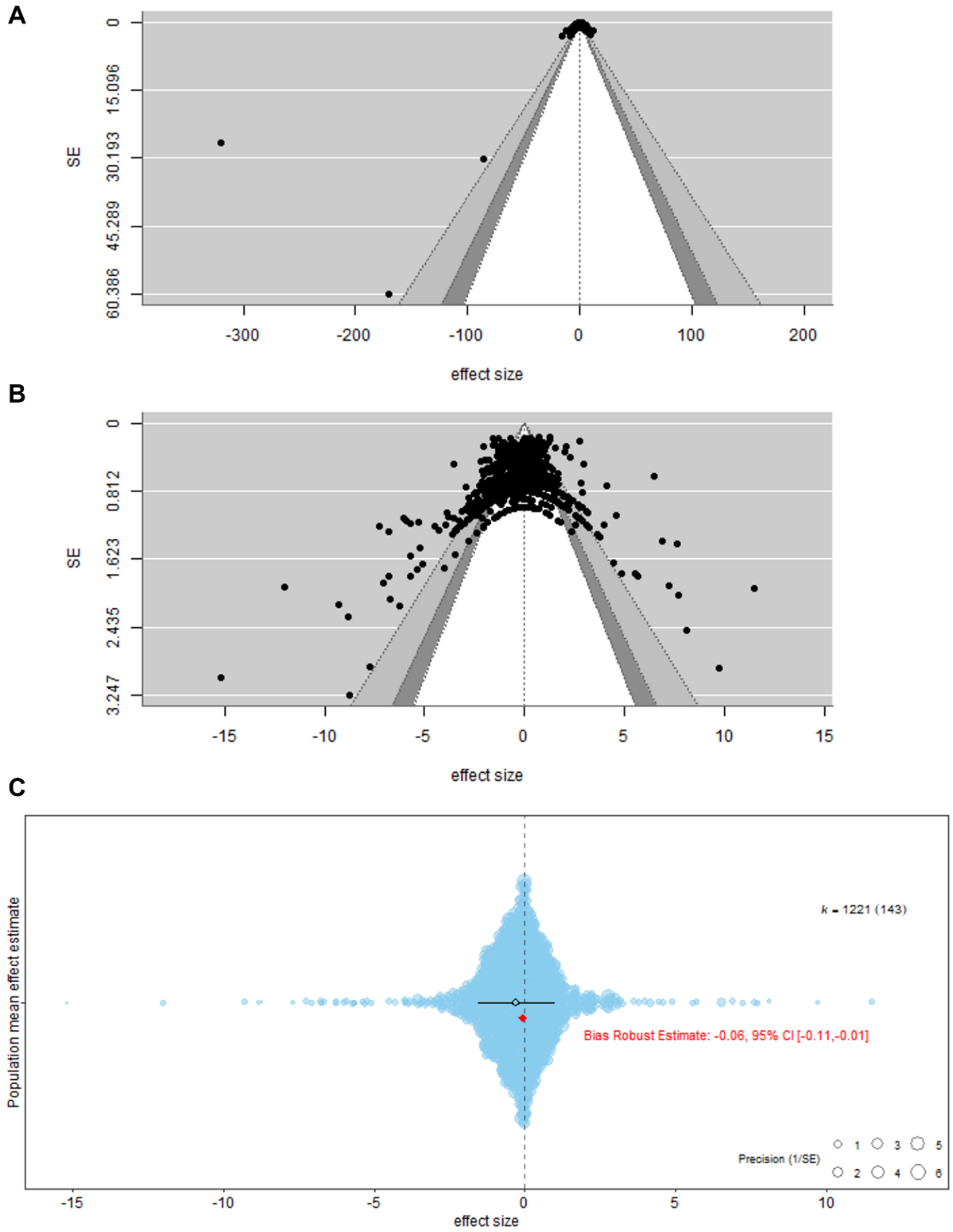


**Figure S1.** Funnel plots for the **whole dataset** with (A) and without (B) three outlier values with extreme effect sizes (all three outlier comparisons stem from shredder-mediated decomposition rates calculated as $k_c/k_f$ from tropical leaves reported directly from the authors or in the text from the studies 19 and 84 (see Table S3 for whole reference)) and publication bias represented as an orchard plot of the whole dataset without outliers (C). This orchard plot illustrates the distribution of standardised mean differences (Hedges' g) from 1221 comparisons (from 143 studies) evaluating the population mean effect estimate. Each blue circle corresponds to an effect size, with the size of the circle reflecting the precision (1/SE) of the estimate. The black horizontal lines show the overall mean effect size with a 95 % (thick) and 99 % (thin) line as confidence intervals (CI). The bias robust estimate is highlighted in red, showing Hedges' g of -0.06 with a 95 % CI ranging from -0.11 to -0.01. This suggests a small but statistically significant negative effect, accounting for potential bias in the data and that there is no evidence of publication bias in our data because the interpretation of results does not change after accounting for potential bias in the data.

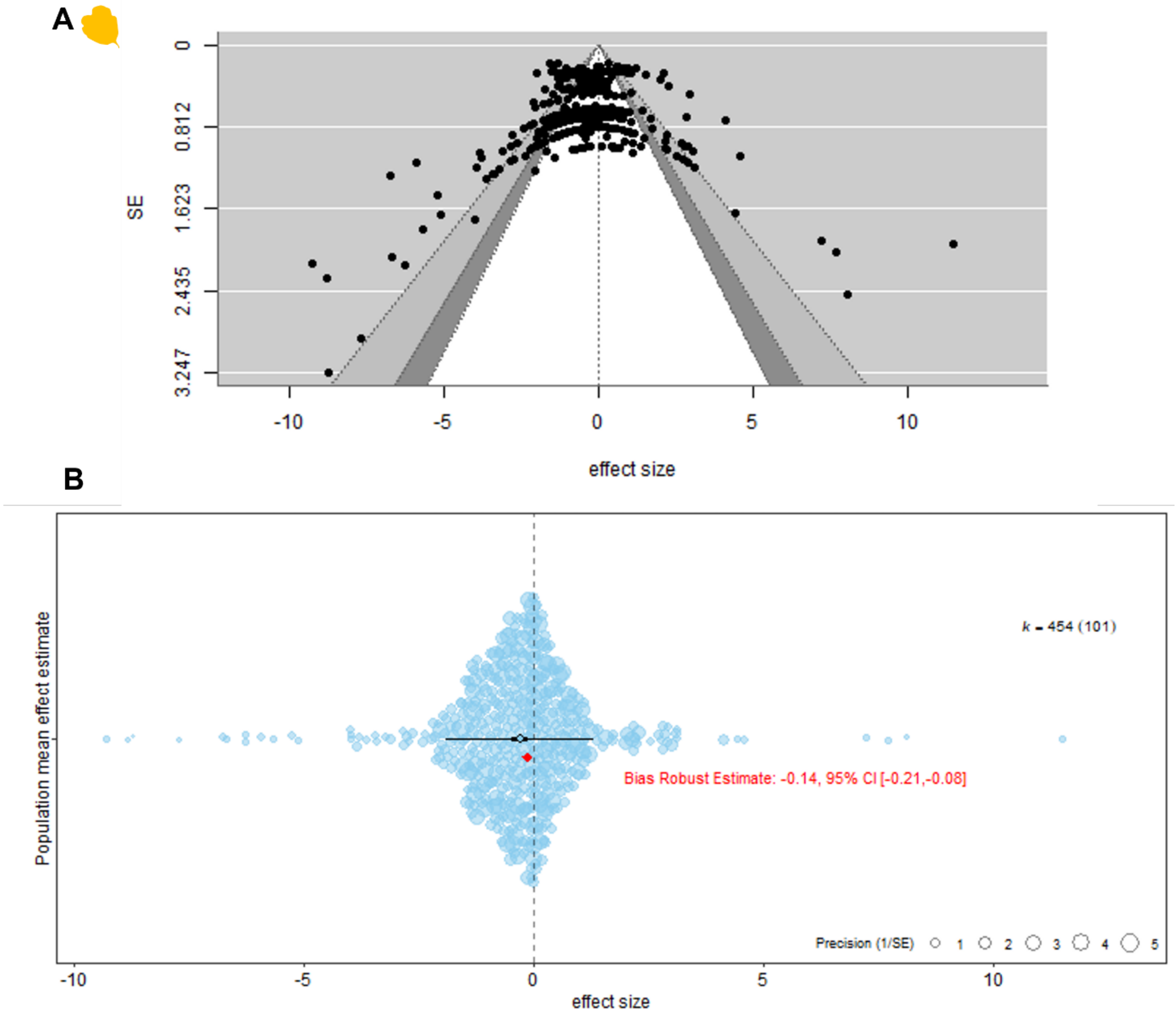


**Figure S2.** Funnel plots for the **detritus** (A) and publication bias represented as an orchard plot of this dataset (B). This orchard plot illustrates the distribution of standardised mean differences (Hedges' g) from 454 comparisons (from 101 studies) evaluating the population mean effect estimate. Each blue circle corresponds to an effect size, with the size of the circle reflecting the precision (1/SE) of the estimate. The black horizontal lines show the overall mean effect size with a 95 % (thick) and 99 % (thin) line as confidence intervals (CI). The bias robust estimate is highlighted in red, showing Hedges' g of -0.14 with a 95 % CI ranging from -0.21 to -0.08. This suggests a small but statistically significant negative effect, accounting for potential bias in the data and that there is no evidence of publication bias in our data because the interpretation of results does not change after accounting for potential bias in the data.

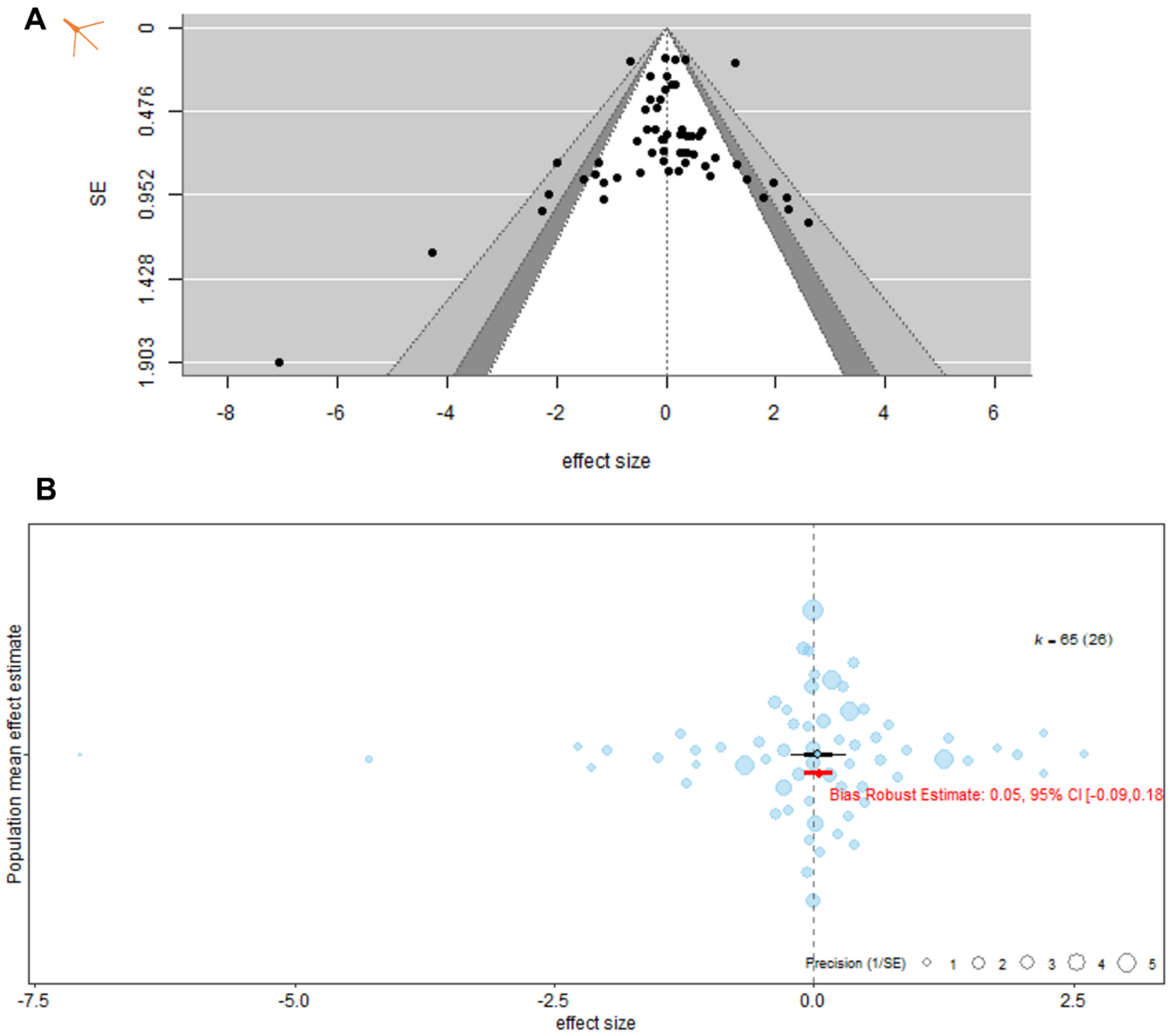


**Figure S3.** Funnel plots for the (A) and publication bias represented as an orchard plot of this dataset (B). This orchard plot illustrates the distribution of standardised mean differences (Hedges' g) from 65 comparisons (from 26 studies) evaluating the population mean effect estimate. Each blue circle corresponds to an effect size, with the size of the circle reflecting the precision (1/SE) of the estimate. The black horizontal lines show the overall mean effect size with a 95 % (thick) and 99 % (thin) line as confidence intervals (CI). The bias robust estimate is highlighted in red, showing Hedges' g of 0.05 with a 95 % CI ranging from -0.09 to 0.18. This suggests statistically non-significant effects, accounting for potential bias in the data and that there is no evidence of publication bias in our data because the interpretation of results does not change after accounting for potential bias in the data.

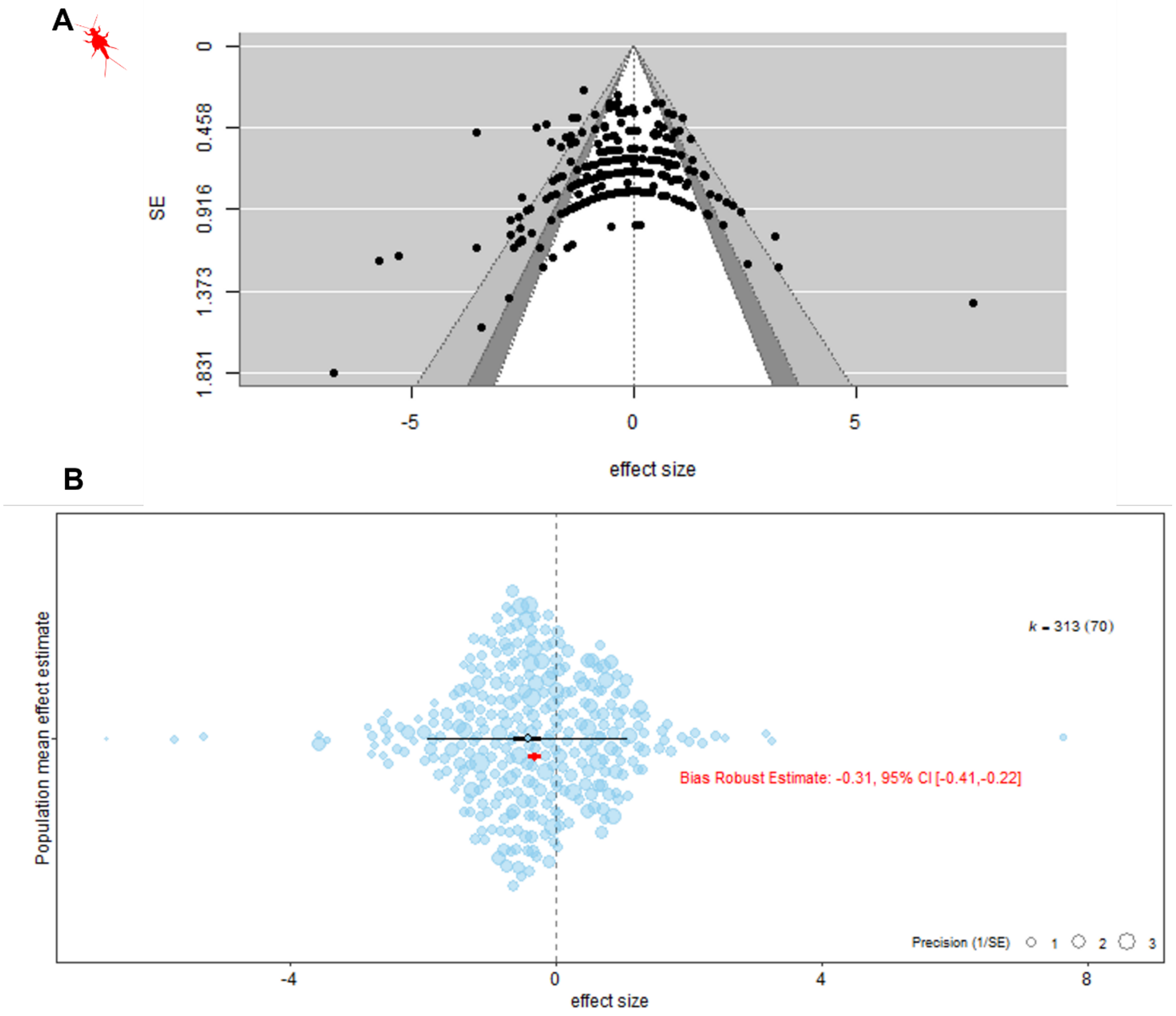


**Figure S4.** Funnel plots for the **shredders** (A) and publication bias represented as an orchard plot of this dataset (B). This orchard plot illustrates the distribution of standardised mean differences (Hedges' g) from 313 comparisons (from 70 studies) evaluating the population mean effect estimate. Each blue circle corresponds to an effect size, with the size of the circle reflecting the precision (1/SE) of the estimate. The black horizontal lines show the overall mean effect size with a 95 % (thick) and 99 % (thin) line as confidence intervals (CI). The bias robust estimate is highlighted in red, showing Hedges' g of -0.31 with a 95 % CI ranging from -0.41 to -0.22. This suggests statistically significant negative effect, accounting for potential bias in the data and that there is no evidence of publication bias in our data because the interpretation of results does not change after accounting for potential bias in the data.

.

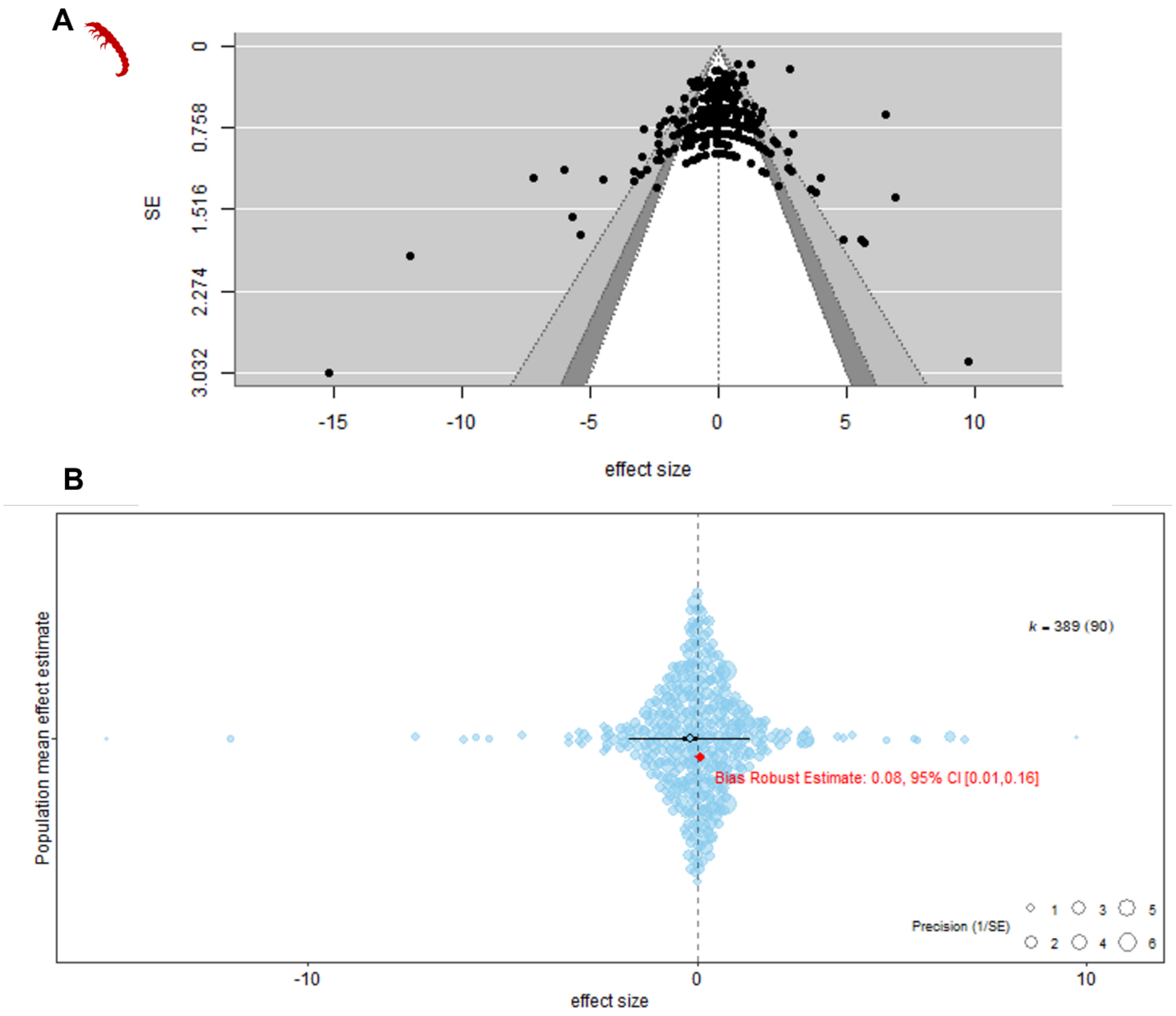


**Figure S5.** Funnel plots for the **omnivores** (A) and publication bias represented as an orchard plot of this dataset (B). This orchard plot illustrates the distribution of standardised mean differences (Hedges' g) from 389 comparisons (from 90 studies) evaluating the population mean effect estimate. Each blue circle corresponds to an effect size, with the size of the circle reflecting the precision (1/SE) of the estimate. The black horizontal lines show the overall mean effect size with a 95 % (thick) and 99 % (thin) line as confidence intervals (CI). The bias robust estimate is highlighted in red, showing a Hedges' g of 0.08 with a 95 % CI ranging from 0.01 to 0.16. This suggests that there might be bias in the data.

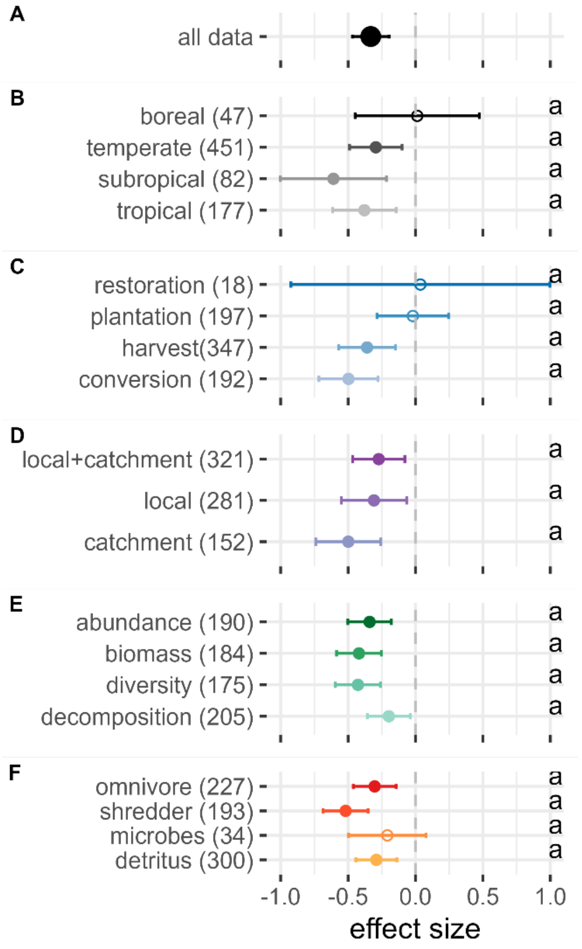


**Figure S6.** Impacts of human-induced alterations in the watershed vegetation on freshwater detrital food webs from only reported comparions (i.e., excluding imputed and estimated values). The global response (all data) is shown on the first row A) and is separated by moderators in the following rows for B) climate, C) type of vegetation change, D) spatial scale, E) metric and F) trophic level. The numbers in brackets after each moderator level represent the number of comparisons. For each moderator level, the circle represents the mean effect size with 95 % confidence intervals (CIs) computed from the random effects model. Filled circles indicate statistically significant effects, whereas empty circles have CIs that cross the 0-line and are thus statistically non-significant. Negative effects sizes (standardised mean differences Hedges' g between altered and reference vegetation conditions < 0) indicate that the values in the altered conditions were lower compared to the reference conditions. Groups sharing the same letter (e.g., 'a', 'ab', 'b') are not significantly different from each other as their CIs overlap.

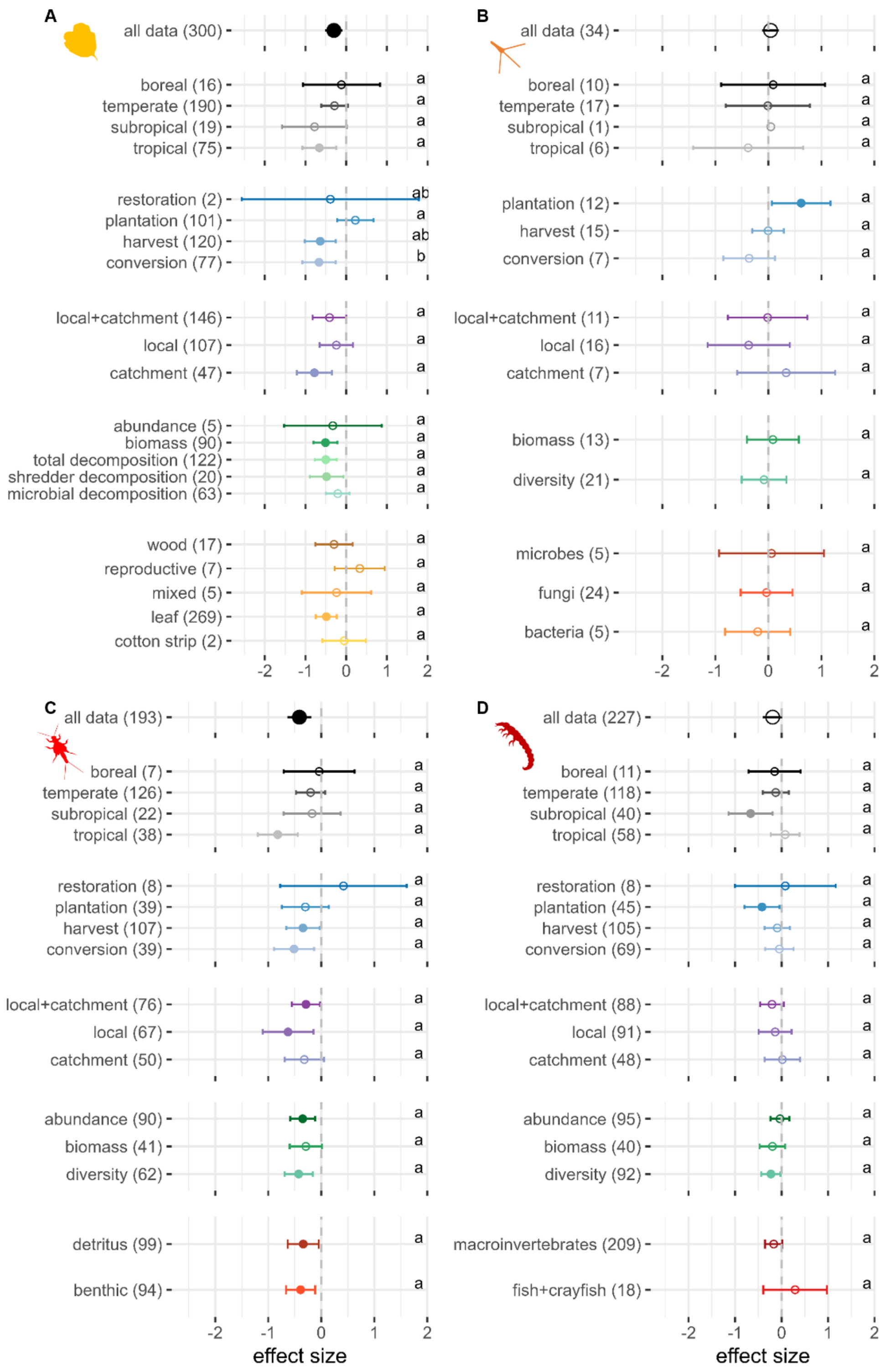


**Figure S7.** Moderator analysis separated for the subsets of each trophic level from only reported comparions (i.e., excluding imputed and estimated values) with A) detritus, B) microbes, C) shredders and D) omnivores. For each moderator level, the circle represents the mean effect size with 95 % confidence intervals (CIs) computed from the random effects model. Filled circles indicate statistically significant effects, whereas empty circles have CIs that cross the 0-line and are thus statistically non-significant. Negative effects sizes (standardised mean differences Hedges' g between altered and reference vegetation conditions < 0) indicate that the values in the altered conditions were lower compared to the reference conditions. Groups sharing the same letter (e.g., 'a', 'ab', 'b') are not significantly different from each other as their CIs overlap.

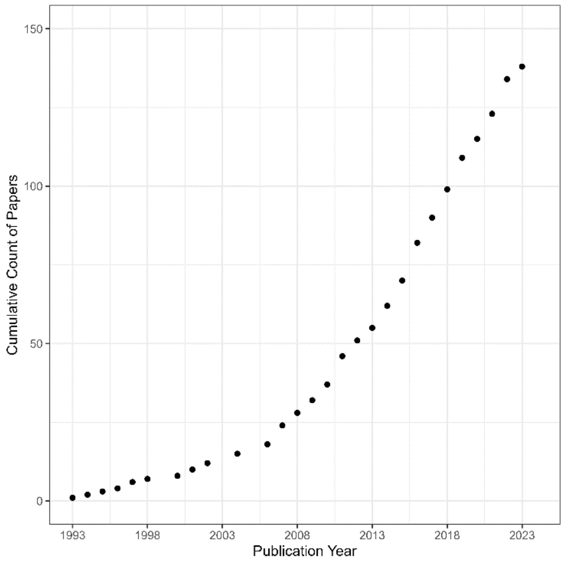


**Figure S8.** Cumulative number of publications (n = 144).

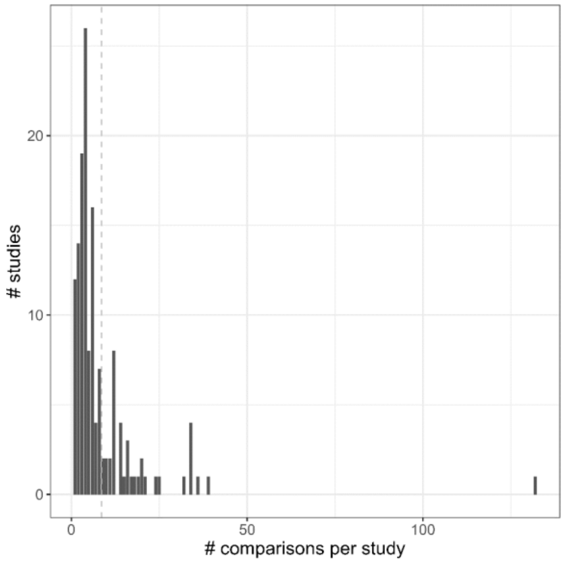


**Figure S9.** Histogram of data distribution with number of observations per study. The mean of 8.6 observations per study is indicated with the grey dashed line.

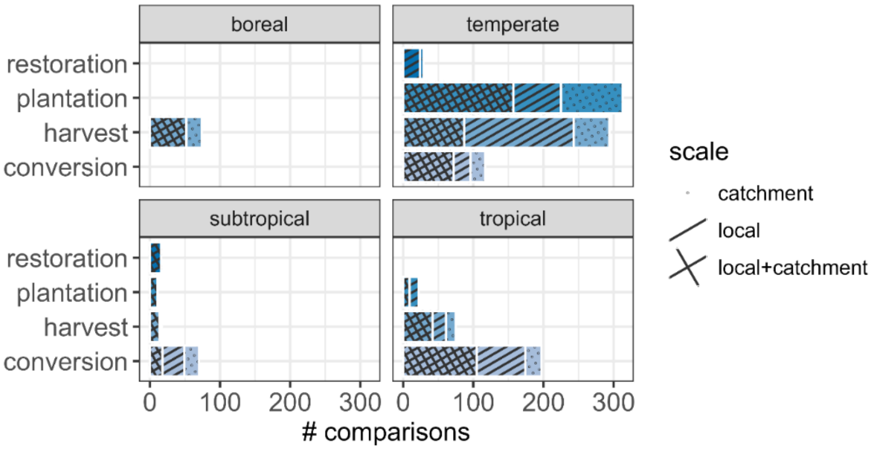


**Figure S10.** Distribution of comparisons between different climate (panels), types of vegetation change (bars) and spatial scales of vegetation change (pattern) with dotted bars indicating vegetation change at a catchment scale, striped bars indicating local changes and criss-crossed bars depicting local+catchment scale changes.

**Table S1**. Identity, levels and definitions of moderators used in this review.

| Moderator | Levels | Definition |
|---|---|---|
| study ID | several | Full references are given in Table S3 |
| year | continuous | Publication year |
| latitude | continuous | Latitude in decimal degrees |
| longitude | continuous | Longitude in decimal degrees |
| water body | several | Water body as described in the primary study, e.g., stream, river, bromeliad |
| climate | boreal | Boreal climate zone |
| | temperate | Temperate climate zone |
| | sub-tropical | Sub-tropical climate zone |
| | tropical | Tropical climate zone |
| scale | local | Forest change only occurs in the riparian area, not at the catchment level (e.g., Kennedy & Hobbie, 2004). |
| | catchment | Forest change occurs at the catchment level but not in the riparian area, where native vegetation is still present (e.g., Martínez et al., 2013). |
| | local+catchment | Forest change occurs at the catchment and riparian level (e.g., Lecerf et al., 2005). |
| type of vegetation change | restoration | Direct anthropogenic alteration through restoration measures aiming at restoring the native, natural riparian vegetation (e.g., Giling et al., 2016; McNeish et al., 2016) |
| | harvest | Direct anthropogenic alteration through harvesting or reducing the volume (e.g., logging or clear-cutting) of the natural riparian vegetation in the form of forestry management (e.g., Kominoski et al., 2011; Musetta-Lambert et al. 2017) |
| | land use conversion | Direct anthropogenic alteration through changes in the vegetation type (e.g., from forest to pasture) resulting in overall degradation, reduction in width and cover of natural riparian vegetation (e.g., Ferreira et al., 2015; Ono et al., 2020) |
| | plantation | Direct anthropogenic alteration through replacement of native forests by other types of riparian vegetation such as *Eucalyptus* or conifers (e.g., Hladyz et al., 2011, Martínez et al., 2013). |
| reference condition | several | Description of reference condition of the riparian vegetation as reported in the primary study |
| altered condition | several | Description of altered condition of the riparian vegetation as reported in the primary study |
| trophy | 1 | Detritus (e.g., leaf litter, wood) |
| | 2 | Microbial community (e.g., aquatic fungi, bacteria) |
| | 3 | Shredder community |
| | 4 | Omnivore community |
| type of detritus | leaf | Detritus in the form of leaves from riparian tree species (e.g., Oester et al. 2023) |
| | wood | Detritus in the form of wood, including bark, branches and twigs (e.g., Valente-Neto et al., 2015) |
| | mixed | Detritus in the form of leaves and wood mixed together (e.g., Ono et al., 2020) |
| | cotton strip | Detritus in the form of cotton strips (e.g., Mancuso et al., 2023) |
| | reproductive | Detritus in the form of reproductive material such as fruit, seeds, flowers, cones (e.g., Kadeka et al. 2021) |
| taxon of detritus | several | Detritus taxa as reported in the primary study |
| type of microbes | bacteria | Microbial community of bacteria |
| | fungi | Microbial community of fungi |
| | microbe | Microbial community of microbes (not specified or a mix of bacteria, fungi and algae) |
| type of consumers | macroinvertebrates | Macroinvertebrate community (i.e., insect larva, molluscs, small crustaceans, etc.) |
| | shredders | Detritivorous macroinvertebrates based on the functional guild of shredders (i.e., macroinvertebrate preferentially shredding on coarse organic matter) |
| | fish+crayfish | Detritivorous fish and crayfish community |
| type of substrate | CPOM | Location of sample collection based on CPOM (coarse particulate organic matter) |
| | benthic | Location of sample collection based on the benthic habitat |
| | water column | Location of sample collection based on the water column |
| taxon of substrate | several | Taxon name of the CPOM (e.g. species name of the leaf litter origin) |
| metric | a-diversity | α-diversity provided by the primary study (preferentially richness, but if not provided also other metrics were considered) |
| | abundance | Abundance of microbes or invertebrates provided in the primary study |
| | biomass | Biomass of microbes or invertebrates provided in the primary study |
| | Total decompositon | Total decomposition of detritus |

| | | |
|---|---|---|
| | Shredder-mediated decomposition | Part of total decomposition mediated by shredders and omnivores |
| | Microbial decomposition | Part of total decomposition provided by microbes |
| unit | several | Unit of α-diversity, abundance, biomass or decomposition as reported in the primary study (the only exception was that decomposition reported as "% mass remaining" was converted to "% mass lost", by subtracting "%mass remaining" from 100 %, to be consistent with the other metrics such as decomposition rates and g mass lost) |
| data origin | provided | Data derived from either text, tables or authors directly |
| | estimated | Data derived from figures and extracted through WebPlotDigitizer |
| imputation | original | Standard deviation derived from study |
| | imputed | Standard deviation imputed based on similar studies (in terms of fw_category and metric) following Koricheva et al. (2013). |

**Table S2**. Datasets, moderators and levels tested in the analyses, sample size (n), Rosenberg's fail-safe number ($N_{fs}$) (dataset is robust to publication bias if $N_{fs} > 5 \times n + 10$, with n= number of effect sizes), estimated mean effect size Hedges' g (standardized mean differences between altered and reference riparian vegetation conditions) with 95 % confidence intervals (CI) computed from the random effect model (negative effects sizes indicate that the values in the altered conditions were lower compared to the reference conditions), p-values (significant effects exist if $p < 0.05$), percentage of total variation that is due to between-study variation ($I^2$) and test for heterogeneity between levels within moderators ($Q_M$).

| Data | Moderator | Levels | n | $N_{fs}$ | Estimate [95% CI] | p-value | $I^2$ (%) |
|---|---|---|---|---|---|---|---|
| Detritus | Entire dataset | | 468 | 4675 | -0.30 [-0.49,-0.12] | < 0.01 | 94.61 |
| | Climate | Boreal | 32 | | -0.28 [-0.87,0.31] | 0.35 | 88.31 |
| | | Temperate | 285 | | -0.17 [-0.40,0.06] | 0.14 | 95.10 |
| | | Sub-Tropical | 25 | | -0.29 [-0.94,0.36] | 0.37 | 72.12 |
| | | Tropical | 126 | | -0.65 [-1.01,-0.29] | < 0.01 | 93.08 |
| | Metric | Abundance | 10 | | -0.08 [-0.90,0.74] | 0.84 | 72.89 |
| | | Biomass | 124 | | -0.38 [-0.61,-0.15] | < 0.01 | 85.85 |
| | | Microbial decomposition | 103 | | -0.16 [-0.38,0.05] | 0.14 | 82.83 |
| | | Shredder-mediated decomposition | 35 | | -0.41 [-0.72,-0.11] | < 0.01 | 51.82 |
| | | Total decomposition | 196 | | -0.33 [-0.52,-0.13] | < 0.01 | 89.72 |
| | Type | Leaf | 416 | | -0.30 [-0.49,-0.11] | < 0.01 | 94.11 |
| | | Cotton strip | 5 | | 0.16 [-0.31,0.63] | 0.50 | 36.77 |
| | | Mixed | 11 | | -0.91 [-1.58,-0.24] | < 0.01 | 45.35 |
| | | Reproductive | 7 | | 0.31 [-0.24,0.86] | 0.27 | 27.60 |
| | | Wood | 29 | | -0.28 [-0.65,0.09] | 0.14 | 83.99 |
| | Vegetation Change | Conversion | 152 | | -0.48 [-0.78,-0.19] | < 0.01 | 91.05 |
| | | Harvest | 160 | | -0.55 [-0.84,-0.26] | < 0.01 | 93.19 |
| | | Plantation | 140 | | 0.34 [0.01,0.66] | 0.04 | 95.81 |
| | | Restoration | 16 | | -0.62 [-1.39,0.15] | 0.11 | 78.50 |
| | Scale | Catchment | 70 | | -0.51 [-0.85,-0.18] | < 0.01 | 90.67 |
| | | Local | 145 | | -0.08 [-0.39,0.23] | 0.61 | 85.62 |
| | | Local+Catchment | 253 | | -0.35 [-0.64,-0.07] | 0.01 | 96.62 |
| Microbes | Entire dataset | | 65 | | 0.04 [-0.09,0.18] | 0.54 | 37.23 |
| | Climate | Boreal | 15 | | 0.09 [-0.22,0.40] | 0.59 | 10.49 |
| | | Temperate | 40 | | 0.08 [-0.08,0.24] | 0.32 | 42.25 |
| | | Sub-Tropical | 1 | | 0.05 [-1.57,1.66] | 0.96 | 1.82 |
| | | Tropical | 9 | | -0.33 [-0.78,0.11] | 0.14 | 8.35 |
| | Metric | Biomass | 36 | | -0.15 [-0.35,0.05] | 0.15 | 48.93 |
| | | Diversity | 29 | | 0.23 [0.01,0.45] | 0.04 | 44.91 |
| | Type | Microbe | 7 | | 0.07 [-0.41,0.54] | 0.78 | 12.02 |
| | | Bacteria | 9 | | 0.04 [-0.20,0.28] | 0.74 | 31.31 |
| | | Fungi | 49 | | 0.03 [-0.13,0.20] | 0.71 | 33.26 |
| | Vegetation Change | Conversion | 17 | | -0.11 [-0.41,0.19] | 0.48 | 68.64 |
| | | Harvest | 29 | | -0.04 [-0.27,0.20] | 0.77 | 55.50 |
| | | Plantation | 19 | | 0.42 [0.01,0.81] | 0.05 | 21.88 |
| | Scale | Catchment | 27 | | 0.14 [-0.05,0.32] | 0.15 | 45.04 |
| | | Local | 20 | | -0.09 [-0.37,0.19] | 0.53 | 26.33 |
| | | Local+Catchment | 18 | | -0.04 [-0.34,0.25] | 0.77 | 10.36 |
| Shredders | Entire dataset | | 313 | 4661 | -0.41 [-0.62,-0.2] | < 0.01 | 88.54 |
| | Climate | Boreal | 12 | | -0.09 [-0.72,0.54] | 0.79 | 76.27 |
| | | Temperate | 197 | | -0.14 [-0.42,0.13] | 0.3 | 89.02 |
| | | Sub-Tropical | 33 | | -0.80 [-1.28,-0.33] | < 0.01 | 79.24 |
| | | Tropical | 71 | | -0.74 [-1.11,-0.38] | < 0.01 | 84.32 |
| | Metric | Abundance | 151 | | -0.38 [-0.60,-0.16] | < 0.01 | 80.02 |
| | | Biomass | 64 | | -0.27 [-0.55,0.01] | 0.06 | 65.69 |
| | | Diversity | 98 | | -0.55 [-0.79,-0.30] | < 0.01 | 70.94 |
| | Type | Benthic | 147 | | -0.49 [-0.74,-0.24] | < 0.01 | 82.48 |
| | | CPOM | 166 | | -0.34 [-0.58,-0.09] | 0.01 | 84.15 |
| | Vegetation Change | Conversion | 84 | | -0.59 [-0.91,-0.27] | < 0.01 | 82.34 |
| | | Harvest | 119 | | -0.41 [-0.74,-0.07] | 0.02 | 87.96 |
| | | Plantation | 99 | | -0.15 [-0.55,0.25] | 0.45 | 89.63 |
| | | Restoration | 11 | | -0.15 [-1.15,0.84] | 0.76 | 88.95 |
| | Scale | Catchment | 65 | | -0.56 [-0.94,-0.19] | < 0.01 | 84.79 |
| | | Local | 98 | | -0.30 [-0.72,0.11] | 0.15 | 88.57 |
| | | Local+Catchment | 150 | | -0.41 [-0.69,-0.13] | < 0.01 | 87.22 |
| Omnivores | Entire dataset | | 389 | 1936 | -0.19 [-0.38,0.00] | 0.04 | 69.32 |
| | Climate | Boreal | 15 | | -0.20 [-0.87,0.46] | 0.55 | 80.27 |
| | | Temperate | 231 | | -0.20 [-0.46,0.07] | 0.14 | 92.26 |
| | | Sub-Tropical | 55 | | -0.60 [-1.1,-0.09] | 0.02 | 84.17 |
| | | Tropical | 88 | | 0.02 [-0.33,0.38] | 0.90 | 88.68 |
| | Metric | Abundance | 175 | | -0.12 [-0.32,0.08] | 0.25 | 79.08 |

| | | | | | |
|---|---|---|---|---|---|
| | Biomass | 55 | -0.12 [-0.39,0.15] | 0.37 | 62.05 |
| | Diversity | 159 | -0.29 [-0.49,-0.08] | 0.01 | 82.96 |
| Type | Fish+crayfish | 23 | 0.06 [-0.81,0.92] | 0.90 | 90.00 |
| | Macroinvertebrates | 366 | -0.21 [-0.40,-0.01] | 0.04 | 90.34 |
| Vegetation Change | Conversion | 131 | -0.20 [-0.50,0.09] | 0.17 | 83.35 |
| | Harvest | 152 | -0.01 [-0.31,0.30] | 0.97 | 92.41 |
| | Plantation | 88 | -0.45 [-0.83,-0.07] | 0.02 | 89.47 |
| | Restoration | 18 | -0.28 [-1.06,0.50] | 0.48 | 88.50 |
| Scale | Catchment | 83 | -0.25 [-0.59,0.09] | 0.15 | 90.37 |
| | Local | 144 | -0.02 [-0.34,0.31] | 0.93 | 88.54 |
| | Local+Catchment | 162 | -0.30 [-0.56,-0.04] | 0.02 | 87.54 |

**Table S3**. Effects of moderators tested in the analyses based on the dataset with only reported comparions (i.e., excluding imputed and estimated values). Test for heterogeneity between levels within moderators ($Q_M$), degrees of freedom (df) and p-values (significant differences among moderator levels exist if p-values < 0.05; see Figure S6) are shown.

| Moderator | Levels | $Q_M$ | df | p-value |
|---|---|---|---|---|
| All data | | 2550.84 | 747 | < 0.01 |
| Climate | 4 levels: boreal, temperate, subtropical, tropical | 4.36 | 3 | 0.22 |
| Type of vegetation change | 4 levels: restoration, plantation, harvest, conversion | 8.67 | 3 | 0.03 |
| Spatial scale | 3 levels: local+catchment, local, catchment | 2.65 | 2 | 0.27 |
| Metric | 4 levels: abundance, biomass, diversity, decomposition | 14.84 | 3 | < 0.01 |
| Trophic level | 4 levels: omnivores, shredders, microbes, detritus | 15.33 | 3 | < 0.01 |

**Table S4.** Effects of moderators tested for different trophic levels for the subset of only reported comparisons (i.e., excluding imputed and estimated values). Test for heterogeneity between levels within moderators ($Q_M$), degrees of freedom (df) and p-values (significant differences among moderator levels exist if p-values < 0.05; see Figure 7) are shown.

| Moderators | Levels | $Q_M$ | df | p-value |
|---|---|---|---|---|
| A. Detritus | | | | |
| All | | 1646.94 | 299 | < 0.01 |
| Climate | 4 levels: boreal, temperate, subtropical, tropical | 3.29 | 3 | 0.35 |
| Type of vegetation change | 4 levels: restoration, plantation, harvest, conversion | 13.30 | 3 | 0.004 |
| Spatial scale | 3 levels: local+catchment, local, catchment | 5.68 | 2 | 0.06 |
| Metric | 5 levels: abundance, biomass, total decomposition, shredder decomposition, microbial decomposition | 12.65 | 4 | 0.01 |
| Type of detritus | 5 levels: wood, reproductive, mixed, leaf, cotton strip | 11.39 | 4 | 0.02 |
| B. Microbes | | | | |
| All | | 64.18 | 33 | < 0.01 |
| Climate | 4 levels: boreal, temperate, subtropical, tropical | 0.48 | 3 | 0.92 |
| Type of vegetation change | 3 levels: restoration, plantation, harvest | 6.92 | 2 | 0.03 |
| Spatial scale | 3 levels: local+catchment, local, catchment | 1.34 | 2 | 0.51 |
| Metric | 2 levels: biomass, diversity | 0.37 | 1 | 0.55 |
| Type of microbes | 3 levels: microbes, fungi, bacteria | 0.53 | 2 | 0.77 |
| C. Shredders | | | | |
| All | | 383.28 | 192 | < 0.01 |
| Climate | 4 levels: boreal, temperate, subtropical, tropical | 8.37 | 3 | 0.04 |
| Type of vegetation change | 4 levels: restoration, plantation, harvest, conversion | 2.33 | 3 | 0.51 |
| Spatial scale | 3 levels: local+catchment, local, catchment | 1.48 | 2 | 0.48 |
| Metric | 2 levels: abundance, biomass, diversity | 0.80 | 2 | 0.67 |
| Type of substrate | 5 levels: detritus, benthic | 0.08 | 1 | 0.78 |
| D. Omnivores | | | | |
| All | | 447.74 | 226 | < 0.01 |
| Climate | 4 levels: boreal, temperate, subtropical, tropical | 6.69 | 3 | 0.08 |
| Type of vegetation change | 4 levels: restoration, plantation, harvest, conversion | 2.89 | 3 | 0.41 |
| Spatial scale | 3 levels: local+catchment, local, catchment | 0.93 | 2 | 0.63 |
| Metric | 2 levels: abundance, biomass, diversity | 4.75 | 2 | 0.09 |
| Type of omnivores | 5 levels: macroinvertebrates, fish+crayfish | 1.61 | 1 | 0.20 |

**Table S5**. Source, full reference, information on data extracted and excluded from the studies selected for inclusion, and main moderators.

| Study ID | Source | Full reference | Data extracted Reference | Altered | Data excluded | Climate | Type of vegetation change | Scale of vegetation change | Trophic levels | Metrics |
|---|---|---|---|---|---|---|---|---|---|---|
| 1 | (Abelho & Graça, 1996) | Abelho, M., & Graça, M. A. S. (1996). Effects of eucalyptus afforestation on leaf litter dynamics and macroinvertebrate community structure of streams in Central Portugal. *Hydrobiologia*, 324(3), 195–204. https://doi.org/10.1007/BF00016391 | Deciduous | Eucalyptus | Mixed land use | temperate | plantation | local | detritus, omnivore | biomass, decomposition, abundance, diversity |
| 2 | (Aguiar et al., 2018) | Aguiar, A. C. F., Neres-Lima, V., & Moulton, T. P. (2018). Relationships of shredders, leaf processing and organic matter along a canopy cover gradient in tropical streams. *Journal of Limnology*, 77(1), Article 1. https://doi.org/10.4081/jlimnol.2017.1684 | VAL | CAP | Other intermediate sites | tropical | conversion | local | detritus, shredder | biomass, decomposition |
| 3 | (Albertson et al., 2018) | Albertson, L. K., Ouellet, V., & Daniels, M. D. (2018). Impacts of stream riparian buffer land use on water temperature and food availability for fish. *Journal of Freshwater Ecology*, 33(1), 195–210. https://doi.org/10.1080/02705060.2017.1422558 | ChesLen | Taylor Run | Other intermediate sites | temperate | conversion | local | omnivore | abundance, biomass, diversity |
| 4 | (Astudillo et al., 2016) | Astudillo, M. R., Novelo-Gutiérrez, R., Vázquez, G., García-Franco, J. G., & Ramírez, A. (2016). Relationships between land cover, riparian vegetation, stream characteristics, and aquatic insects in cloud forest streams, Mexico. *Hydrobiologia*, 768(1), 167–181. https://doi.org/10.1007/s10750-015-2545-1 | Chivizcoyo | El Chorrito | Other intermediate sites | tropical | harvest | local+catchment | omnivore, shredder | abundance, diversity |
| 5 | (Bacca et al., 2023) | Bacca, J. C., Rampanelli Cararo, E., Alves Lima-Rezende, C., Tavares Martins, R., Macedo-Reis, L. E., Dal Magro, J., & De Souza Rezende, R. (2023). Land-use effects on aquatic macroinvertebrate diversity in subtropical highland grasslands streams. *Limnetica*, 42(2), 1. https://doi.org/10.23818/limn.42.16 | Chivizcoyo | GNRV; Silviculture | Grassland with riparian buffer | subtropical | harvest, conversion | local | detritus, omnivore | biomass, abundance, diversity |
| 6 | (Bärlocher et al., 2010) | Bärlocher, F., Helson, J. E., & Williams, D. D. (2010). Aquatic hyphomycete communities across a land-use gradient of Panamanian streams. *Fundamental and Applied Limnology*, 177(3), 209–221. https://doi.org/10.1127/1863-9135/2010/0177-0209 | Pristine | Rural | Urban | tropical | conversion | local | microbes | diversity |
| 8 | (Barrios et al., 2022) | Barrios, M., Burwood, M., Kröger, A., Calvo, C., Ríos-Touma, B., & Teixeira-de-Mello, F. (2022). Riparian cover buffers the effects of abiotic and biotic predictors of leaf decomposition in subtropical streams. *Aquatic Sciences*, 84(4), 55. https://doi.org/10.1007/s00027-022-00886-z | RFS | OCS | | subtropical | conversion | local | omnivore, shredder, detritus | abundance, decomposition |
| 9 | (Basaguren & Pozo, 1994) | Basaguren, A., & Pozo, J. (1994). Leaf litter processing of alder and eucalyptus in the Agüera stream system (Northern Spain). II: Macroinvertebrates associated. Leaf*Archiv für Hidrobiologie*, 132(1), 57–68. | S1 | S7 | SA, S9 | temperate | plantation | catchment | detritus | decomposition |
| 10 | (Benfield et al., 2001) | Benfield, E. F., Webster, J. R., Tank, J. L., & Hutchens, J. J. (2001). Long-Term Patterns in Leaf Breakdown in | Big Hurricane | Big Hurricane | Hugh White Creek and later | temperate | harvest | catchment | detritus | decomposition |

| | | | | | | | | | | |
|---|---|---|---|---|---|---|---|---|---|---|
| | | Streams in Response to Watershed Logging. *International Review of Hydrobiology*, 86(4–5), 467–474. https://doi.org/10.1002/1522-2632(200107)86:4/5<467::AID-IROH467>3.0.CO;2-1 | Branch before logging | Branche during logging | measurements with different leaf litter species | | | | | |
| 11 | (Bertaso et al., 2015) | Bertaso, T. R. N., Spies, M. R., Kotzian, C. B., & Flores, M. L. T. (2015). Effects of forest conversion on the assemblages' structure of aquatic insects in subtropical regions. *Revista Brasileira de Entomologia*, 59(1), 43–49. https://doi.org/10.1016/j.rbe.2015.02.005 | Forested | Converted | | subtropical | conversion | catchment | omnivore, shredder | abundance, diversity |
| 13 | (Burrows et al., 2014) | Burrows, R. M., Magierowski, R. H., Fellman, J. B., Clapcott, J. E., Munks, S. A., Roberts, S., Davies, P. E., & Barmuta, L. A. (2014). Variation in stream organic matter processing among years and benthic habitats in response to forest clearfelling. *Forest Ecology and Management*, *327*, 136–147. https://doi.org/10.1016/j.foreco.2014.04.041 | D, E, F, H, in April 2008 | A and B in April 2008 | Other dates, fine sediment sites | temperate | harvest | local+catchment | detritus | decomposition |
| 14 | (Burrows et al., 2012) | Burrows, R. M., Magierowski, R. H., Fellman, J. B., & Barmuta, L. A. (2012). Woody debris input and function in old-growth and clear-felled headwater streams. *Forest Ecology and Management*, 286, 73–80. https://doi.org/10.1016/j.foreco.2012.08.038 | Old growth forest | CBS-affected | | temperate | harvest | local+catchment | detritus | decomposition, abundance |
| 15 | (Burwood et al., 2021) | Burwood, M., Clemente, J., Meerhoff, M., Iglesias, C., Goyenola, G., Fosalba, C., Pacheco, J. P., & Teixeira De Mello, F. (2021). Macroinvertebrate communities and macrophyte decomposition could be affected by land use intensification in subtropical lowland streams. *Limnetica*, 40(2), 343–357. https://doi.org/10.23818/limn.40.23 | Extensive | Intensive | | subtropical | conversion | local+catchment | detritus, omnivore, shredder | decomposition, abundance, biomass, diversity |
| 16 | (Bylak et al., 2022) | Bylak, A., Kukuła, K., Ortyl, B., Hałoń, E., Demczyk, A., Janora-Hołyszko, K., Maternia, J., Szczurowski, Ł., & Ziobro, J. (2022). Small stream catchments in a developing city context: The importance of land cover changes on the ecological status of streams and the possibilities for providing ecosystem services. *Science of the Total Environment*, 815, 151974. https://doi.org/10.1016/j.scitotenv.2021.151974 | 1L | 3L | 2L | temperate | conversion | local+catchment | omnivore | diversity |
| 17 | (Carroll & Jackson, 2009) | Carroll, D., & Jackson, G. R. (2009). Observed relationships between urbanization and riparian cover, shredder abundance, and stream leaf litter standing crops. *Fundamental and Applied Limnology*, 213–225. https://doi.org/10.1127/1863-9135/2008/0173-0213 | Most covered | Least covered | Urban | temperate | conversion | local+catchment | shredder, detritus | abundance, diversity, biomass |
| 18 | (Casas et al., 2013) | Casas, J. J., Larrañaga, A., Menéndez, M., Pozo, J., Basaguren, A., Martínez, A., Pérez, J., González, J. M., Mollá, S., Casado, C., Descals, E., Roblas, N., López-González, J. A., & Valenzuela, J. L. (2013). Leaf litter decomposition of native and introduced tree species of contrasting quality in headwater streams: How does the regional setting matter? *Science of the Total Environment*, 458–460, 197–208. https://doi.org/10.1016/j.scitotenv.2013.04.004 | CC | SN | CLC, SG | temperate | plantation | local+catchment | shredder, detritus | abundance, decomposition, biomass |

| | | | | | | | | | | |
|---|---|---|---|---|---|---|---|---|---|---|
| 19 | (Casotti et al., 2015) | Casotti, C. G., Kiffer, W. P., Costa, L. C., Rangel, J. V., Casagrande, L. C., & Moretti, M. S. (2015). Assessing the importance of riparian zones conservation for leaf decomposition in streams. *Natureza & Conservação*, 13(2), 178–182. https://doi.org/10.1016/j.ncon.2015.11.011 | Banana | Norte | Luxemburgo and Macucuo | tropical | plantation | local | detritus, omnivore, shredder | decomposition, diversity, abundance |
| 21 | (de Mello Cionek et al., 2021) | de Mello Cionek, V., Fogaça, F. N. O., Moulton, T. P., Pazianoto, L. H. R., Landgraf, G. O., & Benedito, E. (2021). Influence of leaf miners and environmental quality on litter breakdown in tropical headwater streams. *Hydrobiologia*, 848(6), 1311–1331. https://doi.org/10.1007/s10750-021-04529-6 | 21 | 13; 24 | | tropical | conversion | local+catchment | detritus, microbes | decomposition, biomass |
| 22 | (Collier et al., 2004) | Collier, K. J., Smith, B. J., & Halliday, N. J. (2004). Colonization and use of pine wood versus native wood in New Zealand plantation forest streams: Implications for riparian management. *Aquatic Conservation: Marine and Freshwater Ecosystems*, 14(2), 179–199. https://doi.org/10.1002/aqc.599 | PE4 | PE2 | | temperate | harvest | catchment | microbes, omnivore | biomass, diversity, abundance |
| 23 | (Collier & Halliday, 2000) | Collier, K.J. & Halliday, J.N. (2000). Macroinvertebrate-wood associations during decay of plantation pine in New Zealand pumice-bed streams: stable habitat or trophic subsidy? J. North Am. Benthol. Soc., 19, 94–111. | 13 | 2 | Other sites | temperate | restoration | catchment | omnivore | diversity, abundance |
| 24 | (Cordero–Rivera et al., 2017) | Cordero–Rivera, A., Álvarez, A. M., & Álvarez, M. (2017). Eucalypt plantations reduce the diversity of macroinvertebrates in small forested streams. *Animal Biodiversity and Conservation*, 40(1), Article 1. https://doi.org/10.32800/abc.2017.40.0087 | Least Eucalyptus cover | Most Eucalyptus cover | Other sites | temperate | plantation | catchment | omnivore | diversity |
| 25 | (Cornejo et al., 2020) | Cornejo, A., Pérez, J., López-Rojo, N., Tonin, A. M., Rovira, D., Checa, B., Jaramillo, N., Correa, K., Villarreal, A., Villarreal, V., García, G., Pérez, E., Ríos González, T. A., Aguirre, Y., Correa-Araneda, F., & Boyero, L. (2020). Agriculture impairs stream ecosystem functioning in a tropical catchment. *Science of the Total Environment*, 745, 140950. https://doi.org/10.1016/j.scitotenv.2020.140950 | PA | BA | AA | tropical | conversion | local+catchment | detritus, omnivore, shredder | abundance, decomposition, diversity, biomass |
| 26 | (Chellaiah & Yule, 2019) | Chellaiah, D., & Yule, C. M. (2019). Litter quality influences bacterial communities more strongly than changes in riparian buffer quality in oil palm streams. *Aquatic Microbial Ecology*, 83(2), 167–179. https://doi.org/10.3354/ame01909 | NF | OPNB | OPF, OPOP | tropical | plantation | local+catchment | detritus, microbes | decomposition, diversity |
| 27 | (Chellaiah & Yule, 2018) | Chellaiah, D., & Yule, C. M. (2018). Litter decomposition is driven by microbes and is more influenced by litter quality than environmental conditions in oil palm streams with different riparian types. *Aquatic Sciences*, 80(4), 43. https://doi.org/10.1007/s00027-018-0595-y | NF | OPNB | OPF, OPOP | tropical | plantation | local+catchment | detritus | decomposition |
| 28 | (da Costa et al., 2020) | da Costa, I. D., Petry, A. C., & Mazzoni, R. (2020). Fish assemblages respond to forest cover in small Amazonian basins. *Limnologica*, 81, 125757. https://doi.org/10.1016/j.limno.2020.125757 | Forested | Deforested | | tropical | harvest | catchment | omnivore, detritus | diversity, abundance |

| | | | | | | | | | |
|---|---|---|---|---|---|---|---|---|---|
| 29 | (Lemes da Silva et al., 2020) | Lemes da Silva, A. L., Lemes, W. P., Andriotti, J., Petrucio, M. M., & Feio, M. J. (2020). Recent land-use changes affect stream ecosystem processes in a subtropical island in Brazil. *Austral Ecology*, 45(5), 644–658. https://doi.org/10.1111/aec.12879 | Forested | Rural | Urban | subtropical | conversion | catchment | detritus, shredder, omnivore | decomposition, abundance |
| 30 | (Danger & Robson, 2004) | Danger, A. R., & Robson, B. J. (2004). The effects of land use on leaf-litter processing by macroinvertebrates in an Australian temperate coastal stream. *Aquatic Sciences*, 66(3), 296–304. https://doi.org/10.1007/s00027-004-0718-5 | Forested | Pasture | Boundary | temperate | conversion | local+catchment | detritus, shredder | decomposition, abundance |
| 31 | (Daoust et al., 2019) | Daoust, K., Kreutzweiser, D. P., Guo, J., Creed, I. F., & Sibley, P. K. (2019). Climate-influenced catchment hydrology overrides forest management effects on stream benthic macroinvertebrates in a northern hardwood forest. *Forest Ecology and Management*, 452, 117540. https://doi.org/10.1016/j.foreco.2019.117540 | Forested | Harvested | | temperate | harvest | catchment | omnivore | diversity |
| 33 | (de Nadaï-Monoury et al., 2014) | de Nadaï-Monoury, E., Gilbert, F., & Lecerf, A. (2014). Forest canopy cover determines invertebrate diversity and ecosystem process rates in depositional zones of headwater streams. *Freshwater Biology*, 59(7), 1532–1545. https://doi.org/10.1111/fwb.12364 | Forested | Logged | | temperate | harvest | local+catchment | detritus, omnivore, shredder | decomposition, diversity, abundance |
| 34 | (Delong & Brusven, 1994) | Delong, M. D., & Brusven, M. A. (1994). Allochthonous input of organic matter from different riparian habitats of an agriculturally impacted stream. *Environmental Management*, 18(1), 59–71. https://doi.org/10.1007/BF02393750 | 1,6 | 3,7 | Other sites | temperate | conversion | local+catchment | detritus | biomass |
| 35 | (Dézerald et al., 2014) | Dézerald, O., Talaga, S., Leroy, C., Carrias, J.-F., Corbara, B., Dejean, A., & Céréghino, R. (2014). Environmental determinants of macroinvertebrate diversity in small water bodies: Insights from tank-bromeliads. *Hydrobiologia*, 723(1), 77–86. https://doi.org/10.1007/s10750-013-1464-2 | Primary forest | Citrus plantation | Other sites | tropical | plantation | local | omnivore, shredder | diversity |
| 36 | (Díez et al., 2002) | Díez, J., Elosegi, A., Chauvet, E., & Pozo, J. (2002). Breakdown of wood in the Agüera stream. *Freshwater Biology*, 47(11), 2205–2215. https://doi.org/10.1046/j.1365-2427.2002.00965.x | D1 | E1 | Other sites | temperate | plantation | local+catchment | detritus, microbes | decomposition, biomass |
| 37 | (dos Santos et al., 2015) | dos Santos, F. B., Ferreira, F. C., & Esteves, K. E. (2015). Assessing the importance of the riparian zone for stream fish communities in a sugarcane dominated landscape (Piracicaba River Basin, Southeast Brazil). *Environmental Biology of Fishes*, 98(8), 1895–1912. https://doi.org/10.1007/s10641-015-0406-4 | Forest | Sugarcane | | subtropical | conversion | local+catchment | omnivore | abundance, diversity, biomass |
| 38 | (Dudgeon, 2006) | Dudgeon, D. (2006). The impacts of human disturbance on stream benthic invertebrates and their drift in North Sulawesi, Indonesia. *Freshwater Biology*, 51(9), 1710–1729. https://doi.org/10.1111/j.1365-2427.2006.01596.x | WS | AS | Other sites | tropical | conversion | local | omnivore | diversity, abundance |
| 39 | (Dudley et al., 2021) | Dudley, M. P., Solomon, K., Wenger, S., Jackson, C. R., Freeman, M., Elliott, K. J., Miniat, C. F., & Pringle, C. M. (2021). Do crayfish affect stream ecosystem response to | Cut+Burn PreYR 2014 | Cut+Burn Burn YR 2016 | Other sites and timepoints | temperate | restoration | local | omnivore, detritus | abundance, decomposition |

| | | | | | | | | | | |
|---|---|---|---|---|---|---|---|---|---|---|
| | | riparian vegetation removal? *Freshwater Biology*, 66(7), 1423–1435. https://doi.org/10.1111/fwb.13728 | | | | | | | | |
| 40 | (Ebling et al., 2024) | Ebling, L. A., Pastore, B. L., Biasi, C., Hepp, L. U., & Restello, R. M. (2024). Does environmental variability in Atlantic Forest streams affect aquatic hyphomycete and invertebrate assemblages associated with leaf litter? *Hydrobiologia*, 851(7), 1761–1777. https://doi.org/10.1007/s10750-023-05415-z | S3 | S4 | Other sites | subtropical | conversion | catchment | detritus, microbes, omnivore, shredder | decomposition, diversity, abundance |
| 41 | (Elosegi et al., 2018) | Elosegi, A., Nicolás, A., & Richardson, J. S. (2018). Priming of leaf litter decomposition by algae seems of minor importance in natural streams during autumn. *PLOS ONE*, 13(9), e0200180. https://doi.org/10.1371/journal.pone.0200180 | Closed canopy; Forest | Open canopy; Agriculture | | temperate | conversion | local, catchment | detritus | decomposition |
| 42 | (Elosegi et al., 2007) | Elosegi, A., Díez, J., & Pozo, J. (2007). Contribution of dead wood to the carbon flux in forested streams. *Earth Surface Processes and Landforms*, 32(8), 1219–1228. https://doi.org/10.1002/esp.1549 | Salderrey | Jergueron | | temperate | plantation | catchment | detritus | decomposition |
| 43 | (Ely & Wallace, 2010) | Ely, D. T., & Wallace, J. B. (2010). Long-term functional group recovery of lotic macroinvertebrates from logging disturbance. *Canadian Journal of Fisheries and Aquatic Sciences*, 67(7), 1126–1134. https://doi.org/10.1139/F10-045 | Reference | Disturbed | | temperate | harvest | catchment | shredder, omnivore, detritus | abundance, biomass |
| 44 | (Emilson et al., 2016) | Emilson, C. E., Kreutzweiser, D. P., Gunn, J. M., & Mykytczuk, N. C. S. (2016). Effects of land use on the structure and function of leaf-litter microbial communities in boreal streams. *Freshwater Biology*, 61(7), 1049–1061. https://doi.org/10.1111/fwb.12765 | Undisturbed | Logged | Sites with other types of disturbances | boreal | harvest | catchment | detritus, microbes | decomposition, biomass, diversity |
| 45 | (Encalada et al., 2010) | Encalada, A. C., Calles, J., Ferreira, V., Canhoto, C. M., & Graça, M. A. S. (2010). Riparian land use and the relationship between the benthos and litter decomposition in tropical montane streams. *Freshwater Biology*, 55(8), 1719–1733. https://doi.org/10.1111/j.1365-2427.2010.02406.x | Forest | Pasture | | tropical | conversion | local | detritus, microbes, omnivore, shredder | decomposition, biomass, diversity, abundance |
| 46 | (Erdozain et al., 2021) | Erdozain, M., Kidd, K. A., Emilson, E. J. S., Capell, S. S., Luu, T., Kreutzweiser, D. P., & Gray, M. A. (2021). Forest management impacts on stream integrity at varying intensities and spatial scales: Do biological effects accumulate spatially? *Science of the Total Environment*, 763, 144043. https://doi.org/10.1016/j.scitotenv.2020.144043 | NBR | NBI | | temperate | harvest | local+catchment | omnivore, shredder, detritus | abundance, diversity, decomposition |
| 47 | (Erdozain et al., 2018) | Erdozain, M., Kidd, K., Kreutzweiser, D., & Sibley, P. (2018). Linking stream ecosystem integrity to catchment and reach conditions in an intensively managed forest landscape. *Ecosphere*, 9(5), e02278. https://doi.org/10.1002/ecs2.2278 | Reference | Harvested | | temperate | harvest | local+catchment | detritus, omnivore, shredder | decomposition, abundance, diversity |
| 49 | (Espinoza-Toledo et al., 2021) | Espinoza-Toledo, A., Mendoza-Carranza, M., Castillo, M. M., Barba-Macías, E., & Capps, K. A. (2021). Taxonomic and functional responses of macroinvertebrates to riparian forest conversion in tropical streams. *Science of the Total* | Forest | Agriculture | | tropical | conversion | local+catchment | detritus, omnivore | biomass, abundance, diversity |

| | | | | | | | | | | |
|---|---|---|---|---|---|---|---|---|---|---|
| | | *Environment*, 757, 143972. https://doi.org/10.1016/j.scitotenv.2020.143972 | | | | | | | | |
| 50 | (Evangelista et al., 2014) | Evangelista, C., Boiche, A., Lecerf, A., & Cucherousset, J. (2014). Ecological opportunities and intraspecific competition alter trophic niche specialization in an opportunistic stream predator. *Journal of Animal Ecology*, 83(5), 1025–1034. https://doi.org/10.1111/1365-2656.12208 | Most covered | Least covered | Other sites | temperate | conversion | local | omnivore, shredder | biomass |
| 51 | (Fargen et al., 2015) | Fargen, C., Emery, S. M., & Carreiro, M. M. (2015). Influence of *Lonicera maackii* invasion on leaf litter decomposition and macroinvertebrate communities in an urban stream. *Natural Areas Journal*, 35(3), 392–403. https://doi.org/10.3375/043.035.0303 | Removed | Invaded | | temperate | restoration | local | detritus, shredder, omnivore | decomposition, abundance |
| 52 | (Fasching et al., 2020) | Fasching, C., Akotoye, C., Bižić, M., Fonvielle, J., Ionescu, D., Mathavarajah, S., Zoccarato, L., Walsh, D. A., Grossart, H.-P., & Xenopoulos, M. A. (2020). Linking stream microbial community functional genes to dissolved organic matter and inorganic nutrients. *Limnology and Oceanography*, 65(S1), S71–S87. https://doi.org/10.1002/lno.11356 | Forest | Agriculture | | temperate | conversion | local+catchment | microbes | diversity |
| 53 | (Ferreira et al., 2019) | Ferreira, V., Boyero, L., Calvo, C., Correa, F., Figueroa, R., Gonçalves, J. F., Goyenola, G., Graça, M. A. S., Hepp, L. U., Kariuki, S., López-Rodríguez, A., Mazzeo, N., M'Erimba, C., Monroy, S., Peil, A., Pozo, J., Rezende, R., & Teixeira-de-Mello, F. (2019). A global assessment of the effects of *Eucalyptus* plantations on stream ecosystem functioning. *Ecosystems*, 22(3), 629–642. https://doi.org/10.1007/s10021-018-0292-7 | Forest | Plantation | | temperate, tropical | plantation | local | detritus | decomposition |
| 54 | (Ferreira et al., 2017) | Ferreira, V., Faustino, H., Raposeiro, P. M., & Gonçalves, V. (2017). Replacement of native forests by conifer plantations affects fungal decomposer community structure but not litter decomposition in Atlantic island streams. *Forest Ecology and Management*, 389, 323–330. https://doi.org/10.1016/j.foreco.2017.01.004 | Forest | Plantation | | temperate | plantation | local | detritus, microbes | decomposition, diversity |
| 55 | (Ferreira et al., 2015) | Ferreira, V., Larrañaga, A., Gulis, V., Basaguren, A., Elosegi, A., Graça, M. A. S., & Pozo, J. (2015). The effects of eucalypt plantations on plant litter decomposition and macroinvertebrate communities in Iberian streams. *Forest Ecology and Management*, 335, 129–138. https://doi.org/10.1016/j.foreco.2014.09.013 | Forest | Plantation | | temperate | plantation | local | detritus, omnivore, shredder | decomposition, diversity, abundance |
| 56 | (Ferreira et al., 2015) | Ferreira, W. R., Ligeiro, R., Macedo, D. R., Hughes, R. M., Kaufmann, P. R., Oliveira, L. G., & Callisto, M. (2015). Is the diet of a typical shredder related to the physical habitat of headwater streams in the Brazilian Cerrado? *Annales de Limnologie*, 51(2), 115–127. https://doi.org/10.1051/limn/2015004 | Upper Araguari River Basin | Upper Sao Francisco River Basin | | tropical | conversion | local+catchment | omnivore, shredder | abundance, diversity |
| 57 | (Ferreira et al., 2006) | Ferreira, V., Elosegi, A., Gulis, V., Pozo, J., & Graça, M. A. S. (2006). Eucalyptus plantations affect fungal communities associated with leaf-litter decomposition in | Forest | Plantation | | temperate | plantation | local | microbes | biomass, diversity |

| | | | | | | | | | | |
|---|---|---|---|---|---|---|---|---|---|---|
| | | Iberian streams. *Archiv für Hydrobiologie*, 166, 467–490. https://doi.org/10.1127/0003-9136/2006/0166-0467 | | | | | | | | |
| 59 | (Valente-Neto et al., 2015) | Valente-Neto, F., Koroiva, R., Fonseca-Gessner, A. A., & Roque, F. de O. (2015). The effect of riparian deforestation on macroinvertebrates associated with submerged woody debris. *Aquatic Ecology*, *49*(1), 115–125. https://doi.org/10.1007/s10452-015-9510-y | Forest | Pasture | Agriculture | tropical | conversion | local+catchment | omnivore | abundance, diversity |
| 60 | (Four et al., 2017) | Four, B., Arce, E., Danger, M., Gaillard, J., Thomas, M., & Banas, D. (2017). Catchment land use-dependent effects of barrage fishponds on the functioning of headwater streams. *Environmental Science and Pollution Research*, 24(6), 5452–5468. https://doi.org/10.1007/s11356-016-8273-x | Forest | Agriculture | Downstream of fish pond sites | temperate | conversion | local+catchment | detritus, microbes, shredder, omnivore | decomposition, biomass, abundance, diversity |
| 61 | (Fu et al., 2016) | Fu, L., Jiang, Y., Ding, J., Liu, Q., Peng, Q.-Z., & Kang, M.-Y. (2016). Impacts of land use and environmental factors on macroinvertebrate functional feeding groups in the Dongjiang River basin, southeast China. *Journal of Freshwater Ecology*, 31(1), 21–35. https://doi.org/10.1080/02705060.2015.1017847 | Forest | Agriculture | | subtropical | conversion | catchment | shredder | abundance, diversity |
| 62 | (Fugère et al., 2018) | Fugère, V., Jacobsen, D., Finestone, E. H., & Chapman, L. J. (2018). Ecosystem structure and function of afrotropical streams with contrasting land use. *Freshwater Biology*, 63(12), 1498–1513. https://doi.org/10.1111/fwb.13178 | Forest | Farmland | | tropical | conversion | local+catchment | detritus, shredder | decomposition, biomass, abundance |
| 63 | (Giling et al., 2015) | Giling, D. P., Nally, R. M., & Thompson, R. M. (2015). How sensitive are invertebrates to riparian-zone replanting in stream ecosystems? *Marine and Freshwater Research*, 67(10), 1500–1511. https://doi.org/10.1071/MF14360 | Replanted | Untreated | Reference | subtropical | restoration | local+catchment | omnivore, shredder | abundance, diversity |
| 64 | (Godoy et al., 2016) | Godoy, B. S., Simião-Ferreira, J., Lodi, S., & Oliveira, L. G. (2016). Functional process zones characterizing aquatic insect communities in streams of the Brazilian Cerrado. *Neotropical Entomology*, 45(2), 159–169. https://doi.org/10.1007/s13744-015-0352-z | FPZ3 | FPZ2 | Other sites | tropical | harvest | local | omnivore, shredder | diversity, abundance |
| 66 | (Goodman et al., 2006) | Goodman, K. J., Hershey, A. E., & Fortino, K. (2006). The effect of forest type on benthic macroinvertebrate structure and ecological function in a pine plantation in the North Carolina Piedmont. *Hydrobiologia*, 559(1), 305–318. https://doi.org/10.1007/s10750-005-0990-y | Natural hardwood forest | Clea-cut; Pine forestry | | temperate | harvest, plantation | local+catchment | detritus, omnivore | biomass, decomposition, diversity |
| 67 | (Goss et al., 2014) | Goss, C. W., Goebel, P. C., & Sullivan, S. M. P. (2014). Shifts in attributes along agriculture-forest transitions of two streams in central Ohio, USA. *Agriculture, Ecosystems & Environment*, 197, 106–117. https://doi.org/10.1016/j.agee.2014.07.026 | Inside forest | Outside forest | | temperate | conversion | local | shredder, detritus | abundance, decomposition |
| 68 | (Göthe et al., 2009) | Göthe, E., Lepori, F., & Malmqvist, B. (2009). Forestry affects food webs in northern Swedish coastal streams. *Fundamental and Applied Limnology*, 175(4), 281–294. https://doi.org/10.1127/1863-9135/2009/0175-0281 | Old-growth forest | Clear-cut forest | | boreal | harvest | local+catchment | omnivore, detritus, shredder | biomass, abundance, diversity |
| 69 | (Guevara et al., 2018) | Guevara, G., Godoy, R., & Franco, M. (2018). Linking riparian forest harvest to benthic macroinvertebrate communities in Andean headwater streams in southern | Unthinned forest | Thinned forest | | tropical | harvest | local+catchment | detritus, shredder, omnivore | biomass, decomposition, |

| | | | | | | | | | | |
|---|---|---|---|---|---|---|---|---|---|---|
| | | Chile. *Limnologica*, 68, 105–114. https://doi.org/10.1016/j.limno.2017.07.007 | | | | | | | | abundance, diversity |
| 70 | (Guevara et al., 2015) | Guevara, G., Godoy, R., Boeckx, P., Jara, C., & Oyarzún, C. (2015). Effects of riparian forest management on Chilean mountain in-stream characteristics. *Ecohydrology & Hydrobiology*, 15(3), 160–170. https://doi.org/10.1016/j.ecohyd.2015.07.003 | Forest | Harvested | | tropical | harvest | catchment | detritus | biomass, decomposition |
| 71 | (Hagen et al., 2006) | Hagen, E. M., Webster, J. R., & Benfield, E. F. (2006). Are leaf breakdown rates a useful measure of stream integrity along an agricultural landuse gradient? *Journal of the North American Benthological Society*, 25(2), 330–343. https://doi.org/10.1899/0887-3593(2006)25[330:ALBRAU]2.0.CO;2 | Forest | Agriculture | Intermediate sites | temperate | conversion | local+catchment | detritus, omnivore, shredder | decomposition, abundance, diversity |
| 72 | (Hebert et al., 2023) | Hebert, T. A., Kuehn, K. A., & Halvorson, H. M. (2023). Land use differentially alters microbial interactions and detritivore feeding during leaf decomposition in headwater streams. *Freshwater Biology*, 68(8), 1386–1399. https://doi.org/10.1111/fwb.14111 | Forest | Agriculture | Sites with nutrient additions | temperate | conversion | catchment | microbes, detritus | biomass, decomposition |
| 73 | (Hisabae et al., 2011) | Hisabae, M., Sone, S., & Inoue, M. (2011). Breakdown and macroinvertebrate colonization of needle and leaf litter in conifer plantation streams in Shikoku, southwestern Japan. *Journal of Forest Research*, 16(2), 108–115. https://doi.org/10.1007/s10310-010-0210-0 | Mixed deciduous forest | Conifer plantations | | temperate | plantation | local | detritus, omnivore | decomposition, abundance |
| 74 | (Hladyz et al., 2010) | Hladyz, S., Tiegs, S. D., Gessner, M. O., Giller, P. S., Rîşnoveanu, G., Preda, E., Nistorescu, M., Schindler, M., & Woodward, G. (2010). Leaf-litter breakdown in pasture and deciduous woodland streams: A comparison among three European regions. *Freshwater Biology*, 55(9), 1916–1929. https://doi.org/10.1111/j.1365-2427.2010.02426.x | Woodland | Pasture | | temperate | conversion | local+catchment | detritus | decomposition |
| 75 | (Houghton et al., 2011) | Houghton, D. C., Berry, E. A., Gilchrist, A., Thompson, J., & Nussbaum, M. A. (2011). Biological changes along the continuum of an agricultural stream: Influence of a small terrestrial preserve and use of adult caddisflies in biomonitoring. *Journal of Freshwater Ecology*, 26(3), 381–397. https://doi.org/10.1080/02705060.2011.563513 | Site 4 | Site 5 | Other sites | temperate | conversion | local | omnivore | abundance, diversity |
| 76 | (Huryn et al., 2002) | Huryn, A. D., Huryn, V. M. B., Arbuckle, C. J., & Tsomides, L. (2002). Catchment land-use, macroinvertebrates and detritus processing in headwater streams: Taxonomic richness versus function. *Freshwater Biology*, 47(3), 401–415. https://doi.org/10.1046/j.1365-2427.2002.00812.x | Forested | Agriculture | Urban and Wetland sites | temperate | conversion | local+catchment | detritus, omnivore, shredder | decomposition, diversity, biomass |
| 77 | (Illyová et al., 2011) | Illyová, M., Beracko, P., & Krno, I. (2011). Influence of land use on hyporheos in catchment streams of the Velka Fatra Mts. *Biologia*, 66(2), 320–327. https://doi.org/10.2478/s11756-011-0018-1 | Forest | Agriculture | | temperate | conversion | catchment | omnivore | abundance |
| 79 | (Iñiguez-Armijos et al., 2018) | Iñiguez-Armijos, C., Hampel, H., & Breuer, L. (2018). Land-use effects on structural and functional composition of benthic and leaf-associated macroinvertebrates in four | Forest | Pasture | | tropical | conversion | local+catchment | omnivore, shredder | abundance, diversity |

| | | | | | | | | | | |
|---|---|---|---|---|---|---|---|---|---|---|
| | | Andean streams. *Aquatic Ecology*, 52(1), 77–92. https://doi.org/10.1007/s10452-017-9646-z | | | | | | | | |
| 80 | (Iñiguez-Armijos et al., 2016) | Iñiguez-Armijos, C., Rausche, S., Cueva, A., Sánchez-Rodríguez, A., Espinosa, C., & Breuer, L. (2016). Shifts in leaf litter breakdown along a forest–pasture–urban gradient in Andean streams. *Ecology and Evolution*, 6(14), 4849–4865. https://doi.org/10.1002/ece3.2257 | Forest | Pasture | Urban | tropical | conversion | local | detritus, microbes | decomposition, biomass |
| 81 | (Zúñiga-Sarango et al., 2020) | Zúñiga-Sarango, W., Gaona, F. P., Reyes-Castillo, V., & Iñiguez-Armijos, C. (2020). Disrupting the biodiversity–ecosystem function relationship: Response of shredders and leaf breakdown to urbanization in Andean streams. *Frontiers in Ecology and Evolution*, 8. https://doi.org/10.3389/fevo.2020.592404 | Forest | Pasture | Urban and mixed sites | tropical | conversion | local+catchment | shredder, detritus | abundance, diversity, decomposition |
| 82 | (Inoue et al., 2012) | Inoue, M., Shinotou, S., Maruo, Y., & Miyake, Y. (2012). Input, retention, and invertebrate colonization of allochthonous litter in streams bordered by deciduous broadleaved forest, a conifer plantation, and a clear-cut site in southwestern Japan. *Limnology*, 13(2), 207–219. https://doi.org/10.1007/s10201-011-0369-x | Forest | Plantation, Clear-cut | | temperate | plantation, harvest | local+catchment | detritus, omnivore, shredder | biomass, abundance |
| 83 | (Ishikawa et al., 2016) | Ishikawa, N. F., Togashi, H., Kato, Y., Yoshimura, M., Kohmatsu, Y., Yoshimizu, C., Ogawa, N. O., Ohte, N., Tokuchi, N., Ohkouchi, N., & Tayasu, I. (2016). Terrestrial-aquatic linkage in stream food webs along a forest chronosequence: Multi-isotopic evidence. *Ecology*, 97(5), 1146–1158. https://doi.org/10.1890/15-1133.1 | KU | S34 | other intermediate sites | subtropical | harvest | local+catchment | omnivore, shredder | abundance, diversity, biomass |
| 84 | (Jinggut et al., 2012) | Jinggut, T., Yule, C. M., & Boyero, L. (2012). Stream ecosystem integrity is impaired by logging and shifting agriculture in a global megadiversity center (Sarawak, Borneo). *Science of the Total Environment*, 437, 83–90. https://doi.org/10.1016/j.scitotenv.2012.07.062 | Pristine | Farmed, Logged | | tropical | conversion, harvest | local | detritus, shredder | decomposition, abundance, diversity, biomass |
| 85 | (Jyväsjärvi et al., 2020) | Jyväsjärvi, J., Koivunen, I., & Muotka, T. (2020). Does the buffer width matter: Testing the effectiveness of forest certificates in the protection of headwater stream ecosystems. *Forest Ecology and Management*, 478, 118532. https://doi.org/10.1016/j.foreco.2020.118532 | Reference | Narrow Buffer | Wide Buffer | boreal | harvest | local+catchment | detritus, omnivore | biomass, decomposition, diversity |
| 86 | (Kadeka et al., 2021) | Kadeka, E. C., Masese, F. O., Lusega, D. M., Sitati, A., Kondowe, B. N., & Chirwa, E. R. (2021). No Difference in instream decomposition among upland agricultural and forested streams in Kenya. *Frontiers in Environmental Science*, 9. https://doi.org/10.3389/fenvs.2021.794525 | Forest | Agriculture | | tropical | conversion | catchment | detritus, omnivore, shredder | biomass, abundance, diversity, decomposition |
| 87 | (Kaylor & Warren, 2018) | Kaylor, M. J., & Warren, D. R. (2018). Canopy closure after four decades of postlogging riparian forest regeneration reduces cutthroat trout biomass in headwater streams through bottom-up pathways. *Canadian Journal of Fisheries and Aquatic Sciences*, 75(4), 513–524. https://doi.org/10.1139/cjfas-2016-0519 | Forest | Harvested | | temperate | harvest | local | detritus | biomass |
| 88 | (Kiffer Jr et al., 2018) | Kiffer Jr, W., Giuberti, T., Victor Serpa, K., Mendes, F., & Moretti, M. (2018). Do changes in riparian zones affect periphyton growth and invertebrate colonization on rocky | Pau Amarelo | Luxemburgo | Macuco | tropical | plantation | local+catchment | omnivore | abundance, biomass, diversity |

| | | | | | | | | | | |
|---|---|---|---|---|---|---|---|---|---|---|
| | | substrates in Atlantic Forest streams? *Iheringia Série Zoologia*, 108, 1–10. https://doi.org/10.1590/1678-4766e2018014 | | | | | | | | |
| 89 | (Kiffney & Richardson, 2010) | Kiffney, P. M., & Richardson, J. S. (2010). Organic matter inputs into headwater streams of southwestern British Columbia as a function of riparian reserves and time since harvesting. *Forest Ecology and Management*, 260(11), 1931–1942. https://doi.org/10.1016/j.foreco.2010.08.016 | Control | Clear-cut | | temperate | harvest | local+catchment | detritus | biomass |
| 90 | (Kominoski et al., 2011) | Kominoski, J. S., Marczak, L. B., & Richardson, J. S. (2011). Riparian forest composition affects stream litter decomposition despite similar microbial and invertebrate communities. *Ecology*, 92(1), 151–159. https://doi.org/10.1890/10-0028.1 | Deciduous forest | Conifer plantation | Mixed | temperate | harvest | local | detritus, omnivore, microbes | decomposition, biomass, abundance, diversity |
| 91 | (Kreutzweiser et al., 2008) | Kreutzweiser, D. P., Good, K. P., Capell, S. S., & Holmes, S. B. (2008). Leaf-litter decomposition and macroinvertebrate communities in boreal forest streams linked to upland logging disturbance. *Journal of the North American Benthological Society*, 27(1), 1–15. https://doi.org/10.1899/07-034R.1 | Forested | Logged | | boreal | harvest | local+catchment | detritus, omnivore, shredder | decomposition, abundance, diversity |
| 92 | (Laćan et al., 2010) | Laćan, I., Resh, V. H., & McBRIDE, J. R. (2010). Similar breakdown rates and benthic macroinvertebrate assemblages on native and *Eucalyptus globulus* leaf litter in Californian streams. *Freshwater Biology*, 55(4), 739–752. https://doi.org/10.1111/j.1365-2427.2009.02312.x | Native forest | Eucalyptus plantation | | temperate | plantation | local | detritus, omnivore | biomass, abundance, diversity |
| 93 | (Lamberti & Berg, 1995) | Lamberti, G. A., & Berg, M. B. (1995). Invertebrates and other benthic features as indicators of environmental change in Juday Creek, Indiana. *Natural Areas Journal*, *15*(3), 249–258. | Woodland | Agriculture | Urban | temperate | conversion | local+catchment | detritus | biomass, abundance |
| 94 | (Larrañaga et al., 2023) | Larrañaga, A., Perkins, D. M., Basaguren, A., Larrañaga, S., Pozo, J., & Montoya, J. M. (2023). Land use drives detritivore size structure and decomposition through shifts in resource quality and quantity. *Science of the Total Environment*, *892*, 164552. https://doi.org/10.1016/j.scitotenv.2023.164552 | Native Forest | Eucalyptus plantation | | temperate | plantation | local+catchment | omnivore, shredder, detritus | abundance, biomass, diversity, decomposition |
| 98 | (Lecerf et al., 2012) | Lecerf, A., Baudoin, J.-M., Besson, A. A., Lamothe, S., & Lagrue, C. (2012). Is smaller necessarily better? Effects of small-scale forest harvesting on stream ecosystems. *Annales de Limnologie*, *48*(4), 401–409. https://doi.org/10.1051/limn/2012028 | Mature forest | Harvested | | temperate | harvest | local+catchment | omnivore, detritus | abundance, decomposition |
| 99 | (Lecerf & Richardson, 2010) | Lecerf, A., & Richardson, J. S. (2010). Litter decomposition can detect effects of high and moderate levels of forest disturbance on stream condition. *Forest Ecology and Management*, *259*(12), 2433–2443. https://doi.org/10.1016/j.foreco.2010.03.022 | Forest | No reserve | Intermediate sites | temperate | harvest | local+catchment | detritus, shredder, microbes | decomposition, abundance, diversity, biomass |
| 100 | (Lecerf & Chauvet, 2008) | Lecerf, A., & Chauvet, E. (2008). Diversity and functions of leaf-decaying fungi in human-altered streams. *Freshwater Biology*, 53(8), 1658–1672. https://doi.org/10.1111/j.1365-2427.2008.01986.x | Control | Impact | Eutrophication and Mine pollution sites | temperate | harvest | catchment | detritus, microbes | decomposition, biomass, diversity |

| | | | | | | | | | | |
|---|---|---|---|---|---|---|---|---|---|---|
| 102 | (Link et al., 2022) | Link, M., Schreiner, V. C., Graf, N., Szöcs, E., Bundschuh, M., Battes, K. P., Cîmpean, M., Sures, B., Grabner, D., Buse, J., & Schäfer, R. B. (2022). Pesticide effects on macroinvertebrates and leaf litter decomposition in areas with traditional agriculture. *Science of the Total Environment*, 828, 154549. https://doi.org/10.1016/j.scitotenv.2022.154549 | With forest upstream | Without forest upstream | | temperate | conversion | local+catchment | omnivore, shredder, detritus | diversity, abundance, decomposition |
| 103 | (Lopes et al., 2015) | Lopes, M. P., Martins, R. T., Silveira, L. S., & Alves, R. G. (2015). The leaf breakdown of *Picramnia sellowii* (Picramniales: Picramniaceae) as index of anthropic disturbances in tropical streams. *Brazilian Journal of Biology*, *75*, 846–853. https://doi.org/10.1590/1519-6984.00414 | Forest | Impaired | | tropical | harvest | local | detritus, microbes, shredder | decomposition, biomass, abundance |
| 104 | (Lorion & Kennedy, 2009) | Lorion, C. M., & Kennedy, B. P. (2009). Riparian forest buffers mitigate the effects of deforestation on fish assemblages in tropical headwater streams. *Ecological Applications*, *19*(2), 468–479. https://doi.org/10.1890/08-0050.1 | Forest | Pasture | Intermediate sites | tropical | harvest | local+catchment | omnivore | abundance, biomass, diversity |
| 105 | (Lubanga et al., 2021) | Lubanga, H. L., Manyala, J. O., Sitati, A., Yegon, M. J., & Masese, F. O. (2021). Spatial variability in water quality and macroinvertebrate assemblages across a disturbance gradient in the Mara River Basin, Kenya. *Ecohydrology & Hydrobiology*, *21*(21), 718–730. https://doi.org/10.1016/j.ecohyd.2021.03.001 | Undisturbed | Disturbed | Moderately Disturbed | tropical | conversion | local+catchment | omnivore, shredder | diversity, abundance |
| 106 | (Mancuso et al., 2023) | Mancuso, J., Tank, J. L., Mahl, U. H., Vincent, A., & Tiegs, S. D. (2023). Monthly variation in organic-matter decomposition in agricultural stream and riparian ecosystems. *Aquatic Sciences*, *85*(3), 83. https://doi.org/10.1007/s00027-023-00975-7 | Forested | Agriculture | Intermediate sites | temperate | conversion | catchment | detritus | decomposition |
| 110 | (Maridet et al., 1998) | Maridet, L., Wasson, J.-G., Philippe, M., Amoros, C., & Naiman, R. (1998). Trophic structure of three streams with contrasting riparian vegetation and geomorphology. *Archiv für Hydrobiologie*, 61–85. https://doi.org/10.1127/archiv-hydrobiol/144/1998/61 | Vianon | Ozange, Triouzone | | temperate | harvest | catchment, local+catchment | omnivore, shredder | diversity, abundance, biomass |
| 111 | (Márquez et al., 2017) | Márquez, J. A., Principe, R. E., Cibils Martina, L., & Albariño, R. J. (2017). Pine needle litter acts as habitat but not as food source for stream invertebrates. *International Review of Hydrobiology*, 102(1–2), 29–37. https://doi.org/10.1002/iroh.201601856 | Grassland | Afforested | | subtropical | plantation | local+catchment | detritus, omnivore, shredder | decomposition, diversity, abundance |
| 112 | (Martínez et al., 2013) | Martínez, A., Larrañaga, A., Pérez, J., Descals, E., Basaguren, A., & Pozo, J. (2013). Effects of pine plantations on structural and functional attributes of forested streams. *Forest Ecology and Management*, 310, 147–155. https://doi.org/10.1016/j.foreco.2013.08.024 | Forest | Plantation | | temperate | plantation | catchment | detritus, omnivore, shredder, microbes | decomposition, biomass, diversity, abundance |
| 113 | (Masese et al., 2014) | Masese, F. O., Kitaka, N., Kipkemboi, J., Gettel, G. M., Irvine, K., & McClain, M. E. (2014). Litter processing and shredder distribution as indicators of riparian and catchment influences on ecological health of tropical | Forest | Agriculture | Mixed | tropical | conversion | local+catchment | detritus, omnivore, shredder | biomass, diversity, abundance, decomposition |

| | | | | | | | | | | |
|---|---|---|---|---|---|---|---|---|---|---|
| | | streams. *Ecological Indicators*, 46, 23–37. https://doi.org/10.1016/j.ecolind.2014.05.032 | | | | | | | | |
| 114 | (Mckie & Malmqvist, 2009) | Mckie, B. G., & Malmqvist, B. (2009). Assessing ecosystem functioning in streams affected by forest management: Increased leaf decomposition occurs without changes to the composition of benthic assemblages. *Freshwater Biology*, *54*(10), 2086–2100. https://doi.org/10.1111/j.1365-2427.2008.02150.x | Forest | Clear-cut | Mixed | boreal | harvest | local+catchment | detritus, omnivore, shredder | decomposition, abundance, diversity, biomass |
| 115 | (McNeish et al., 2017) | McNeish, R. E., Benbow, M. E., & McEwan, R. W. (2017). Removal of the invasive shrub, *Lonicera maackii* (Amur Honeysuckle), from a headwater stream riparian zone shifts taxonomic and functional composition of the aquatic biota. *Invasive Plant Science and Management*, 10(3), 232–246. https://doi.org/10.1017/inp.2017.22 | Invasive species removed sites | Invaded sites | | temperate | restoration | local | omnivore, shredder | abundance, diversity |
| 116 | (McNeish et al., 2015) | McNeish, R. E., Moore, E. M., Benbow, M. E., & McEwan, R. W. (2015). Removal of the Invasive Shrub, *Lonicera maackii*, from Riparian Forests Influences Headwater Stream Biota and Ecosystem Function. *River Research and Applications*, 31(9), 1131–1139. https://doi.org/10.1002/rra.2808 | Invasive species removed sites | Invaded sites | | temperate | restoration | local | detritus | biomass |
| 117 | (Mctammany et al., 2008) | Mctammany, M. E., Benfield, E. F., & Webster, J. R. (2008). Effects of agriculture on wood breakdown and microbial biofilm respiration in southern Appalachian streams. *Freshwater Biology*, 53(4), 842–854. https://doi.org/10.1111/j.1365-2427.2007.01936.x | Forest; Recovered | AG-L, AG-H | Other sites | temperate | conversion, restoration | local, local+catchment, catchment | detritus | decomposition |
| 118 | (Menéndez et al., 2013) | Menéndez, M., Descals, E., Riera, T., & Moya, O. (2013). Do non-native *Platanus hybrida* riparian plantations affect leaf litter decomposition in streams? *Hydrobiologia*, *716*(1), 5–20. https://doi.org/10.1007/s10750-013-1539-0 | Forest | Plantation | | temperate | plantation | local | detritus, omnivore, shredder, microbes | decomposition, abundance, diversity |
| 119 | (Menninger & Palmer, 2007) | Menninger, H. L., & Palmer, M. A. (2007). Herbs and grasses as an allochthonous resource in open-canopy headwater streams. *Freshwater Biology*, 52(9), 1689–1699. https://doi.org/10.1111/j.1365-2427.2007.01797.x | RB | FCQ | CC | temperate | conversion | catchment | detritus, omnivore, shredder | biomass, abundance |
| 120 | (Mesa et al., 2013) | Mesa, L. M., Reynaga, M. C., Correa, M. del V., & Sirombra, M. G. (2013). Effects of anthropogenic impacts on benthic macroinvertebrates assemblages in subtropical mountain streams. *Iheringia. Série Zoologia*, 103, 342–349. https://doi.org/10.1590/S0073-47212013000400002 | Unimpacted | Impacted | | subtropical | conversion | local | detritus, shredder | biomass, abundance, diversity |
| 121 | (Mlambo et al., 2019) | Mlambo, M. C., Paavola, R., Fritze, H., Louhi, P., & Muotka, T. (2019). Leaf litter decomposition and decomposer communities in streams affected by intensive forest biomass removal. *Ecological Indicators*, *101*, 364–372. https://doi.org/10.1016/j.ecolind.2019.01.035 | Natural | Logged with stumps removed | Logged without stumps removed | boreal | harvest | catchment | detritus, shredder, microbes | decomposition, abundance, diversity, biomass |
| 122 | (Molinero et al., 1996) | Molinero, J., Pozo, J., & Gonzalez, E. (1996). Litter breakdown in streams of the Agüera catchment: Influence of dissolved nutrients and land use. Freshwater Biology, 36(3), 745–756. https://doi.org/10.1046/j.1365-2427.1996.00125.x | Forest | Plantation | | temperate | plantation | local+catchment | detritus | decomposition, biomass |

| | | | | | | | | | |
|---|---|---|---|---|---|---|---|---|---|
| 123 | (Monroy et al., 2017) | Monroy, S., Martínez, A., López-Rojo, N., Pérez-Calpe, A. V., Basaguren, A., & Pozo, J. (2017). Structural and functional recovery of macroinvertebrate communities and leaf litter decomposition after a marked drought: Does vegetation type matter? *Science of the Total Environment*, *599–600*, 1241–1250. https://doi.org/10.1016/j.scitotenv.2017.05.093 | Deciduous forest | Eucalyptus and Pine plantation | | temperate | plantation | local+catchment | omnivore, shredder, detritus | abundance, diversity, biomass, decomposition |
| 124 | (Morabowen et al., 2019) | Morabowen, A., Crespo-Pérez, V., & Ríos-Touma, B. (2019). Effects of agricultural landscapes and land uses in highly biodiverse tropical streams of the Ecuadorian Choco. *Inland Waters*, *9*(3), 289–300. https://doi.org/10.1080/20442041.2018.1527597 | Forest | Organic farming; Monoculture | | tropical | conversion | catchment, local+catchment | omnivore, detritus | diversity, biomass |
| 125 | (Murphy & Giller, 2001) | Murphy, J. F., & Giller, P. S. (2001). Hazel leaf breakdown in two low-order streams differing in the functional efficiency of their detritivore assemblages. *Archiv für Hydrobiologie*, 249–267. https://doi.org/10.1127/archiv-hydrobiol/150/2001/249 | Forest | Conifer plantation | Transplantation experiment results | temperate | plantation | local+catchment | detritus, omnivore, shredder | decomposition, abundance |
| 126 | (Musetta-Lambert et al., 2017) | Musetta-Lambert, J., Muto, E., Kreutzweiser, D., & Sibley, P. (2017). Wildfire in boreal forest catchments influences leaf litter subsidies and consumer communities in streams: Implications for riparian management strategies. *Forest Ecology and Management*, *391*, 29–41. https://doi.org/10.1016/j.foreco.2017.01.028 | Forest | Harvested | Fire | boreal | harvest | local+catchment | detritus, omnivore, shredder | biomass, decomposition, abundance, diversity |
| 127 | (Myers et al., 2007) | Myers, L., Mihuc, T., & Woodcock, T. (2007). The impacts of forest management on invertebrate communities associated with submerged leaves in forested Adirondack streams. *Journal of Freshwater Ecology*, *22*(2), 325–331. https://doi.org/10.1080/02705060.2007.9665054 | Preserve | Logged | | temperate | harvest | catchment | omnivore | abundance |
| 128 | (Oelbermann & Gordon, 2000) | Oelbermann, M., & Gordon, A. M. (2000). Quantity and quality of autumnal litterfall into a rehabilitated agricultural stream. *Journal of Environmental Quality*, *29*(2), 603–611. https://doi.org/10.2134/jeq2000.00472425002900020031x | Wide buffer | Narrow buffer | Other sites | temperate | conversion | local | detritus | biomass |
| 129 | (Oester et al., 2023) | Oester, R., dos Reis Oliveira, P. C., Moretti, M. S., Altermatt, F., & Bruder, A. (2023). Leaf-associated macroinvertebrate assemblage and leaf litter breakdown in headwater streams depend on local riparian vegetation. *Hydrobiologia*, *850*(15), 3359–3374. https://doi.org/10.1007/s10750-022-05049-7 | Forested | Non-forested | | temperate | harvest | local | detritus, omnivore, shredder | decomposition, abundance, diversity, biomass |
| 130 | (Ono et al., 2020) | Ono, E. R., Manoel, P. S., Melo, A. L. U., & Uieda, V. S. (2020). Effects of riparian vegetation removal on the functional feeding group structure of benthic macroinvertebrate assemblages. *Community Ecology*, *21*(2), 145–157. https://doi.org/10.1007/s42974-020-00014-7 | Forest | Pasture | | tropical | conversion | local | shredder, omnivore, detritus | abundance, diversity |
| 131 | (Paul et al., 2006) | Paul, M. J., Meyer, J. L., & Couch, C. A. (2006). Leaf breakdown in streams differing in catchment land use. | Forest | Agriculture | Suburban and urban sites | temperate | conversion | catchment | detritus, microbes, shredder | decomposition, biomass, abundance |

| | | | | | | | | | | |
|---|---|---|---|---|---|---|---|---|---|---|
| | | *Freshwater Biology*, *51*(9), 1684–1695. https://doi.org/10.1111/j.1365-2427.2006.01612.x | | | | | | | | |
| 132 | (Principe et al., 2015) | Principe, R. E., Márquez, J. A., Martina, L. C., Jobbágy, E. G., & Albariño, R. J. (2015). Pine afforestation changes more strongly community structure than ecosystem functioning in grassland mountain streams. *Ecological Indicators*, *57*, 366–375. https://doi.org/10.1016/j.ecolind.2015.04.033 | Natural grassland | Pine plantation | | subtropical | plantation | local+catchment | detritus, omnivore, shredder | decomposition, diversity, abundance |
| 133 | (Reid et al., 2013) | Reid, D. J., Lake, P. S., & Quinn, G. P. (2013). Influences of agricultural landuse and seasonal changes in abiotic conditions on invertebrate colonisation of riparian leaf detritus in intermittent streams. *Aquatic Sciences*, *75*(2), 285–297. https://doi.org/10.1007/s00027-012-0273-4 | Reserve | Farmland | | subtropical | conversion | local | omnivore | diversity, abundance |
| 134 | (Rezende et al., 2021) | Rezende, R. de S., Cararo, E. R., Bernardi, J. P., Chimello, V., Lima-Rezende, C. A., Albeny-Simões, D., Dal Magro, J., & Gonçalves, J. F. (2021). Land cover affects the breakdown of *Pinus elliottii* needles litter by microorganisms in soil and stream systems of subtropical riparian zones. *Limnologica*, *90*, 125905. https://doi.org/10.1016/j.limno.2021.125905 | Forest, grassland with riparian vegetation | Silvicultre, grassland without riparian vegetation | | subtropical | harvest, conversion | local+catchment, local | detritus | biomass |
| 135 | (Riipinen et al., 2009) | Riipinen, M. P., Fleituch, T., Hladyz, S., Woodward, G., Giller, P., & Dobson, M. (2009). Invertebrate community structure and ecosystem functioning in European conifer plantation streams. *Freshwater Biology*, 55, 346–359. https://doi.org/10.1111/j.1365-2427.2009.02278.x | Broadleaf forest | Conifer plantation | | temperate | plantation | local+catchment | detritus, shredder | decomposition, abundance, diversity, biomass |
| 137 | (Roberts & Bilby, 2009) | Roberts, M. L., & Bilby, R. E. (2009). Urbanization alters litterfall rates and nutrient inputs to small Puget Lowland streams. *Journal of the North American Benthological Society*, *28*(4), 941–954. https://doi.org/10.1899/07-160.1 | Reference | Disturbed | Intermediate sites | temperate | conversion | local | detritus | biomass |
| 138 | (Rossi et al., 2019) | Rossi, F., Mallet, C., Portelli, C., Donnadieu, F., Bonnemoy, F., & Artigas, J. (2019). Stimulation or inhibition: Leaf microbial decomposition in streams subjected to complex chemical contamination. *Science of the Total Environment*, *648*, 1371–1383. https://doi.org/10.1016/j.scitotenv.2018.08.197 | Forest | Agriculure | Urban | temperate | conversion | catchment | detritus, microbes | decomposition, biomass, diversity |
| 140 | (Rubio-Ríos et al., 2023) | Rubio-Ríos, J., Salinas-Bonillo, M. J., Pérez, J., Fenoy, E., Boyero, L., & Casas, J. J. (2023). Alder stands promote N-cycling but not leaf litter mass loss in Mediterranean streams flowing through pine plantations. *Forest Ecology and Management*, *542*, 121072. https://doi.org/10.1016/j.foreco.2023.121072 | With Alder | Without Alder | Mixtures | temperate | restoration | local | detritus | decomposition |
| 141 | (Ryan & Kelly-Quinn, 2016) | Ryan, D. K., & Kelly-Quinn, M. (2016). Riparian vegetation management for water temperature regulation: Implications for the production of macroinvertebrate prey of salmonids. *Fisheries Management and Ecology*, *23*(6), 519–530. https://doi.org/10.1111/fme.12193 | Shaded riparian cover | Unshaded riparian cover | | temperate | conversion | local | omnivore | abundance |
| 142 | (Ryder et al., 2011) | Ryder, L., de Eyto, E., Gormally, M., Skeffington, M. S., Dillane, M., & Poole, R. (2011). Riparian zone creation in established coniferous forests in Irish upland peat | Pre-felling | Post-felling | Other sites | temperate | harvest | local | omnivore | diversity, abundance |

| | | | | | | | | | | |
|---|---|---|---|---|---|---|---|---|---|---|
| | | catchments: Physical, chemical and biological implications. *Biology and Environment: Proceedings of the Royal Irish Academy*, *111B*(1), 41–60. | | | | | | | | |
| 143 | (Sakai et al., 2013) | Sakai, M., Natuhara, Y., Fukushima, K., Imanishi, A., Imai, K., & Kato, M. (2013). Ecological functions of persistent Japanese cedar litter in structuring stream macroinvertebrate assemblages. *Journal of Forest Research*, *18*(2), 190–199. https://doi.org/10.1007/s10310-012-0339-0 | Forest | Plantation | | temperate | plantation | catchment | detritus, omnivore, shredder | biomass, abundance, diversity |
| 145 | (Sarriquet et al., 2006) | Sarriquet, P. E., Delettre, Y. R., & Marmonier, P. (2006). Effects of catchment disturbance on stream invertebrates: Comparison of different habitats (vegetation, benthic and interstitial) using bio-ecological groups. *Annales de Limnologie*, *42*(4), 205–219. https://doi.org/10.1051/limn/2006022 | Forest | Agriculture | Intermediate and otherwise heavily disturbed sites | temperate | conversion | local | omnivore | diversity, abundance |
| 146 | (Shah et al., 2021) | Shah, N. W., Nisbet, T. R., & Broadmeadow, S. B. (2021). The impacts of conifer afforestation and climate on water quality and freshwater ecology in a sensitive peaty catchment: A 25 year study in the upper River Halladale in North Scotland. *Forest Ecology and Management*, *502*, 119616. https://doi.org/10.1016/j.foreco.2021.119616 | BB | UH | Other sites | temperate | conversion | local+catchment | omnivore | diversity, abundance |
| 147 | (Silva-Araújo et al., 2020) | Silva-Araújo, M., Silva-Junior, E. F., Neres-Lima, V., Feijó-Lima, R., Tromboni, F., Lourenço-Amorim, C., Thomas, S. A., Moulton, T. P., & Zandonà, E. (2020). Effects of riparian deforestation on benthic invertebrate community and leaf processing in Atlantic forest streams. *Perspectives in Ecology and Conservation*, *18*(4), 277–282. https://doi.org/10.1016/j.pecon.2020.09.004 | Least deforested | Most deforested | Intermediate sites | tropical | harvest | local | detritus, omnivore, shredder | decomposition, abundance, diversity |
| 150 | (Suga & Tanaka, 2013) | Suga, C. M., & Tanaka, M. O. (2013). Influence of a forest remnant on macroinvertebrate communities in a degraded tropical stream. *Hydrobiologia*, *703*(1), 203–213. https://doi.org/10.1007/s10750-012-1360-1 | Downstream forest | Upstream sugarcane | | tropical | conversion | local | omnivore, shredder | abundance, diversity |
| 151 | (Tanaka et al., 2015) | Tanaka, M. O., Fernandes, J. de F., Suga, C. M., Hanai, F. Y., & Souza, A. L. T. de. (2015). Abrupt change of a stream ecosystem function along a sugarcane-forest transition: Integrating riparian and in-stream characteristics. *Agriculture, Ecosystems & Environment*, 207, 171–177. https://doi.org/10.1016/j.agee.2015.04.014 | Downstream forest | Upstream sugarcane | | tropical | conversion | local | detritus | decomposition |
| 152 | (Tolkkinen et al., 2015) | Tolkkinen, M., Mykrä, H., Annala, M., Markkola, A. M., Vuori, K. M., & Muotka, T. (2015). Multi-stressor impacts on fungal diversity and ecosystem functions in streams: Natural vs. anthropogenic stress. *Ecology*, *96*(3), 672–683. https://doi.org/10.1890/14-0743.1 | Pristine | disturbed | | boreal | harvest | local+catchment | detritus, microbes | decomposition, biomass, diversity |
| 153 | (Tonello et al., 2021) | Tonello, G., Decian, V. S., Restello, R. M., & Hepp, L. U. (2021). The conversion of natural riparian forests into agricultural land affects ecological processes in Atlantic forest streams. *Limnologica*, *91*, 125927. https://doi.org/10.1016/j.limno.2021.125927 | Least agriculture | Most agriculture | Intermediate sites | subtropical | conversion | local | detritus, omnivore, shredder | decomposition, abundance |

| | | | | | | | | | |
|---|---|---|---|---|---|---|---|---|---|
| 154 | (Torres & Ramírez, 2014) | Torres, P. J., & Ramírez, A. (2014). Land use effects on leaf litter breakdown in low-order streams draining a rapidly developing tropical watershed in Puerto Rico. *Revista de Biología Tropical*, *62*(S2), Article S2. https://doi.org/10.15517/rbt.v62i0.15783 | Forest | Agriculture | Urban | tropical | conversion | local+catchment | detritus | decomposition |
| 155 | (Townsend et al., 2004) | Townsend, C. R., Downes, B. J., Peacock, K., & Arbuckle, C. J. (2004). Scale and the detection of land-use effects on morphology, vegetation and macroinvertebrate communities of grassland streams. *Freshwater Biology*, *49*(4), 448–462. https://doi.org/10.1111/j.1365-2427.2004.01192.x | Tussock | Pasture | | temperate | conversion | local+catchment | omnivore | diversity, abundance |
| 156 | (Truchy et al., 2022) | Truchy, A., Sponseller, R. A., Ecke, F., Angeler, D. G., Kahlert, M., Bundschuh, M., Johnson, R. K., & McKie, B. G. (2022). Responses of multiple structural and functional indicators along three contrasting disturbance gradients. *Ecological Indicators*, *135*, 108514. https://doi.org/10.1016/j.ecolind.2021.108514 | Most forested | Least forested | Intermediate sites | boreal | harvest | catchment | omnivore, shredder, detritus, microbes | abundance, diversity, decomposition, biomass |
| 157 | (Tupinambás et al., 2007) | Tupinambás, T. H., Callisto, M., & Santos, G. B. (2007). Benthic macroinvertebrate assemblages structure in two headwater streams, south-eastern Brazil. *Revista Brasileira de Zoologia*, 24(4), 887–897. https://doi.org/10.1590/S0101-81752007000400005 | 1B, 2B | 1A, 2A ,3A, 3B | | tropical | conversion | local | omnivore | abundance, diversity |
| 158 | (Turunen et al., 2017) | Turunen, J., Aroviita, J., Marttila, H., Louhi, P., Laamanen, T., Tolkkinen, M., Luhta, P.-L., Kløve, B., & Muotka, T. (2017). Differential responses by stream and riparian biodiversity to in-stream restoration of forestry-impacted streams. *Journal of Applied Ecology*, *54*(5), 1505–1514. https://doi.org/10.1111/1365-2664.12897 | Pristine | Impaired | Restored sites | boreal | harvest | local+catchment | omnivore, detritus, microbes | abundance, diversity, decomposition, biomass |
| 159 | (Valente-Neto et al., 2015) | Valente-Neto, F., Koroiva, R., Fonseca-Gessner, A. A., & Roque, F. de O. (2015). The effect of riparian deforestation on macroinvertebrates associated with submerged woody debris. *Aquatic Ecology*, 49(1), 115–125. https://doi.org/10.1007/s10452-015-9510-y | Most complete riparian zone | Least complete riparian zone | Intermediate sites | tropical | conversion | local | omnivore | abundance, diversity |
| 161 | (Walsh et al., 2002) | Walsh, C. J., Gooderham, J. P. R., Grace, M. R., Sdraulig, S., Rosyidi, M. I., & Lelono, A. (2002). The relative influence of diffuse- and point-source disturbances on a small upland stream in East Java, Indonesia: A preliminary investigation. *Hydrobiologia*, *487*(1), 183–192. https://doi.org/10.1023/A:1022990822198 | UF | UV | Intermediate sites | tropical | plantation | local | omnivore | diversity |
| 162 | (Whiles & Wallace, 1997) | Whiles, M. R., & Wallace, J. B. (1997). Leaf litter decomposition and macroinvertebrate communities in headwater streams draining pine and hardwood catchments. *Hydrobiologia*, *353*(1), 107–119. https://doi.org/10.1023/A:1003054827248 | Hardwood | Pine plantation | | temperate | plantation | catchment | shredder, omnivore, detritus | abundance, biomass, decomposition |
| 163 | (Yeung et al., 2017) | Yeung, A. C. Y., Lecerf, A., & Richardson, J. S. (2017). Assessing the long-term ecological effects of riparian management practices on headwater streams in a coastal temperate rainforest. *Forest Ecology and Management*, | Forest | Clear-cut | | temperate | harvest | local+catchment | detritus, shredder | decomposition, abundance, diversity |

| | | | | | | | | | |
|---|---|---|---|---|---|---|---|---|---|
| | | *384*, 100–109. https://doi.org/10.1016/j.foreco.2016.10.044 | | | | | | | |
| 164 | (Youngquist et al., 2020) | Youngquist, M. B., Wiley, C., Eggert, S. L., D’Amato, A. W., Palik, B. J., & Slesak, R. A. (2020). Foundation species loss affects leaf breakdown and aquatic invertebrate resource use in black ash wetlands. *Wetlands*, *40*(4), 839–852. https://doi.org/10.1007/s13157-019-01221-3 | Forest | Clear-cut | temperate | harvest | local | detritus | decomposition |
| 166 | (Zhang et al., 2009) | Zhang, Y., Richardson, J. S., & Pinto, X. (2009). Catchment-scale effects of forestry practices on benthic invertebrate communities in Pacific coastal streams. *Journal of Applied Ecology*, *46*(6), 1292–1303. https://doi.org/10.1111/j.1365-2664.2009.01718.x | Forest | Logged | temperate | harvest | catchment | omnivore, shredder | biomass, abundance |

### Full reference list


Abelho, M., & Graça, M. A. S. (1996). Effects of eucalyptus afforestation on leaf litter dynamics and macroinvertebrate community structure of streams in Central Portugal. *Hydrobiologia*, *324*(3), 195–204. https://doi.org/10.1007/BF00016391

Aguiar, A. C. F., Neres-Lima, V., & Moulton, T. P. (2018). Relationships of shredders, leaf processing and organic matter along a canopy cover gradient in tropical streams. *Journal of Limnology*, *77*(1). https://doi.org/10.4081/jlimnol.2017.1684

Albertson, L. K., Ouellet, V., & Daniels, M. D. (2018). Impacts of stream riparian buffer land use on water temperature and food availability for fish. *Journal of Freshwater Ecology*, *33*(1), 195–210. https://doi.org/10.1080/02705060.2017.1422558

Astudillo, M. R., Novelo-Gutiérrez, R., Vázquez, G., García-Franco, J. G., & Ramírez, A. (2016). Relationships between land cover, riparian vegetation, stream characteristics, and aquatic insects in cloud forest streams, Mexico. *Hydrobiologia*, *768*(1), 167–181. https://doi.org/10.1007/s10750-015-2545-1

Bacca, J. C., Rampanelli Cararo, E., Alves Lima-Rezende, C., Tavares Martins, R., Macedo-Reis, L. E., Dal Magro, J., & De Souza Rezende, R. (2023). Land-use effects on aquatic macroinvertebrate diversity in subtropical highland grasslands streams. *Limnetica*, *42*(2), 1. https://doi.org/10.23818/limn.42.16

Bärlocher, F., Helson, J. E., & Williams, D. D. (2010). Aquatic hyphomycete communities across a land-use gradient of Panamanian streams. *Fundamental and Applied Limnology*, *177*(3), 209–221. https://doi.org/10.1127/1863-9135/2010/0177-0209

Barrios, M., Burwood, M., Kröger, A., Calvo, C., Ríos-Touma, B., & Teixeira-de-Mello, F. (2022). Riparian cover buffers the effects of abiotic and biotic predictors of leaf decomposition in subtropical streams. *Aquatic Sciences*, *84*(4), 55. https://doi.org/10.1007/s00027-022-00886-z

Basaguren, A., & Pozo, J. (1994). Leaf litter processing of alder and eucalyptus in the Agüera stream system (Northern Spain). II: Macroinvertebrates associated. *Archiv für Hydrobiologie*, *132*(1), 57–68.

Benfield, E. F., Webster, J. R., Tank, J. L., & Hutchens, J. J. (2001). Long-term patterns in leaf breakdown in streams in response to watershed logging. *International Review of Hydrobiology*, *86*(4–5), 467–474. https://doi.org/10.1002/1522-2632(200107)86:4/5<467::AID-IROH467>3.0.CO;2-1

Bertaso, T. R. N., Spies, M. R., Kotzian, C. B., & Flores, M. L. T. (2015). Effects of forest conversion on the assemblages' structure of aquatic insects in subtropical regions. *Revista Brasileira de Entomologia*, *59*(1), 43–49. https://doi.org/10.1016/j.rbe.2015.02.005

Burrows, R. M., Magierowski, R. H., Fellman, J. B., & Barmuta, L. A. (2012). Woody debris input and function in old-growth and clear-felled headwater streams. *Forest Ecology and Management*, *286*, 73–80. https://doi.org/10.1016/j.foreco.2012.08.038

Burrows, R. M., Magierowski, R. H., Fellman, J. B., Clapcott, J. E., Munks, S. A., Roberts, S., Davies, P. E., & Barmuta, L. A. (2014). Variation in stream organic matter processing among years and benthic habitats in response to forest clearfelling. *Forest Ecology and Management*, *327*, 136–147. https://doi.org/10.1016/j.foreco.2014.04.041

Burwood, M., Clemente, J., Meerhoff, M., Iglesias, C., Goyenola, G., Fosalba, C., Pacheco, J. P., & Teixeira De Mello, F. (2021). Macroinvertebrate communities and macrophyte decomposition could be affected by land use intensification in subtropical lowland streams. *Limnetica*, *40*(2), 343–357. https://doi.org/10.23818/limn.40.23

Bylak, A., Kukuła, K., Ortyl, B., Hałoń, E., Demczyk, A., Janora-Hołyszko, K., Maternia, J., Szczurowski, Ł., & Ziobro, J. (2022). Small stream catchments in a developing city context: The importance of land cover changes on the ecological status of streams and the possibilities for providing ecosystem services. *Science of the Total Environment*, *815*, 151974. https://doi.org/10.1016/j.scitotenv.2021.151974

Carroll, D., & Jackson, G. R. (2009). Observed relationships between urbanization and riparian cover, shredder abundance, and stream leaf litter standing crops. *Fundamental and Applied Limnology*, 213–225. https://doi.org/10.1127/1863-9135/2008/0173-0213

Casas, J. J., Larrañaga, A., Menéndez, M., Pozo, J., Basaguren, A., Martínez, A., Pérez, J., González, J. M., Mollá, S., Casado, C., Descals, E., Roblas, N., López-González, J. A., & Valenzuela, J. L. (2013). Leaf litter decomposition of native and introduced tree species of contrasting quality in headwater streams: How does the regional setting matter? *Science of the Total Environment*, *458–460*, 197–208. https://doi.org/10.1016/j.scitotenv.2013.04.004

Casotti, C. G., Kiffer, W. P., Costa, L. C., Rangel, J. V., Casagrande, L. C., & Moretti, M. S. (2015). Assessing the importance of riparian zones conservation for leaf decomposition in streams. *Natureza & Conservação*, *13*(2), 178–182. https://doi.org/10.1016/j.ncon.2015.11.011

Chellaiah, D., & Yule, C. M. (2018). Litter decomposition is driven by microbes and is more influenced by litter quality than environmental conditions in oil palm streams with different riparian types. *Aquatic Sciences*, *80*(4), 43. https://doi.org/10.1007/s00027-018-0595-y
Chellaiah, D., & Yule, C. M. (2019). Litter quality influences bacterial communities more strongly than changes in riparian buffer quality in oil palm streams. *Aquatic Microbial Ecology*, *83*(2), 167–179. https://doi.org/10.3354/ame01909
Collier, K. J., & Halliday, J. N. (2000). Macroinvertebrate-wood associations during decay of plantation pine in New Zealand pumice-bed streams: Stable habitat or trophic subsidy? *Journal of the North American Benthological Society*, *19*(1), 94–111. https://doi.org/10.2307/1468284
Collier, K. J., Smith, B. J., & Halliday, N. J. (2004). Colonization and use of pine wood versus native wood in New Zealand plantation forest streams: Implications for riparian management. *Aquatic Conservation: Marine and Freshwater Ecosystems*, *14*(2), 179–199. https://doi.org/10.1002/aqc.599
Cordero–Rivera, A., Álvarez, A. M., & Álvarez, M. (2017). Eucalypt plantations reduce the diversity of macroinvertebrates in small forested streams. *Animal Biodiversity and Conservation*, *40*(1), Article 1. https://doi.org/10.32800/abc.2017.40.0087
Cornejo, A., Pérez, J., López-Rojo, N., Tonin, A. M., Rovira, D., Checa, B., Jaramillo, N., Correa, K., Villarreal, A., Villarreal, V., García, G., Pérez, E., Ríos González, T. A., Aguirre, Y., Correa-Araneda, F., & Boyero, L. (2020). Agriculture impairs stream ecosystem functioning in a tropical catchment. *Science of the Total Environment*, *745*, 140950. https://doi.org/10.1016/j.scitotenv.2020.140950
da Costa, I. D., Petry, A. C., & Mazzoni, R. (2020). Fish assemblages respond to forest cover in small Amazonian basins. *Limnologica*, *81*, 125757. https://doi.org/10.1016/j.limno.2020.125757
Danger, A. R., & Robson, B. J. (2004). The effects of land use on leaf-litter processing by macroinvertebrates in an Australian temperate coastal stream. *Aquatic Sciences*, *66*(3), 296–304. https://doi.org/10.1007/s00027-004-0718-5
Daoust, K., Kreutzweiser, D. P., Guo, J., Creed, I. F., & Sibley, P. K. (2019). Climate-influenced catchment hydrology overrides forest management effects on stream benthic macroinvertebrates in a northern hardwood forest. *Forest Ecology and Management*, *452*, 117540. https://doi.org/10.1016/j.foreco.2019.117540
de Mello Cionek, V., Fogaça, F. N. O., Moulton, T. P., Pazianoto, L. H. R., Landgraf, G. O., & Benedito, E. (2021). Influence of leaf miners and environmental quality on litter breakdown in tropical headwater streams. *Hydrobiologia*, *848*(6), 1311–1331. https://doi.org/10.1007/s10750-021-04529-6
de Nadaï-Monoury, E., Gilbert, F., & Lecerf, A. (2014). Forest canopy cover determines invertebrate diversity and ecosystem process rates in depositional zones of headwater streams. *Freshwater Biology*, *59*(7), 1532–1545. https://doi.org/10.1111/fwb.12364
Delong, M. D., & Brusven, M. A. (1994). Allochthonous input of organic matter from different riparian habitats of an agriculturally impacted stream. *Environmental Management*, *18*(1), 59–71. https://doi.org/10.1007/BF02393750
Dézerald, O., Talaga, S., Leroy, C., Carrias, J.-F., Corbara, B., Dejean, A., & Céréghino, R. (2014). Environmental determinants of macroinvertebrate diversity in small water bodies: Insights from tank-bromeliads. *Hydrobiologia*, *723*(1), 77–86. https://doi.org/10.1007/s10750-013-1464-2
Díez, J., Elosegi, A., Chauvet, E., & Pozo, J. (2002). Breakdown of wood in the Agüera stream. *Freshwater Biology*, *47*(11), 2205–2215. https://doi.org/10.1046/j.1365-2427.2002.00965.x
dos Santos, F. B., Ferreira, F. C., & Esteves, K. E. (2015). Assessing the importance of the riparian zone for stream fish communities in a sugarcane dominated landscape (Piracicaba River Basin, Southeast Brazil). *Environmental Biology of Fishes*, *98*(8), 1895–1912. https://doi.org/10.1007/s10641-015-0406-4
Dudgeon, D. (2006). The impacts of human disturbance on stream benthic invertebrates and their drift in North Sulawesi, Indonesia. *Freshwater Biology*, *51*(9), 1710–1729. https://doi.org/10.1111/j.1365-2427.2006.01596.x
Dudley, M. P., Solomon, K., Wenger, S., Jackson, C. R., Freeman, M., Elliott, K. J., Miniat, C. F., & Pringle, C. M. (2021). Do crayfish affect stream ecosystem response to riparian vegetation removal? *Freshwater Biology*, *66*(7), 1423–1435. https://doi.org/10.1111/fwb.13728
Ebling, L. A., Pastore, B. L., Biasi, C., Hepp, L. U., & Restello, R. M. (2024). Does environmental variability in Atlantic Forest streams affect aquatic hyphomycete and invertebrate assemblages associated with leaf litter? *Hydrobiologia*, *851*(7), 1761–1777. https://doi.org/10.1007/s10750-023-05415-z
Elosegi, A., Díez, J., & Pozo, J. (2007). Contribution of dead wood to the carbon flux in forested streams. *Earth Surface Processes and Landforms*, *32*(8), 1219–1228. https://doi.org/10.1002/esp.1549

Elosegi, A., Nicolás, A., & Richardson, J. S. (2018). Priming of leaf litter decomposition by algae seems of minor importance in natural streams during autumn. *PLOS ONE*, *13*(9), e0200180. https://doi.org/10.1371/journal.pone.0200180

Ely, D. T., & Wallace, J. B. (2010). Long-term functional group recovery of lotic macroinvertebrates from logging disturbance. *Canadian Journal of Fisheries and Aquatic Sciences*, *67*(7), 1126–1134. https://doi.org/10.1139/F10-045

Emilson, C. E., Kreutzweiser, D. P., Gunn, J. M., & Mykytczuk, N. C. S. (2016). Effects of land use on the structure and function of leaf-litter microbial communities in boreal streams. *Freshwater Biology*, *61*(7), 1049–1061. https://doi.org/10.1111/fwb.12765

Encalada, A. C., Calles, J., Ferreira, V., Canhoto, C. M., & Graça, M. A. S. (2010). Riparian land use and the relationship between the benthos and litter decomposition in tropical montane streams. *Freshwater Biology*, *55*(8), 1719–1733. https://doi.org/10.1111/j.1365-2427.2010.02406.x

Erdozain, M., Kidd, K. A., Emilson, E. J. S., Capell, S. S., Luu, T., Kreutzweiser, D. P., & Gray, M. A. (2021). Forest management impacts on stream integrity at varying intensities and spatial scales: Do biological effects accumulate spatially? *Science of the Total Environment*, *763*, 144043. https://doi.org/10.1016/j.scitotenv.2020.144043

Erdozain, M., Kidd, K., Kreutzweiser, D., & Sibley, P. (2018). Linking stream ecosystem integrity to catchment and reach conditions in an intensively managed forest landscape. *Ecosphere*, *9*(5), e02278. https://doi.org/10.1002/ecs2.2278

Espinoza-Toledo, A., Mendoza-Carranza, M., Castillo, M. M., Barba-Macías, E., & Capps, K. A. (2021). Taxonomic and functional responses of macroinvertebrates to riparian forest conversion in tropical streams. *Science of the Total Environment*, *757*, 143972. https://doi.org/10.1016/j.scitotenv.2020.143972

Evangelista, C., Boiche, A., Lecerf, A., & Cucherousset, J. (2014). Ecological opportunities and intraspecific competition alter trophic niche specialization in an opportunistic stream predator. *Journal of Animal Ecology*, *83*(5), 1025–1034. https://doi.org/10.1111/1365-2656.12208

Fargen, C., Emery, S. M., & Carreiro, M. M. (2015). Influence of *Lonicera maackii* invasion on leaf litter decomposition and macroinvertebrate communities in an urban stream. *Natural Areas Journal*, *35*(3), 392–403. https://doi.org/10.3375/043.035.0303

Fasching, C., Akotoye, C., Bižić, M., Fonvielle, J., Ionescu, D., Mathavarajah, S., Zoccarato, L., Walsh, D. A., Grossart, H.-P., & Xenopoulos, M. A. (2020). Linking stream microbial community functional genes to dissolved organic matter and inorganic nutrients. *Limnology and Oceanography*, *65*(S1), S71–S87. https://doi.org/10.1002/lno.11356

Ferreira, V., Boyero, L., Calvo, C., Correa, F., Figueroa, R., Gonçalves, J. F., Goyenola, G., Graça, M. A. S., Hepp, L. U., Kariuki, S., López-Rodríguez, A., Mazzeo, N., M'Erimba, C., Monroy, S., Peil, A., Pozo, J., Rezende, R., & Teixeira-de-Mello, F. (2019). A Global Assessment of the Effects of Eucalyptus Plantations on Stream Ecosystem Functioning. *Ecosystems*, *22*(3), 629–642. https://doi.org/10.1007/s10021-018-0292-7

Ferreira, V., Elosegi, A., Gulis, V., Pozo, J., & Graça, M. A. S. (2006). Eucalyptus plantations affect fungal communities associated with leaf-litter decomposition in Iberian streams. *Archiv für Hydrobiologie*, *166*, 467–490. https://doi.org/10.1127/0003-9136/2006/0166-0467

Ferreira, V., Faustino, H., Raposeiro, P. M., & Gonçalves, V. (2017). Replacement of native forests by conifer plantations affects fungal decomposer community structure but not litter decomposition in Atlantic island streams. *Forest Ecology and Management*, *389*, 323–330. https://doi.org/10.1016/j.foreco.2017.01.004

Ferreira, V., Larrañaga, A., Gulis, V., Basaguren, A., Elosegi, A., Graça, M. A. S., & Pozo, J. (2015). The effects of eucalypt plantations on plant litter decomposition and macroinvertebrate communities in Iberian streams. *Forest Ecology and Management*, *335*, 129–138. https://doi.org/10.1016/j.foreco.2014.09.013

Ferreira, W. R., Ligeiro, R., Macedo, D. R., Hughes, R. M., Kaufmann, P. R., Oliveira, L. G., & Callisto, M. (2015). Is the diet of a typical shredder related to the physical habitat of headwater streams in the Brazilian Cerrado? *Annales de Limnologie*, *51*(2), 115–127. https://doi.org/10.1051/limn/2015004

Four, B., Arce, E., Danger, M., Gaillard, J., Thomas, M., & Banas, D. (2017). Catchment land use-dependent effects of barrage fishponds on the functioning of headwater streams. *Environmental Science and Pollution Research*, *24*(6), 5452–5468. https://doi.org/10.1007/s11356-016-8273-x

Fu, L., Jiang, Y., Ding, J., Liu, Q., Peng, Q.-Z., & Kang, M.-Y. (2016). Impacts of land use and environmental factors on macroinvertebrate functional feeding groups in the Dongjiang River basin, southeast China. *Journal of Freshwater Ecology*, *31*(1), 21–35. https://doi.org/10.1080/02705060.2015.1017847

Fugère, V., Jacobsen, D., Finestone, E. H., & Chapman, L. J. (2018). Ecosystem structure and function of afrotropical streams with contrasting land use. *Freshwater Biology*, *63*(12), 1498–1513. https://doi.org/10.1111/fwb.13178

Giling, D. P., Nally, R. M., & Thompson, R. M. (2015). How sensitive are invertebrates to riparian-zone replanting in stream ecosystems? *Marine and Freshwater Research*, *67*(10), 1500–1511. https://doi.org/10.1071/MF14360

Godoy, B. S., Simião-Ferreira, J., Lodi, S., & Oliveira, L. G. (2016). Functional process zones characterizing aquatic insect communities in streams of the Brazilian Cerrado. *Neotropical Entomology*, *45*(2), 159–169. https://doi.org/10.1007/s13744-015-0352-z

Goodman, K. J., Hershey, A. E., & Fortino, K. (2006). The effect of forest type on benthic macroinvertebrate structure and ecological function in a pine plantation in the North Carolina Piedmont. *Hydrobiologia*, *559*(1), 305–318. https://doi.org/10.1007/s10750-005-0990-y

Goss, C. W., Goebel, P. C., & Sullivan, S. M. P. (2014). Shifts in attributes along agriculture-forest transitions of two streams in central Ohio, USA. *Agriculture, Ecosystems & Environment*, *197*, 106–117. https://doi.org/10.1016/j.agee.2014.07.026

Göthe, E., Lepori, F., & Malmqvist, B. (2009). Forestry affects food webs in northern Swedish coastal streams. *Fundamental and Applied Limnology*, *175*(4), 281–294. https://doi.org/10.1127/1863-9135/2009/0175-0281

Guevara, G., Godoy, R., Boeckx, P., Jara, C., & Oyarzún, C. (2015). Effects of riparian forest management on Chilean mountain in-stream characteristics. *Ecohydrology & Hydrobiology*, *15*(3), 160–170. https://doi.org/10.1016/j.ecohyd.2015.07.003

Guevara, G., Godoy, R., & Franco, M. (2018). Linking riparian forest harvest to benthic macroinvertebrate communities in Andean headwater streams in southern Chile. *Limnologica*, *68*, 105–114. https://doi.org/10.1016/j.limno.2017.07.007

Hagen, E. M., Webster, J. R., & Benfield, E. F. (2006). Are leaf breakdown rates a useful measure of stream integrity along an agricultural landuse gradient? *Journal of the North American Benthological Society*, *25*(2), 330–343. https://doi.org/10.1899/0887-3593(2006)25[330:ALBRAU]2.0.CO;2

Hebert, T. A., Kuehn, K. A., & Halvorson, H. M. (2023). Land use differentially alters microbial interactions and detritivore feeding during leaf decomposition in headwater streams. *Freshwater Biology*, *68*(8), 1386–1399. https://doi.org/10.1111/fwb.14111

Hisabae, M., Sone, S., & Inoue, M. (2011). Breakdown and macroinvertebrate colonization of needle and leaf litter in conifer plantation streams in Shikoku, southwestern Japan. *Journal of Forest Research*, *16*(2), 108–115. https://doi.org/10.1007/s10310-010-0210-0

Hladyz, S., Tiegs, S. D., Gessner, M. O., Giller, P. S., Rîşnoveanu, G., Preda, E., Nistorescu, M., Schindler, M., & Woodward, G. (2010). Leaf-litter breakdown in pasture and deciduous woodland streams: A comparison among three European regions. *Freshwater Biology*, *55*(9), 1916–1929. https://doi.org/10.1111/j.1365-2427.2010.02426.x

Houghton, D. C., Berry, E. A., Gilchrist, A., Thompson, J., & Nussbaum, M. A. (2011). Biological changes along the continuum of an agricultural stream: Influence of a small terrestrial preserve and use of adult caddisflies in biomonitoring. *Journal of Freshwater Ecology*, *26*(3), 381–397. https://doi.org/10.1080/02705060.2011.563513

Huryn, A. D., Huryn, V. M. B., Arbuckle, C. J., & Tsomides, L. (2002). Catchment land-use, macroinvertebrates and detritus processing in headwater streams: Taxonomic richness versus function. *Freshwater Biology*, *47*(3), 401–415. https://doi.org/10.1046/j.1365-2427.2002.00812.x

Illyová, M., Beracko, P., & Krno, I. (2011). Influence of land use on hyporheos in catchment streams of the Velka Fatra Mts. *Biologia*, *66*(2), 320–327. https://doi.org/10.2478/s11756-011-0018-1

Iñiguez-Armijos, C., Hampel, H., & Breuer, L. (2018). Land-use effects on structural and functional composition of benthic and leaf-associated macroinvertebrates in four Andean streams. *Aquatic Ecology*, *52*(1), 77–92. https://doi.org/10.1007/s10452-017-9646-z

Iñiguez-Armijos, C., Rausche, S., Cueva, A., Sánchez-Rodríguez, A., Espinosa, C., & Breuer, L. (2016). Shifts in leaf litter breakdown along a forest–pasture–urban gradient in Andean streams. *Ecology and Evolution*, *6*(14), 4849–4865. https://doi.org/10.1002/ece3.2257

Inoue, M., Shinotou, S., Maruo, Y., & Miyake, Y. (2012). Input, retention, and invertebrate colonization of allochthonous litter in streams bordered by deciduous broadleaved forest, a conifer plantation, and a clear-cut site in southwestern Japan. *Limnology*, *13*(2), 207–219. https://doi.org/10.1007/s10201-011-0369-x

Ishikawa, N. F., Togashi, H., Kato, Y., Yoshimura, M., Kohmatsu, Y., Yoshimizu, C., Ogawa, N. O., Ohte, N., Tokuchi, N., Ohkouchi, N., & Tayasu, I. (2016). Terrestrial-aquatic linkage in stream food webs along a forest

chronosequence: Multi-isotopic evidence. *Ecology*, *97*(5), 1146–1158. https://doi.org/10.1890/15-1133.1
Jinggut, T., Yule, C. M., & Boyero, L. (2012). Stream ecosystem integrity is impaired by logging and shifting agriculture in a global megadiversity center (Sarawak, Borneo). *Science of the Total Environment*, *437*, 83–90. https://doi.org/10.1016/j.scitotenv.2012.07.062
Jyväsjärvi, J., Koivunen, I., & Muotka, T. (2020). Does the buffer width matter: Testing the effectiveness of forest certificates in the protection of headwater stream ecosystems. *Forest Ecology and Management*, *478*, 118532. https://doi.org/10.1016/j.foreco.2020.118532
Kadeka, E. C., Masese, F. O., Lusega, D. M., Sitati, A., Kondowe, B. N., & Chirwa, E. R. (2021). No difference in instream decomposition among upland agricultural and forested streams in Kenya. *Frontiers in Environmental Science*, *9*. https://doi.org/10.3389/fenvs.2021.794525
Kaylor, M. J., & Warren, D. R. (2018). Canopy closure after four decades of postlogging riparian forest regeneration reduces cutthroat trout biomass in headwater streams through bottom-up pathways. *Canadian Journal of Fisheries and Aquatic Sciences*, *75*(4), 513–524. https://doi.org/10.1139/cjfas-2016-0519
Kiffer Jr, W., Giuberti, T., Victor Serpa, K., Mendes, F., & Moretti, M. (2018). Do changes in riparian zones affect periphyton growth and invertebrate colonization on rocky substrates in Atlantic Forest streams? *Iheringia Série Zoologia*, *108*, 1–10. https://doi.org/10.1590/1678-4766e2018014
Kiffney, P. M., & Richardson, J. S. (2010). Organic matter inputs into headwater streams of southwestern British Columbia as a function of riparian reserves and time since harvesting. *Forest Ecology and Management*, *260*(11), 1931–1942. https://doi.org/10.1016/j.foreco.2010.08.016
Kominoski, J. S., Marczak, L. B., & Richardson, J. S. (2011). Riparian forest composition affects stream litter decomposition despite similar microbial and invertebrate communities. *Ecology*, *92*(1), 151–159. https://doi.org/10.1890/10-0028.1
Kreutzweiser, D. P., Good, K. P., Capell, S. S., & Holmes, S. B. (2008). Leaf-litter decomposition and macroinvertebrate communities in boreal forest streams linked to upland logging disturbance. *Journal of the North American Benthological Society*, *27*(1), 1–15. https://doi.org/10.1899/07-034R.1
Laćan, I., Resh, V. H., & McBRIDE, J. R. (2010). Similar breakdown rates and benthic macroinvertebrate assemblages on native and *Eucalyptus globulus* leaf litter in Californian streams. *Freshwater Biology*, *55*(4), 739–752. https://doi.org/10.1111/j.1365-2427.2009.02312.x
Lamberti, G. A., & Berg, M. B. (1995). Invertebrates and other benthic features as indicators of environmental change in Juday Creek, Indiana. *Natural Areas Journal*, *15*(3), 249–258.
Larrañaga, A., Perkins, D. M., Basaguren, A., Larrañaga, S., Pozo, J., & Montoya, J. M. (2023). Land use drives detritivore size structure and decomposition through shifts in resource quality and quantity. *Science of the Total Environment*, *892*, 164552. https://doi.org/10.1016/j.scitotenv.2023.164552
Lecerf, A., Baudoin, J.-M., Besson, A. A., Lamothe, S., & Lagrue, C. (2012). Is smaller necessarily better? Effects of small-scale forest harvesting on stream ecosystems. *Annales de Limnologie*, *48*(4), 401–409. https://doi.org/10.1051/limn/2012028
Lecerf, A., & Chauvet, E. (2008). Diversity and functions of leaf-decaying fungi in human-altered streams. *Freshwater Biology*, *53*(8), 1658–1672. https://doi.org/10.1111/j.1365-2427.2008.01986.x
Lecerf, A., & Richardson, J. S. (2010). Litter decomposition can detect effects of high and moderate levels of forest disturbance on stream condition. *Forest Ecology and Management*, *259*(12), 2433–2443. https://doi.org/10.1016/j.foreco.2010.03.022
Lemes da Silva, A. L., Lemes, W. P., Andriotti, J., Petrucio, M. M., & Feio, M. J. (2020). Recent land-use changes affect stream ecosystem processes in a subtropical island in Brazil. *Austral Ecology*, *45*(5), 644–658. https://doi.org/10.1111/aec.12879
Link, M., Schreiner, V. C., Graf, N., Szöcs, E., Bundschuh, M., Battes, K. P., Cîmpean, M., Sures, B., Grabner, D., Buse, J., & Schäfer, R. B. (2022). Pesticide effects on macroinvertebrates and leaf litter decomposition in areas with traditional agriculture. *Science of the Total Environment*, *828*, 154549. https://doi.org/10.1016/j.scitotenv.2022.154549
Lopes, M. P., Martins, R. T., Silveira, L. S., & Alves, R. G. (2015). The leaf breakdown of *Picramnia sellowii* (Picramniales: Picramniaceae) as index of anthropic disturbances in tropical streams. *Brazilian Journal of Biology*, *75*, 846–853. https://doi.org/10.1590/1519-6984.00414
Lorion, C. M., & Kennedy, B. P. (2009). Riparian forest buffers mitigate the effects of deforestation on fish assemblages in tropical headwater streams. *Ecological Applications*, *19*(2), 468–479. https://doi.org/10.1890/08-0050.1

Lubanga, H. L., Manyala, J. O., Sitati, A., Yegon, M. J., & Masese, F. O. (2021). Spatial variability in water quality and macroinvertebrate assemblages across a disturbance gradient in the Mara River Basin, Kenya. *Ecohydrology & Hydrobiology*, *21*(21), 718–730. https://doi.org/10.1016/j.ecohyd.2021.03.001

Mancuso, J., Tank, J. L., Mahl, U. H., Vincent, A., & Tiegs, S. D. (2023). Monthly variation in organic-matter decomposition in agricultural stream and riparian ecosystems. *Aquatic Sciences*, *85*(3), 83. https://doi.org/10.1007/s00027-023-00975-7

Maridet, L., Wasson, J.-G., Philippe, M., Amoros, C., & Naiman, R. (1998). Trophic structure of three streams with contrasting riparian vegetation and geomorphology. *Archiv für Hydrobiologie*, 61–85. https://doi.org/10.1127/archiv-hydrobiol/144/1998/61

Márquez, J. A., Principe, R. E., Cibils Martina, L., & Albariño, R. J. (2017). Pine needle litter acts as habitat but not as food source for stream invertebrates. *International Review of Hydrobiology*, *102*(1–2), 29–37. https://doi.org/10.1002/iroh.201601856

Martínez, A., Larrañaga, A., Pérez, J., Descals, E., Basaguren, A., & Pozo, J. (2013). Effects of pine plantations on structural and functional attributes of forested streams. *Forest Ecology and Management*, *310*, 147–155. https://doi.org/10.1016/j.foreco.2013.08.024

Masese, F. O., Kitaka, N., Kipkemboi, J., Gettel, G. M., Irvine, K., & McClain, M. E. (2014). Litter processing and shredder distribution as indicators of riparian and catchment influences on ecological health of tropical streams. *Ecological Indicators*, *46*, 23–37. https://doi.org/10.1016/j.ecolind.2014.05.032

Mckie, B. G., & Malmqvist, B. (2009). Assessing ecosystem functioning in streams affected by forest management: Increased leaf decomposition occurs without changes to the composition of benthic assemblages. *Freshwater Biology*, *54*(10), 2086–2100. https://doi.org/10.1111/j.1365-2427.2008.02150.x

McNeish, R. E., Benbow, M. E., & McEwan, R. W. (2017). Removal of the invasive shrub, *Lonicera maackii* (Amur Honeysuckle), from a headwater stream riparian zone shifts taxonomic and functional composition of the aquatic biota. *Invasive Plant Science and Management*, *10*(3), 232–246. https://doi.org/10.1017/inp.2017.22

McNeish, R. E., Moore, E. M., Benbow, M. E., & McEwan, R. W. (2015). Removal of the invasive shrub, *Lonicera maackii* , from riparian forests influences headwater stream biota and ecosystem function. *River Research and Applications*, *31*(9), 1131–1139. https://doi.org/10.1002/rra.2808

Mctammany, M. E., Benfield, E. F., & Webster, J. R. (2008). Effects of agriculture on wood breakdown and microbial biofilm respiration in southern Appalachian streams. *Freshwater Biology*, *53*(4), 842–854. https://doi.org/10.1111/j.1365-2427.2007.01936.x

Menéndez, M., Descals, E., Riera, T., & Moya, O. (2013). Do non-native *Platanus hybrida* riparian plantations affect leaf litter decomposition in streams? *Hydrobiologia*, *716*(1), 5–20. https://doi.org/10.1007/s10750-013-1539-0

Menninger, H. L., & Palmer, M. A. (2007). Herbs and grasses as an allochthonous resource in open-canopy headwater streams. *Freshwater Biology*, *52*(9), 1689–1699. https://doi.org/10.1111/j.1365-2427.2007.01797.x

Mesa, L. M., Reynaga, M. C., Correa, M. del V., & Sirombra, M. G. (2013). Effects of anthropogenic impacts on benthic macroinvertebrates assemblages in subtropical mountain streams. *Iheringia. Série Zoologia*, *103*, 342–349. https://doi.org/10.1590/S0073-47212013000400002

Mlambo, M. C., Paavola, R., Fritze, H., Louhi, P., & Muotka, T. (2019). Leaf litter decomposition and decomposer communities in streams affected by intensive forest biomass removal. *Ecological Indicators*, *101*, 364–372. https://doi.org/10.1016/j.ecolind.2019.01.035

Molinero, J., Pozo, J., & Gonzalez, E. (1996). Litter breakdown in streams of the Agüera catchment: Influence of dissolved nutrients and land use. *Freshwater Biology*, *36*(3), 745–756. https://doi.org/10.1046/j.1365-2427.1996.00125.x

Monroy, S., Martínez, A., López-Rojo, N., Pérez-Calpe, A. V., Basaguren, A., & Pozo, J. (2017). Structural and functional recovery of macroinvertebrate communities and leaf litter decomposition after a marked drought: Does vegetation type matter? *Science of the Total Environment*, *599–600*, 1241–1250. https://doi.org/10.1016/j.scitotenv.2017.05.093

Morabowen, A., Crespo-Pérez, V., & Ríos-Touma, B. (2019). Effects of agricultural landscapes and land uses in highly biodiverse tropical streams of the Ecuadorian Choco. *Inland Waters*, *9*(3), 289–300. https://doi.org/10.1080/20442041.2018.1527597

Murphy, J. F., & Giller, P. S. (2001). Hazel leaf breakdown in two low-order streams differing in the functional efficiency of their detritivore assemblages. *Archiv für Hydrobiologie*, 249–267. https://doi.org/10.1127/archiv-hydrobiol/150/2001/249

Musetta-Lambert, J., Muto, E., Kreutzweiser, D., & Sibley, P. (2017). Wildfire in boreal forest catchments influences leaf litter subsidies and consumer communities in streams: Implications for riparian management strategies. *Forest Ecology and Management*, *391*, 29–41. https://doi.org/10.1016/j.foreco.2017.01.028

Myers, L., Mihuc, T., & Woodcock, T. (2007). The impacts of forest management on invertebrate communities associated with submerged leaves in forested Adirondack streams. *Journal of Freshwater Ecology*, *22*(2), 325–331. https://doi.org/10.1080/02705060.2007.9665054

Oelbermann, M., & Gordon, A. M. (2000). Quantity and quality of autumnal litterfall into a rehabilitated agricultural stream. *Journal of Environmental Quality*, *29*(2), 603–611. https://doi.org/10.2134/jeq2000.00472425002900020031x

Oester, R., dos Reis Oliveira, P. C., Moretti, M. S., Altermatt, F., & Bruder, A. (2023). Leaf-associated macroinvertebrate assemblage and leaf litter breakdown in headwater streams depend on local riparian vegetation. *Hydrobiologia*, *850*(15), 3359–3374. https://doi.org/10.1007/s10750-022-05049-7

Ono, E. R., Manoel, P. S., Melo, A. L. U., & Uieda, V. S. (2020). Effects of riparian vegetation removal on the functional feeding group structure of benthic macroinvertebrate assemblages. *Community Ecology*, *21*(2), 145–157. https://doi.org/10.1007/s42974-020-00014-7

Paul, M. J., Meyer, J. L., & Couch, C. A. (2006). Leaf breakdown in streams differing in catchment land use. *Freshwater Biology*, *51*(9), 1684–1695. https://doi.org/10.1111/j.1365-2427.2006.01612.x

Principe, R. E., Márquez, J. A., Martina, L. C., Jobbágy, E. G., & Albariño, R. J. (2015). Pine afforestation changes more strongly community structure than ecosystem functioning in grassland mountain streams. *Ecological Indicators*, *57*, 366–375. https://doi.org/10.1016/j.ecolind.2015.04.033

Reid, D. J., Lake, P. S., & Quinn, G. P. (2013). Influences of agricultural landuse and seasonal changes in abiotic conditions on invertebrate colonisation of riparian leaf detritus in intermittent streams. *Aquatic Sciences*, *75*(2), 285–297. https://doi.org/10.1007/s00027-012-0273-4

Rezende, R. de S., Cararo, E. R., Bernardi, J. P., Chimello, V., Lima-Rezende, C. A., Albeny-Simões, D., Dal Magro, J., & Gonçalves, J. F. (2021). Land cover affects the breakdown of *Pinus elliottii* needles litter by microorganisms in soil and stream systems of subtropical riparian zones. *Limnologica*, *90*, 125905. https://doi.org/10.1016/j.limno.2021.125905

Riipinen, M. P., Fleituch, T., Hladyz, S., Woodward, G., Giller, P., & Dobson, M. (2009). Invertebrate community structure and ecosystem functioning in European conifer plantation streams. *Freshwater Biology*, *55*, 346–359. https://doi.org/10.1111/j.1365-2427.2009.02278.x

Roberts, M. L., & Bilby, R. E. (2009). Urbanization alters litterfall rates and nutrient inputs to small Puget Lowland streams. *Journal of the North American Benthological Society*, *28*(4), 941–954. https://doi.org/10.1899/07-160.1

Rossi, F., Mallet, C., Portelli, C., Donnadieu, F., Bonnemoy, F., & Artigas, J. (2019). Stimulation or inhibition: Leaf microbial decomposition in streams subjected to complex chemical contamination. *Science of the Total Environment*, *648*, 1371–1383. https://doi.org/10.1016/j.scitotenv.2018.08.197

Rubio-Ríos, J., Salinas-Bonillo, M. J., Pérez, J., Fenoy, E., Boyero, L., & Casas, J. J. (2023). Alder stands promote N-cycling but not leaf litter mass loss in Mediterranean streams flowing through pine plantations. *Forest Ecology and Management*, *542*, 121072. https://doi.org/10.1016/j.foreco.2023.121072

Ryan, D. K., & Kelly-Quinn, M. (2016). Riparian vegetation management for water temperature regulation: Implications for the production of macroinvertebrate prey of salmonids. *Fisheries Management and Ecology*, *23*(6), 519–530. https://doi.org/10.1111/fme.12193

Ryder, L., de Eyto, E., Gormally, M., Skeffington, M. S., Dillane, M., & Poole, R. (2011). Riparian zone creation in established coniferous forests in Irish upland peat catchments: Physical, chemical and biological Implications. *Biology and Environment: Proceedings of the Royal Irish Academy*, *111B*(1), 41–60.

Sakai, M., Natuhara, Y., Fukushima, K., Imanishi, A., Imai, K., & Kato, M. (2013). Ecological functions of persistent Japanese cedar litter in structuring stream macroinvertebrate assemblages. *Journal of Forest Research*, *18*(2), 190–199. https://doi.org/10.1007/s10310-012-0339-0

Sarriquet, P. E., Delettre, Y. R., & Marmonier, P. (2006). Effects of catchment disturbance on stream invertebrates: Comparison of different habitats (vegetation, benthic and interstitial) using bio-ecological groups. *Annales de Limnologie - International Journal of Limnology*, *42*(4), 205–219. https://doi.org/10.1051/limn/2006022

Shah, N. W., Nisbet, T. R., & Broadmeadow, S. B. (2021). The impacts of conifer afforestation and climate on water quality and freshwater ecology in a sensitive peaty catchment: A 25 year study in the upper River Halladale in North Scotland. *Forest Ecology and Management*, *502*, 119616. https://doi.org/10.1016/j.foreco.2021.119616

Silva-Araújo, M., Silva-Junior, E. F., Neres-Lima, V., Feijó-Lima, R., Tromboni, F., Lourenço-Amorim, C., Thomas, S. A., Moulton, T. P., & Zandonà, E. (2020). Effects of riparian deforestation on benthic invertebrate community and leaf processing in Atlantic forest streams. *Perspectives in Ecology and Conservation*, *18*(4), 277–282. https://doi.org/10.1016/j.pecon.2020.09.004

Suga, C. M., & Tanaka, M. O. (2013). Influence of a forest remnant on macroinvertebrate communities in a degraded tropical stream. *Hydrobiologia*, *703*(1), 203–213. https://doi.org/10.1007/s10750-012-1360-1

Tanaka, M. O., Fernandes, J. de F., Suga, C. M., Hanai, F. Y., & Souza, A. L. T. de. (2015). Abrupt change of a stream ecosystem function along a sugarcane-forest transition: Integrating riparian and in-stream characteristics. *Agriculture, Ecosystems & Environment*, *207*, 171–177. https://doi.org/10.1016/j.agee.2015.04.014

Tolkkinen, M., Mykrä, H., Annala, M., Markkola, A. M., Vuori, K. M., & Muotka, T. (2015). Multi-stressor impacts on fungal diversity and ecosystem functions in streams: Natural vs. anthropogenic stress. *Ecology*, *96*(3), 672–683. https://doi.org/10.1890/14-0743.1

Tonello, G., Decian, V. S., Restello, R. M., & Hepp, L. U. (2021). The conversion of natural riparian forests into agricultural land affects ecological processes in Atlantic forest streams. *Limnologica*, *91*, 125927. https://doi.org/10.1016/j.limno.2021.125927

Torres, P. J., & Ramírez, A. (2014). Land use effects on leaf litter breakdown in low-order streams draining a rapidly developing tropical watershed in Puerto Rico. *Revista de Biología Tropical*, *62*(S2), Article S2. https://doi.org/10.15517/rbt.v62i0.15783

Townsend, C. R., Downes, B. J., Peacock, K., & Arbuckle, C. J. (2004). Scale and the detection of land-use effects on morphology, vegetation and macroinvertebrate communities of grassland streams. *Freshwater Biology*, *49*(4), 448–462. https://doi.org/10.1111/j.1365-2427.2004.01192.x

Truchy, A., Sponseller, R. A., Ecke, F., Angeler, D. G., Kahlert, M., Bundschuh, M., Johnson, R. K., & McKie, B. G. (2022). Responses of multiple structural and functional indicators along three contrasting disturbance gradients. *Ecological Indicators*, *135*, 108514. https://doi.org/10.1016/j.ecolind.2021.108514

Tupinambás, T. H., Callisto, M., & Santos, G. B. (2007). Benthic macroinvertebrate assemblages structure in two headwater streams, south-eastern Brazil. *Revista Brasileira de Zoologia*, *24*(4), 887–897. https://doi.org/10.1590/S0101-81752007000400005

Turunen, J., Aroviita, J., Marttila, H., Louhi, P., Laamanen, T., Tolkkinen, M., Luhta, P.-L., Kløve, B., & Muotka, T. (2017). Differential responses by stream and riparian biodiversity to in-stream restoration of forestry-impacted streams. *Journal of Applied Ecology*, *54*(5), 1505–1514. https://doi.org/10.1111/1365-2664.12897

Valente-Neto, F., Koroiva, R., Fonseca-Gessner, A. A., & Roque, F. de O. (2015). The effect of riparian deforestation on macroinvertebrates associated with submerged woody debris. *Aquatic Ecology*, *49*(1), 115–125. https://doi.org/10.1007/s10452-015-9510-y

Walsh, C. J., Gooderham, J. P. R., Grace, M. R., Sdraulig, S., Rosyidi, M. I., & Lelono, A. (2002). The relative influence of diffuse- and point-source disturbances on a small upland stream in East Java, Indonesia: A preliminary investigation. *Hydrobiologia*, *487*(1), 183–192. https://doi.org/10.1023/A:1022990822198

Whiles, M. R., & Wallace, J. B. (1997). Leaf litter decomposition and macroinvertebrate communities in headwater streams draining pine and hardwood catchments. *Hydrobiologia*, *353*(1), 107–119. https://doi.org/10.1023/A:1003054827248

Yeung, A. C. Y., Lecerf, A., & Richardson, J. S. (2017). Assessing the long-term ecological effects of riparian management practices on headwater streams in a coastal temperate rainforest. *Forest Ecology and Management*, *384*, 100–109. https://doi.org/10.1016/j.foreco.2016.10.044

Youngquist, M. B., Wiley, C., Eggert, S. L., D'Amato, A. W., Palik, B. J., & Slesak, R. A. (2020). Foundation species loss affects leaf breakdown and aquatic invertebrate resource use in black ash wetlands. *Wetlands*, *40*(4), 839–852. https://doi.org/10.1007/s13157-019-01221-3

Zhang, Y., Richardson, J. S., & Pinto, X. (2009). Catchment-scale effects of forestry practices on benthic invertebrate communities in Pacific coastal streams. *Journal of Applied Ecology*, *46*(6), 1292–1303. https://doi.org/10.1111/j.1365-2664.2009.01718.x

Zúñiga-Sarango, W., Gaona, F. P., Reyes-Castillo, V., & Iñiguez-Armijos, C. (2020). Disrupting the biodiversity–ecosystem function relationship: Response of shredders and leaf breakdown to urbanization in Andean streams. *Frontiers in Ecology and Evolution*, *8*. https://doi.org/10.3389/fevo.2020.592404
